# Microbial profiling and single-cell transcriptomics reveal probiotic mechanisms of coral thermal resilience

**DOI:** 10.1101/2025.09.02.673423

**Authors:** Chih-Ying Lu, Yung-Pei Chang, Victor M. Piñon-Gonzalez, Thomas D. Lewin, Kai-Ning Shih, Po-Shun Chuang, Naohisa Wada, Sheng-Ping Yu, Zhao-Rong Yu, Yi-Hua Chen, Mo Chen, Ching-Hsiang Chin, Yu-Jing Chiou, Yi-Ling Chiu, Isabel Jiah-Yih Liao, Hsin-Feng Chang, Jui-Hung Yen, Mei-Yeh Jade Lu, Yi-Jyun Luo, Sen-Lin Tang

## Abstract

Probiotics hold promise for enhancing coral resilience under climate-driven thermal stress, yet their mechanisms remain poorly understood. Here, we evaluate two *Endozoicomonas* species as coral probiotics and characterize their effects on microbial communities and host gene expression. We show that *E. acroporae* Acr-14^T^ enhances thermal tolerance in *Stylophora pistillata*, suppresses opportunistic pathogens, and promotes beneficial microbes. To facilitate transcriptomic profiling, we assembled a chromosome-level genome of *S. pistillata* clade 1 (Pacific lineage) and used it to reveal that *E. acroporae* Acr-14^T^ mitigates heat-induced protein-folding stress and supports host energy homeostasis. Single-cell transcriptomics further uncovered enhanced pro-survival signaling and modulation of the *S*-adenosylmethionine (SAMe) synthesis pathway. Together, our findings identify *E. acroporae* Acr-14^T^ as a robust coral probiotic and provide mechanistic insights into host–microbe interactions that promote coral resilience under thermal stress.

## INTRODUCTION

Coral reefs are among the most biodiverse marine ecosystems on earth, supporting over a million marine species^1,2^ and providing ecosystem services essential to coastal economies.^3^ However, more than half of global coral coverage has been lost due to anthropogenic stressors.^4^ Projections suggest that if warming exceeds 1.5°C above pre-industrial levels, a threshold that already been surpassed by 2020,^5^ up to 90% of tropical coral reefs could vanish at the end of this century.^6,7^ The recent fourth global coral bleaching event reinforces the urgency of identifying interventions that can bolster coral resilience under thermal stress.^8,9^ Coral probiotics represent a promising strategy for enhancing coral health.^10^ The coral probiotic hypothesis proposes that introducing beneficial microbes into reefs can bolster coral resilience,^11^ potentially through microbiome restructuring,^12^ nutrient cycling,^13^ and the production of antioxidant^14^ or antimicrobial compounds.^15^ These beneficial microorganisms for corals (BMC)^16^ have been shown to mitigate bleaching^17,18,19^ and promote recovery from heat stress, including via mechanisms such as DMSP degradation.^20^ Evidence suggests that these beneficial effects are likely multifactorial, arising from direct host interactions, indirect microbial functions, and nutritional supplementation.^21,22,23^ Despite these advances, the full spectrum of mechanisms underlying probiotic benefits remains largely unresolved, and progress toward this understanding is hampered by the inherent complexity of coral holobionts,^24,25^ the scarcity of well-annotated coral genomes, and the underutilization of high-resolution genomic tools such as single-cell transcriptomics.^26^

*Endozoicomonas* is a widespread and abundant bacterial genus associated with healthy corals, with genomic evidence indicating that multiple species possess beneficial metabolic capacities and mutualistic roles.^27^ However, this highly diverse genus may also include members with parasitic or pathogenic lifestyles.^28^ Our previous works demonstrated that two species, *E. acroporae* Acr-14^T^ and *E. montiporae* CL-33^T^, both isolated from Kenting, Taiwan, possess functions potentially advantageous to corals. Specifically, *E. acroporae* Acr-14^T^ exhibits potent extracellular reactive oxygen species (ROS)-scavenging activity^29^ and metabolizes dimethylsulfoniopropionate (DMSP) via the cleavage pathway, influencing coral sulfur cycling.^30^ *E. montiporae* CL-33^T^ carries isocitrate lyase, which may promote host gluconeogenesis through a type III secretion system, as well as ephrin ligands, which may facilitate symbiosis.^31^ Despite these promising traits, the probiotic potential of these two *Endozoicomonas* strains remains untested.

Here, we evaluated the probiotic efficacy of both species in *Stylophora pistillata*, a prevalent coral host in which *E. acroporae* Acr-14^T^ has been detected.^32,33^ We found that *E. acroporae* Acr-14^T^ enhanced *S. pistillata* thermal tolerance, colonized host tissues, suppressed opportunistic bacteria, and promoted beneficial microbes. To uncover the molecular basis of these effects, we generated a chromosome-level genome of *S. pistillata* clade 1 and integrated bulk and single-cell transcriptomic analyses. This approach revealed cell-type-specific coral heat stress responses and showed that *E. acroporae* Acr-14^T^ enhances host thermal tolerance by alleviating the protein-folding burden and modulating SAMe metabolism. Together, these findings position *E. acroporae* Acr-14^T^ as a promising coral probiotic and reveal a mechanistic basis for microbe-mediated enhancement of host thermal resilience.

## RESULTS

We conducted three sequential coral bleaching experiments to evaluate, validate, and investigate the mechanisms of probiotic function (Figure 1A and S1A-S1C). In the efficacy evaluation phase, two candidate probiotics were tested in *S. pistillata* from Mao’ao, northeastern Taiwan. The effective strain was then validated in a geographically distinct *S. pistillata* population from Kenting, southern Taiwan, and subsequently examined through the molecular profiling experiment. In the first two experiments, we monitored bleaching responses and microbial community dynamics, whereas in the third experiment, we additionally localized the probiotic within coral tissues, assessed oxidative stress, constructed a *S. pistillata* clade 1 genome, and profiled transcriptomic responses at both bulk and single-cell resolution.

**Figure 1.**
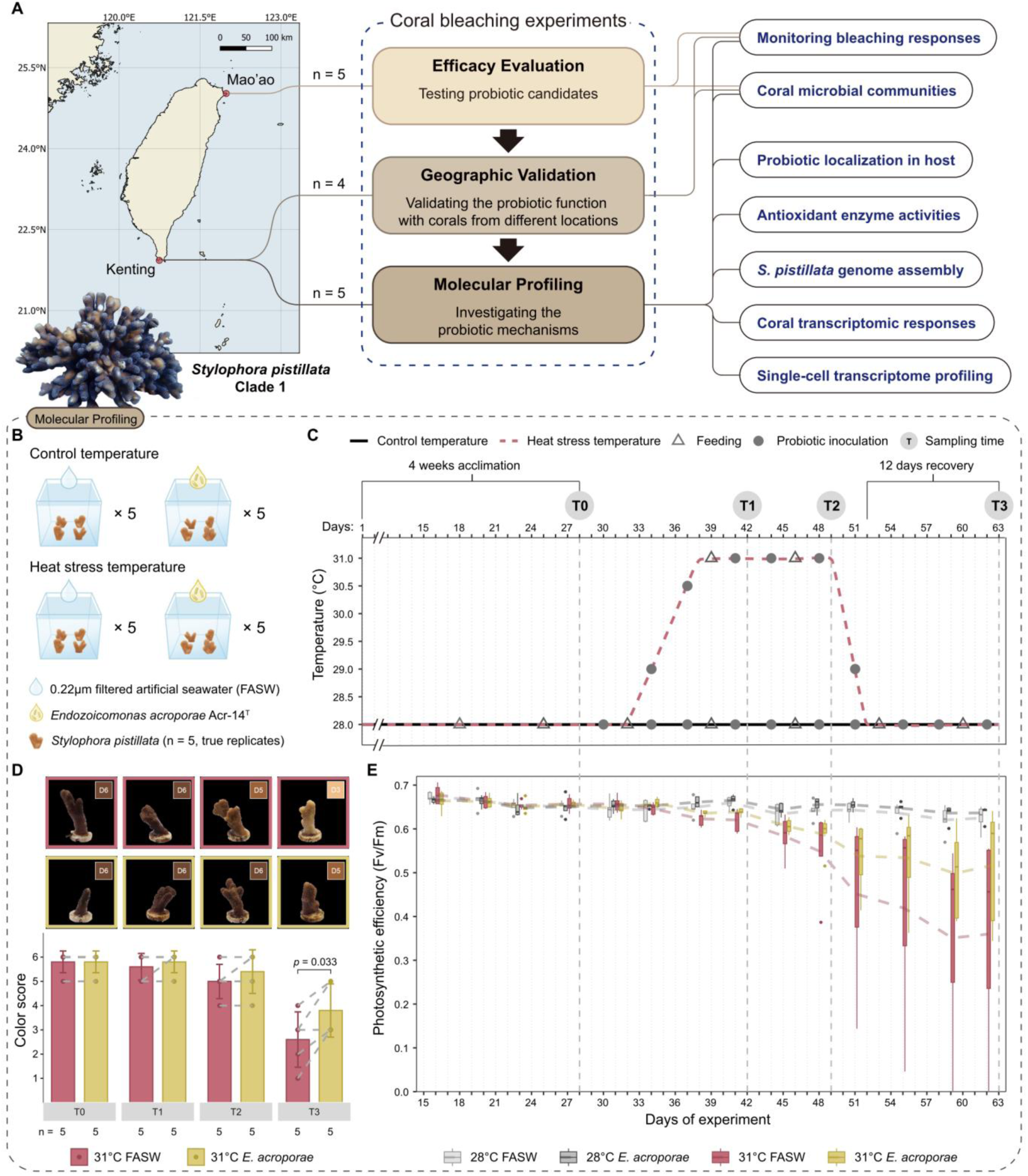
Experimental workflow and probiotic effects on coral bleaching responses. (A) Sampling map and overall research workflow. (B) Treatment and replication design for the molecular profiling experiment. Dashed line group figures corresponding to the same experiment. Each treatment comprised five independent aquarium systems. Each aquarium contained four coral nubbins assigned to specific sampling time points. During each sampling event, five nubbins per treatment were collected, each originating from different coral colonies (n = 5, true replication). (C) Detailed treatment regimens, including temperature settings, and feeding dates, probiotic inoculation, and sample collection. (D) Coral color scores of nubbins collected at the four sampling time points, with photos of nubbins (colony c3). Grey dashed lines connect same-colony data across treatments. Dots represent data point. Data are represented as means ± SD (n = 5; two-sided paired *t*-test). (E) Photosynthetic efficiency (Fv/Fm) of nubbins. In boxplots, dots represent outliers, and dashed lines connect treatment means (n = 5; One-way ANOVA with Tukey post-hoc test). See Figures S1 and S2.

### *E. acroporae* Acr-14^T^ consistently enhances coral thermal tolerance across three independent experiments

In efficacy evaluation, both *E. acroporae* Acr-14^T^ and *E. montiporae* CL-33^T^ were tested using *S. pistillata* from Mao’ao (Figures S1C, S2A and S2C). From day 17 onward, maximum quantum yield (Fv/Fm) measurements revealed that corals treated with *E. acroporae* Acr-14^T^ maintained significantly higher photosynthetic efficiency under heat stress compared to both the 31°C FASW (filtered artificial seawater) and 31°C *E. montiporae* treatments (adjusted *p* < 0.05; Figure S2E). No significant difference was observed between the 31°C FASW and 31°C *E. montiporae* groups (adjusted *p* > 0.05).

To validate these results, we repeated the experiment using *S. pistillata* from Kenting. In this second trial (geographic validation), we included a heat-sterilized *E. acroporae* Acr-14^T^ treatment to examine potential postbiotic^34^ effects (Figures S1C, S2B and S2D). Again, corals treated with live *E. acroporae* Acr-14^T^ maintained higher photosynthetic efficiency under heat stress, showing no statistical differences from the ambient-temperature groups. In contrast, photosynthetic efficiency in the 31°C FASW treatment was significantly lower than the ambient controls by day 24 (adjusted *p* < 0.05; Figure S2F). The heat-sterilized treatment displayed an intermediate phenotype, indistinguishable from either the 31°C FASW or 31°C *E. acroporae* groups (adjusted *p* > 0.05).

In molecular profiling, we incorporated true replication and multiple sampling time points to monitor physiological changes chronologically (Figures 1B, 1C, and S1C). The four sampling times represent post-acclimation (T0), during heat stress (T1), the peak of heat stress (T2), and after a recovery period (T3). Throughout this experiment, photosynthetic efficiency in 31°C *E. acroporae*-treated corals remained comparable to that of ambient-temperature controls (28°C FASW or 28°C *E. acroporae*, adjusted *p* > 0.05), whereas the 31°C FASW group exhibited a significant decline after day 38 (adjusted *p* < 0.05; Figure 1E). This decline was accompanied by visible bleaching and lower color scores compared to the probiotic-treated group at T3 (*p* < 0.05; Figure 1D).

Coral colonies vary in thermal tolerance,^35^ as reflected in our photosynthetic efficiency measurements. Mao’ao and Kenting differ in oceanographic conditions, shaping distinct reef communities and thermal baselines.^36^ Nevertheless, across all tested colonies, *E. acroporae* Acr-14^T^ consistently improved photosynthetic efficiency under heat stress, indicating that the probiotic effect was robust across genotypes. These findings prompted us to investigate how *E. acroporae* Acr-14^T^ interacts with the coral holobiont to mediate thermal protection.

### *E. acroporae* Acr-14^T^ colonizes host tissue and forms microbial aggregates

We investigated whether *E. acroporae* Acr-14^T^ actively colonizes coral hosts and contributes to a stable microbial association. Microbiome profiling using 16S rRNA gene sequencing revealed that coral microbial communities were dominated by *Alphaproteobacteria*, *Gammaproteobacteria*, *Desulfovibrionia*, and *Campylobacteria* (Figures S3A-S3C). In all three experiments, *E. acroporae* Acr-14^T^ was consistently detected in corals receiving the probiotic treatment (Figures 2A-2C, and S3D; Table S1A), while *E. montiporae* CL-33^T^ was scarcely detected, averaging only 0.05% of the relative abundance of treated corals (Figure 2A). Probiotic untreated corals, including baseline samples (T0), contained either no or few (< 0.5%) *Endozoicomonas* sequences (Figures 2A-2C). Moreover, the relative abundance of *E. acroporae* Acr-14^T^ was substantially higher in corals than in tank water, indicating that its detection reflected active host colonization rather than environmental carryover (Figure 2C).

**Figure 2.**
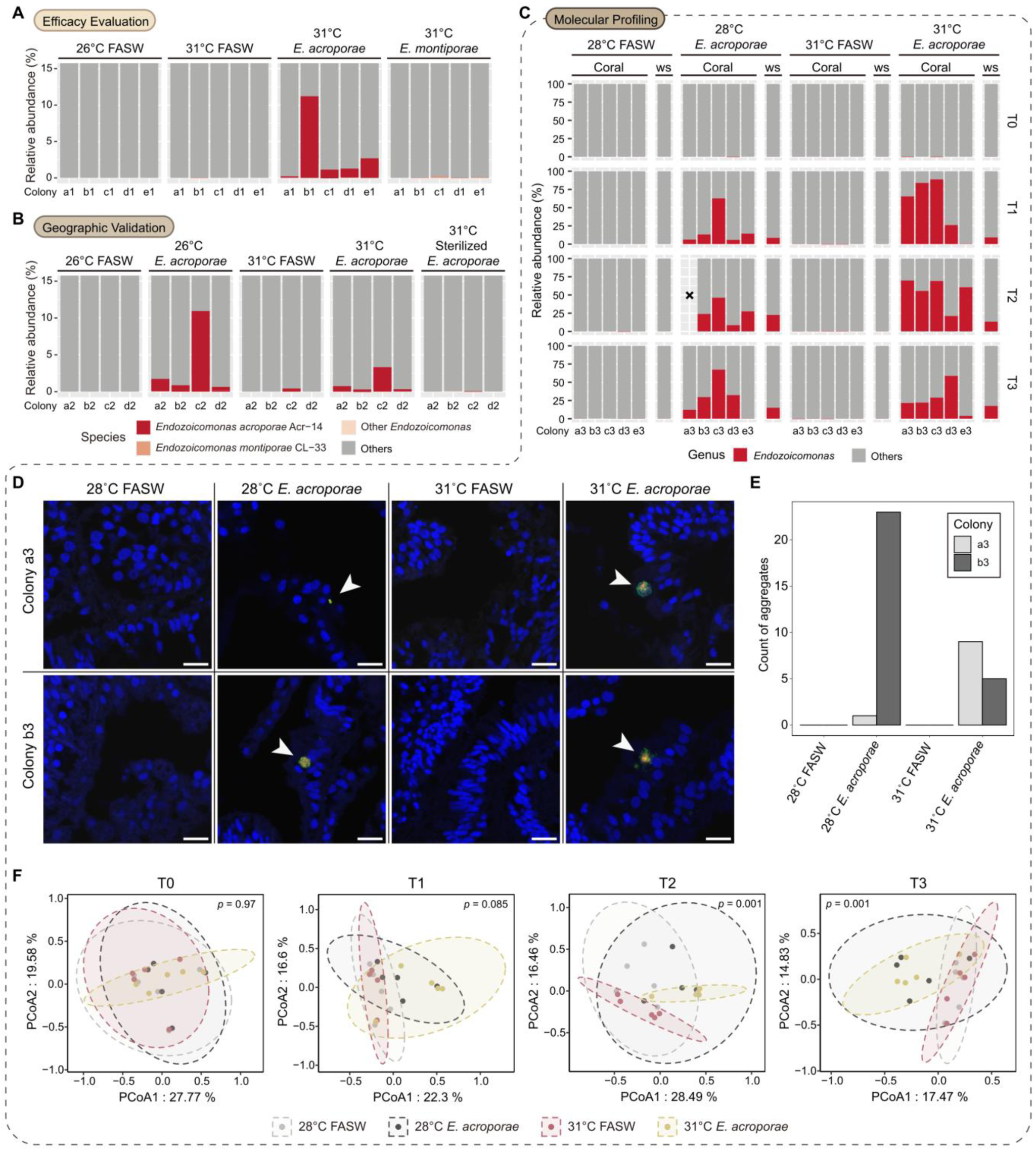
*E. acroporae* Acr-14^T^ colonizes host tissue, forms CAMAs, and reshapes coral microbial communities. (A-C) Relative abundance of *Endozoicomonas* (species-level assignment see STAR Methods and Table S1A) in samples from the efficacy evaluation (A), geographic validation (B), and molecular profiling experiments (C) via short-length 16S rRNA gene sequencing. Dashed line group figures corresponding to the same experiment. Each bar represents one coral colony, with colony codes indicated on the x-axis. Tank water samples (ws); lost sample (x). (D) Confocal images showing merged signals from FISH using the EUB338mix probe^38^ (red, all bacteria), the Endo-Group B probe^39^ (green, *Endozoicomonas-*specific), Non338 probe^40^ (white, negative control) and DAPI (nuclei and host autofluorescence), localizing *Endozoicomonas* (white arrows) in coral tissue at T3. Scale bars: 10 µm. (E) Numbers of *Endozoicomonas* aggregates among the four treatments at T3 (two samples per treatment, five slides per sample, 40 slides in total). (F) Principal coordinates analysis (PCoA) demonstrating treatment effects on coral microbial composition at each time point, with 95% confidence ellipses (PERMANOVA; pairwise comparison *p* values adjusted using Bonferroni correction). See Figures S3 and S4 and Table S1.

To determine whether *E. acroporae* Acr-14^T^ physically associates with the coral host, we performed fluorescence *in situ* hybridization (FISH) on samples collected at T3 of molecular profiling. *Endozoicomonas*-specific signals were detected exclusively in corals treated with *E. acroporae* Acr-14^T^ (Figures 2D, and S4A-S4C). These bacteria formed coral-associated microbial aggregates (CAMAs) in the tissue, the number and size of which correlated with the relative abundance of *E. acroporae* Acr-14^T^ (Figures 2C, 2E, and S4C). Most signals localized to the mesenterial filaments (Figure S4A), with fewer in the tentacles, similar to observations in wild corals.^37^ These findings demonstrate that probiotic treatment establishes a direct symbiotic association between *S. pistillata* and *E. acroporae* Acr-14^T^.

### Heat stress and probiotic treatments significantly reshape coral microbial composition

We next analyzed microbial composition to determine whether probiotic treatments reshape the microbial community and whether these changes correlate with thermal resistance. Because the three coral bleaching experiments involved corals collected from different locations and seasons, the microbial communities differed significantly among experiments (Figure S5A). To minimize batch effects, all analyses were performed independently within each experiment, without cross-experiment pooling. Probiotic treatment had a limited impact on microbial alpha diversity, with a significant increase in amplicon sequence variant (ASV) richness observed only in efficacy evaluation (Figures S3E-S3G). In contrast, significant differences in microbial composition between treatments were observed in all three experiments (Figures S5B, S5C, and 2F), suggesting that specific community structures reflected responses to probiotic or heat stress treatments.

Molecular profiling revealed clear temporal dynamics in microbial composition (Figure 2F). At baseline (T0), no significant differences were detected among treatments. Following treatment initiation (T1), corals treated with *E. acroporae* Acr-14^T^ exhibited a modest, non-significant shift in community structure compared to FASW controls. By peak heat stress (T2), significant compositional differences emerged specifically between the 31°C FASW and 31°C *E. acroporae* treatments (adjusted *p* = 0.03), while after recovery (T3), differences were particularly observed between the 28°C FASW and 31°C *E. acroporae* groups (adjusted *p* = 0.048). Notably, in efficacy evaluation, both *E. acroporae* Acr-14^T^ and *E. montiporae* CL-33^T^ treatments significantly shifted microbial communities compared to the 31°C FASW group (adjusted *p* < 0.05; Figure S5B). However, only *E. acroporae* Acr-14^T^ conferred measurable thermal protection, indicating that microbial restructuring alone is insufficient to explain the probiotic effect.

### Probiotic treatment suppresses pathogens and promotes beneficial microbes

Given that probiotic treatment reshapes coral microbial communities, we next sought to identify microbial taxa associated with *E. acroporae* Acr-14^T^. We performed genus-level differential abundance analyses using ANCOM-BC and MaAsLin2, adopting a consensus approach (STAR Methods). To distinguish microbial signatures associated with probiotic-induced resilience, we focused on genera uniquely associated with either the 31°C FASW or 31°C *E. acroporae* treatments (Figures 3A-3C). A consistent pattern emerged across the three experiments: *E. acroporae* Acr-14^T^ suppressed potential opportunistic pathogens and promoted beneficial microbes. In efficacy evaluation, *Desulfovibrio* and *Alteromonas*, known opportunists^41,42^, were enriched in the 31°C FASW group, but not in *E. acroporae* Acr-14^T^-treated corals (Figure 3A; Table S2A). Similarly, in molecular profiling, *Legionellaceae*^43^ (genus unassigned) was enriched at T2, and *Halodesulfovibrio*^44^ and *Peptostreptococcales-Tissierellales*^45^ (genus unassigned) were enriched at T3 in the 31°C FASW treatment (Figure 3C; Table S2E and S2F). Notably, this pathogen enrichment was absent under mild stress (T1; Degree Heating Weeks (DHW)^46^ = 1.9°C-weeks; Figure 3C; Table S2D), and in the geographic validation experiment (DHW = 2°C-weeks) (Figure 3B; Table S2B), suggesting that opportunists only proliferate under more severe thermal stress conditions.

**Figure 3.**
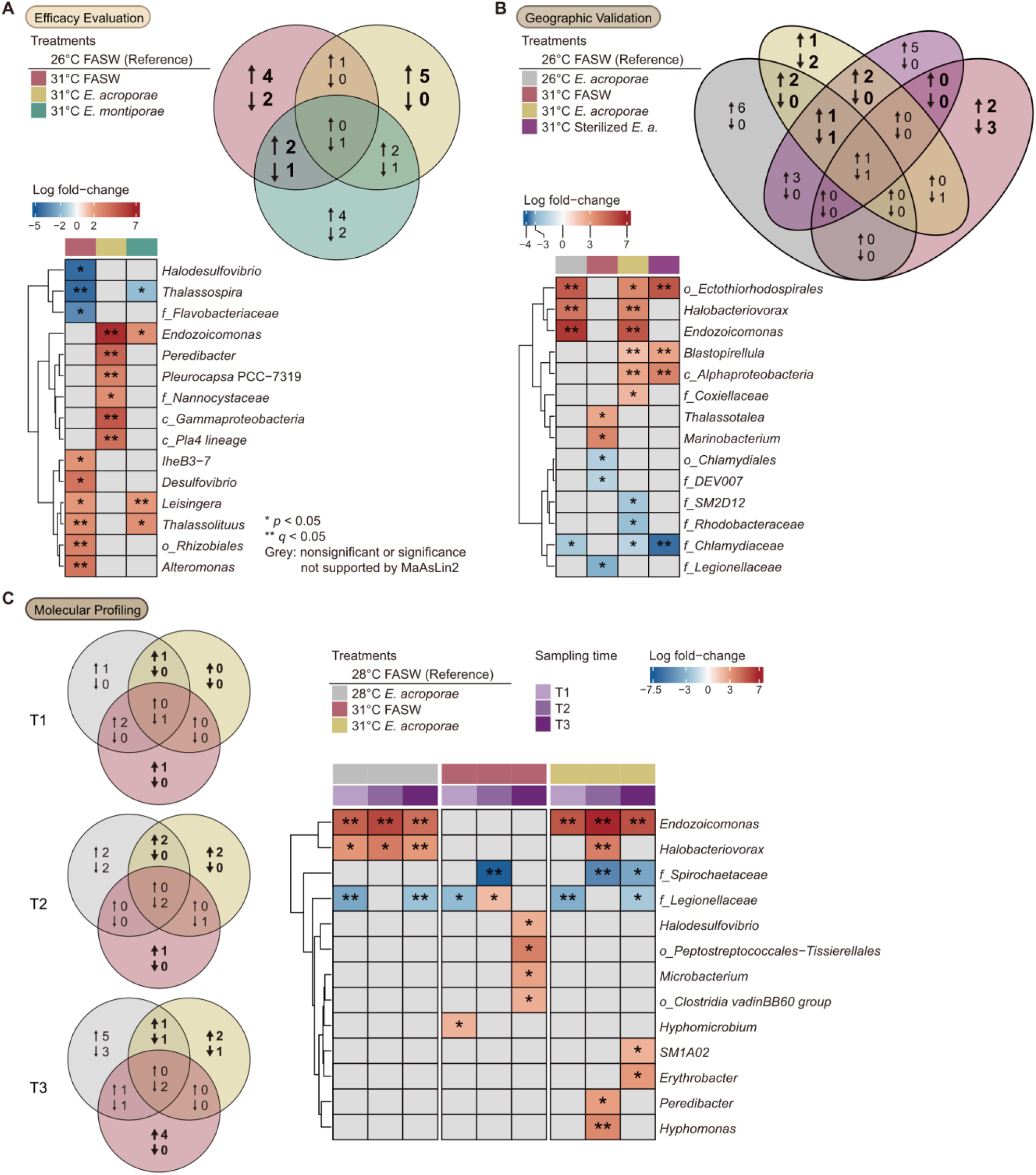
*E. acroporae* Acr-14^T^ suppresses opportunistic pathogens and promotes beneficial microbes. (A-C) Differential abundance of bacterial genera in the efficacy evaluation (A), geographic validation (B), and molecular profiling experiments (C). Venn diagrams summarize differential abundance results among treatments, indicating enriched / depleted genera, with genera of particular interest in bold. Heatmaps illustrate the effect sizes on selected genera. Grey grids represent *p* > 0.05 or results not supported by MaAsLin2 (see Table S2A-S2F). \**p* < 0.05; \*\**q* < 0.05, adjusted by Holm–Bonferroni correction. See Figure S5 and Table S2.

Several beneficial taxa were enriched in *E. acroporae* Acr-14^T^-treated corals. *Thalassospira*, associated with healthy corals,^47^ remained stable in this group, but declined significantly in the 31°C FASW and 31°C *E. montiporae* treatments during the efficacy evaluation experiment (Figure 3A). *Halobacteriovorax*^48^ also showed significant enrichment in *E. acroporae* Acr-14^T^-treated corals in both the geographic validation and molecular profiling experiments (Figures 3B and 3C), suggesting a consistent positive association with *E. acroporae* Acr-14^T^. These findings indicate that *E. acroporae* treatment promotes persistence and / or recruitment of beneficial microbes under heat stress.

Microbial co-occurrence network analysis further supported these observations. Genera enriched in the 31°C FASW or 31°C *E. acroporae* treatments formed distinct clusters in all three experiments (Figures S5D-S5F). Opportunistic pathogens, including *Vibrio*, clustered with 31°C FASW-associated genera (Cluster C2), except in molecular profiling where *Vibrio* and *Alteromonas* formed a separate cluster. In contrast, beneficial taxa such as *Thalassospira* (efficacy evaluation) and P3OB-42 (geographic validation) grouped with *E. acroporae* Acr-14^T^-associated clusters (Cluster C4; Figures S5D and S5E). Moreover, positive correlations between *Endozoicomonas* and *Halobacteriovorax* were consistently observed (Figures S5E and S5F), further suggesting cooperative microbial relationships. Together, these results reinforce the idea that *E. acroporae* Acr-14^T^ functionally modulates the coral microbiome, suppressing opportunists while fostering potentially beneficial taxa, thereby enhancing host resilience under heat stress.

### A chromosome-level genome of *S. pistillata* clade 1 reveals deep divergence from clade 4

To enable transcriptomic analysis of probiotic effects, we generated a chromosome-level genome assembly for the *S. pistillata* clade used in this study, clade 1, which previously lacked genomic resources. This assembly resolved 14 chromosomes based on contact map analysis (Figure S6B). *S. pistillata* has been subdivided into four clades according to geographic distribution and molecular markers (Figure S6A),^49^ with clade 1 occurring across the Pacific–Australia region and clade 4 restricted to the Red Sea–Persian/Arabian Gulf–Kenya area. To assess genomic divergence among clades, we performed pairwise whole-genome comparisons using all available *S. pistillata* assemblies, with *Pocillopora verrucosa* genomes from distinct regions included as comparative controls.

Within-clade comparisons showed high similarity among *S. pistillata* genomes (average nucleotide identity [ANI] > 97.7%, alignment fraction [AF] > 0.79), consistent with values observed among *P. verrucosa* genomes (ANI > 97.6%, AF > 0.75) (Figure S6C). In contrast, comparisons between *S. pistillata* clades 1 and 4 revealed substantial divergence, with ANI values below 91% and AF values under 0.57. Consistent with this, transcriptome alignment efficiency dropped markedly when RNA-seq reads were mapped to non-matching clade references (Figure S6D), highlighting the importance of clade-specific genomic resources for transcriptomic analysis. Together, these results confirm that *S. pistillata* clades are deeply divergent at the genomic level and support previous proposals to recognize them as distinct species.^49^ Our newly assembled clade 1 genome provides a necessary foundation for examining host gene expression in response to probiotic treatment.

### *E. acroporae* treatment alleviates coral stress responses by modulating protein-folding and apoptosis pathways

To understand how *E. acroporae* Acr-14^T^ confers thermal protection, we examined transcriptomic responses of both coral hosts and their algal symbionts. Compared to ambient controls (28°C FASW), most differentially expressed genes (DEGs) in the 28°C *E. acroporae* treatment appeared at T1, whereas the strongest responses in heat-stressed corals (both 31°C treatments) occurred at T2 (Figures 4A and S8C). Overall host gene expression profiles were largely colony-dependent (Figure S7A), whereas symbiont expression reflected both colony and heat stress effects (Figure S8A). Notably, when focusing on DEGs at each time point, host expression profiles at T2 showed clear modulation by probiotic treatment under heat stress (Figure S7B), while a similar pattern was not observed in the symbiont (Figure S8B). These results suggests that the primary effect of *E. acroporae* Acr-14^T^ may be on the coral host.

**Figure 4.**
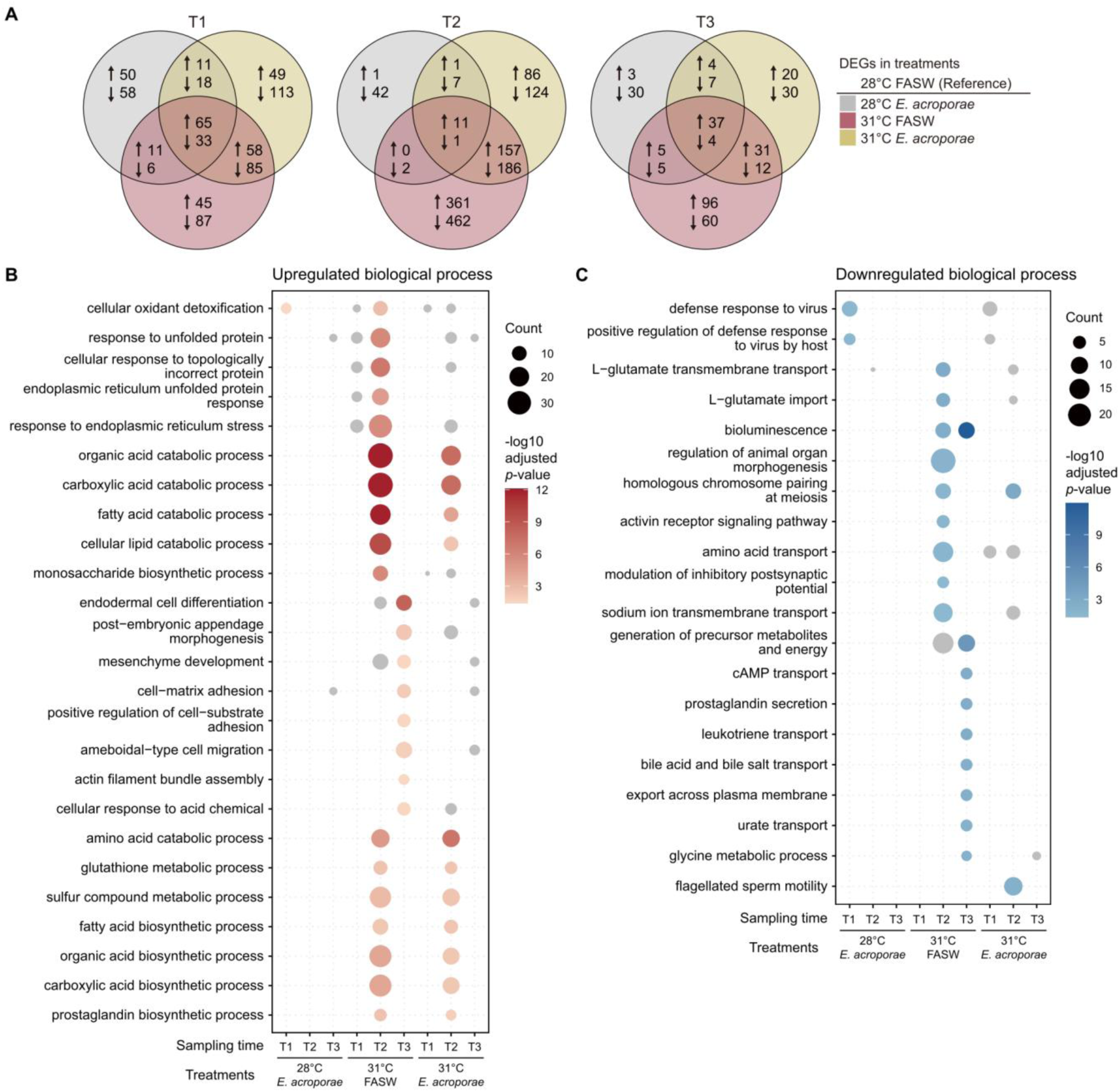
Transcriptomic responses of coral hosts to heat stress and probiotic treatments. (A) Numbers of differentially expressed genes (DEGs) identified in each treatment at each time point, with arrows indicating upregulated and downregulated DEGs (adjusted *p* < 0.05). (B and C) Gene Ontology (GO) enrichment analyses of upregulated (B) and downregulated (C) DEGs among treatments and time points. Bubble size reflects DEG count. Grey bubbles represent GO terms with *p* < 0.05 but adjusted *p* > 0.05 (Benjamini-Hochberg correction). See Figures S7 and S8.

Gene Ontology (GO) enrichment analysis revealed that heat stress (31°C FASW) upregulates genes involved in oxidant detoxification, protein folding, lipid catabolism, and sugar biosynthesis (Figure 4B). By T3, DEGs shifted toward pathways associated with cell differentiation and matrix adhesion. These responses were substantially attenuated in probiotic-treated corals, particularly in pathways related to protein folding and endoplasmic reticulum stress. Additionally, heat stress suppressed genes involved in glutamate transport, energy production, and meiosis, whereas probiotic treatment mitigated many of these suppressive effects (Figure 4C).

Direct comparison between the two 31°C treatments revealed relatively few DEGs (n = 114; Figure S7C), with two genes that are potentially associated with immune responses significantly downregulated in the 31°C *E. acroporae* treatment, including *ANGPTL7* and *VWF*.^50,51^ However, gene set enrichment analysis (GSEA) uncovered consistent downregulation of stress-related pathways in the probiotic-treated group: viral defense at T1, misfolded protein response at T2, and apoptotic signaling at T3 (Figures S7D and S7E). Together, these results suggest that *E. acroporae* Acr-14^T^ buffers the host response to heat stress by alleviating protein-folding and apoptosis-related stress pathways.

In contrast, algal symbionts transcriptional profiles between the two 31°C treatments exhibited fewer differences, and no significant GO enrichment was detected in the 28°C *E. acroporae* group (Figures S8D and S8E). In heat-stressed treatments, upregulated functions enriched at T2 included photosynthesis and chlorophyll biosynthesis (Figure S8D), whereas downregulated genes were associated with nitrate assimilation and peptide biosynthesis (Figure S8E). Notably, significant enrichment of downregulated genes related to ammonium transport was observed only in the 31°C FASW group, suggesting increased ammonium availability in the holobiont. Few DEGs and GSEA signals were detected in the direct comparison between the two 31°C treatments (Figures S8F and S8G), with genes encoding glutamate synthase and glutathione S-transferase specifically upregulated at T2 in the 31°C *E. acroporae* treatment. These findings suggest that *E. acroporae* Acr-14^T^ enhances glutamate synthesis in algal symbionts during heat stress and may facilitate ROS detoxification through glutathione S-transferase upregulation.

### Probiotic treatment modulates coral immune signaling and antioxidant defenses

To complement transcriptomic evidence of reduced stress responses in *E. acroporae*-treated corals, we measured catalase (CAT) and superoxide dismutase (SOD) activities in both coral hosts and their algal symbionts. During heat stress (T1 and T2), coral host CAT activity significantly increased in both the 31°C treatments compared to the control (28°C FASW) (Figure S7F). Algal symbionts also exhibited elevated CAT activity under heat stress, though a significant increase was only observed at T1 in the *E. acroporae* Acr-14^T^-treated group (Figure S7F). Although SOD activity in symbionts tended to rise under heat stress, differences between treatments were not statistically significant at any time (Figure S7G).

Notably, coral host CAT activity was also significantly higher in the 28°C *E. acroporae* treatment than in the 28°C FASW control at T1 (Figure S7F), mirroring transcriptomic results showing upregulation of genes involved in oxidant detoxification (Figure 4B). Additionally, downregulated DEGs associated with antiviral immune responses, e.g., *CGAS, ERBB2,* and *IFIH1*, were enriched in the 28°C *E. acroporae* treatment at T1 (Figure 4C), with similar patterns observed under heat stress in GSEA analysis (Figures S7D and S7E). Together, these findings indicate that *E. acroporae* treatment influences both antioxidant defenses and immune signaling, potentially priming corals to establish symbiosis and to improve stress tolerance.

### A single-cell atlas of *S. pistillata* clade 1 reveals 31 coral cell types

To characterize effects of probiotic treatment at single-cell resolution, we performed single-cell RNA sequencing (scRNA-seq) on four samples, each representing three coral colonies per treatment at T2 (STAR Methods). After filtering ambient background RNA, doublets, and low-quality cells, we retained 35,530 genes across all samples, slightly more than the 33,346 genes detected by bulk RNA-seq from the same colonies. Dimensionality reduction using PCA and UMAP revealed 31 distinct cell clusters in *S. pistillata* clade 1, each defined by uniquely expressed marker genes (Figures 5A and 5B; Table S3).

**Figure 5.**
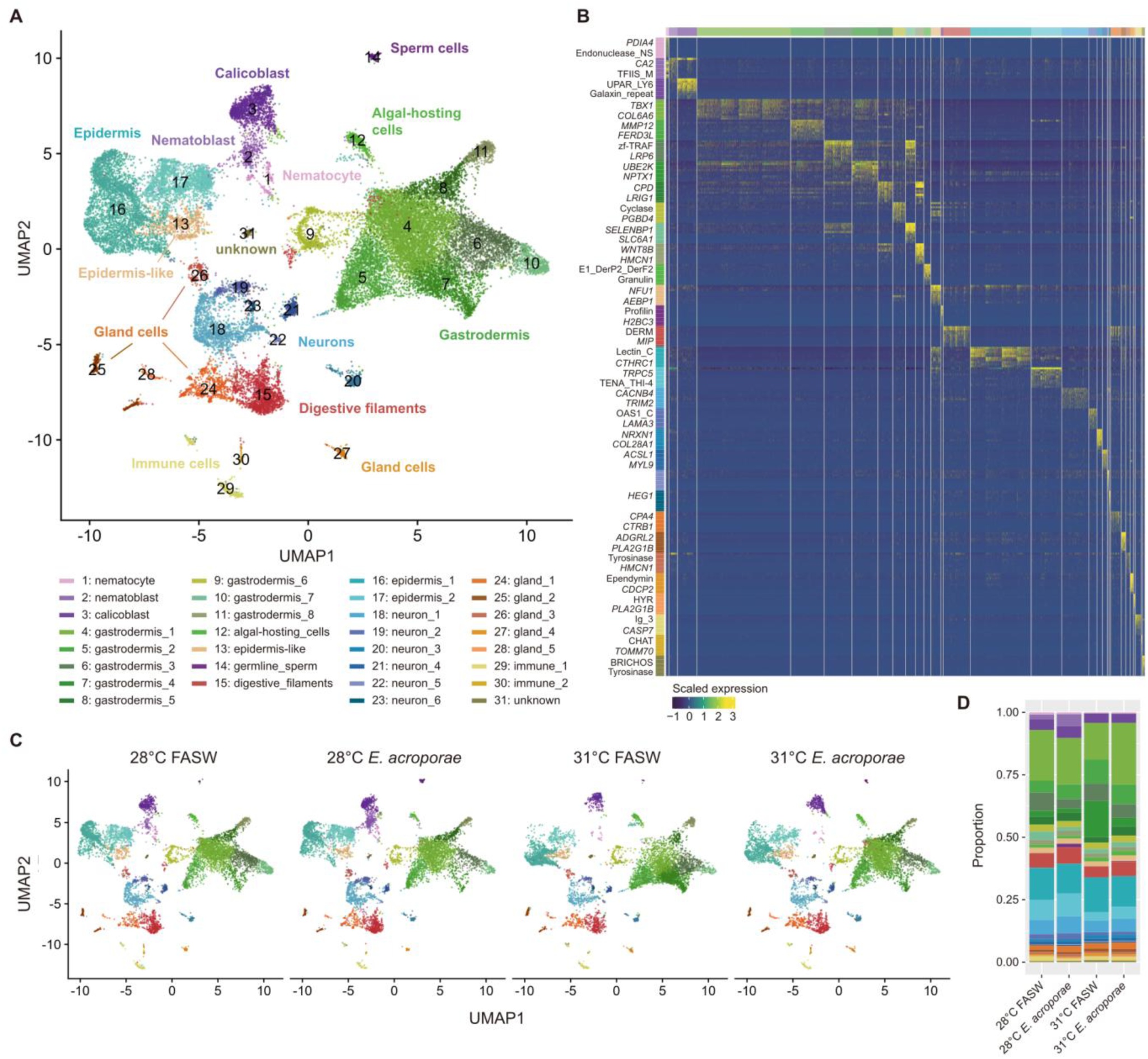
Cell-type atlas of *S. pistillata* clade 1 and its responses to heat stress and probiotic treatment. (A) 2D projection of *S. pistillata* clade 1 single-cell transcriptomes, showing 31 cell clusters across 29,637 cells. (B) Normalized expression of top 10 marker genes (ranked by averaged log2 fold change) in each cell cluster (adjusted *p* < 0.05). Cell types are color-coded on both axes. The x-axis represents all individual cells. Marker genes with annotated gene names or Pfam domains are labeled on the y-axis (a maximum of two labels per cluster, see Table S3). (C) *S. pistillata* cell-type atlas of treatments at T2. (D) Proportion of each cell cluster identified across treatments at T2. See Figures S9, S10, S11 and Table S3.

To assign cell-type identities, we used 491 previously defined marker genes from the *S. pistillata* clade 4 cell atlas,^52^ identifying 280 orthologous markers in clade 1 through reciprocal best hits (RBH; STAR methods; Table S4A). Among these, 181 highly variable markers (average log2 fold-change > 1.7) were used to classify cell clusters. Most clusters were clearly resolved (Figure S9), though cluster 13, initially identified as mitotic host cells, shared several markers with epidermal cells and was reclassified accordingly. Cluster 31 expressed markers associated with multiple cell types and remained unassigned (Figure S9).

Several notable cell types were identified. Nematocytes specifically expressed *POU4F3*, a gene previously linked to these differentiated stinging cells,^53^ whereas their precursors, nematoblasts, shared several markers with nematocytes, but additionally expressed *CA2*, a gene associated with calicoblasts (Figure S10A; Table S3). Secretory potential was evident in several cell types based on the prevalence of secreted protein markers, including gastrodermis_1, epidermis_1, nematocyte, nematoblast, and three distinct gland cell populations (Figure S10B). In contrast, neuron_5 was enriched for GPCR-related genes (Figure S10C), whereas multiple neuronal subtypes exhibited ion channel-associated markers (Figure S10D), consistent with previously defined clade 4 cell types.^52^

We further investigated evolutionary origins of cell-type-specific gene expression patterns through phylostratigraphic analysis.^54^ Cell types such as germline_sperm, gastrodermis_6, and algal-hosting_cells were enriched for ancient markers, whereas nematoblast, epidermis_1, and calicoblast were associated with younger genes (Figure S10E). Notably, both nematocytes and nematoblasts were enriched for Cnidaria-specific markers, whereas calicoblasts showed enrichment for genes that emerged within Hexacorallia and Scleractinia (Figure S10F), echoing findings from the clade 4 cell atlas.^52^ Together, this single-cell atlas provides the first high-resolution view of *S. pistillata* clade 1 cell diversity, establishing a framework to explore how specific cell types contribute to probiotic responses under thermal stress.

### Heat stress alters coral cell-type composition, with indications of probiotic mitigation

We next examined cell-type composition in *S. pistillata* under heat stress and probiotic treatment (Figures 5C and 5D). Cell-type distributions differed significantly among treatments (Chi-squared test, *p* < 2.2e-16). At 31°C, FASW corals showed reduced nematoblast and increased gastrodermis_4, whereas *E. acroporae* treatment kept gastrodermis_4 proportions similar to the control but decreased nematoblast and increased germline_sperm (Figure S11B). The 28°C *E. acroporae* treatment also showed a significant increase in germline_sperm (Figure S11B). These results suggest that *E. acroporae* Acr-14^T^ stabilizes gastrodermis cell types under heat stress and may modulate germline dynamics. However, despite applying a stringent cutoff (FDR < 0.01; |log2FC| > log2(3) ≈ 1.6), the use of a single replicate limits statistical power, and the findings should be considered exploratory and warrant future confirmation.

### Coral heat stress responses occurred predominantly in gastrodermal and epidermal cells

Pseudobulk scRNA-seq gene expression levels were highly correlated with those from bulk RNA-seq across all four treatments (Rs = 0.817–0.825), indicating strong concordance in gene-wise expression patterns between the two methods (Figure S11A). To investigate transcriptomic responses at single-cell resolution, we performed differential expression analyses using the 28°C FASW controls as a reference. The 31°C FASW treatment yielded the highest number of DEGs, while the 28°C *E. acroporae* treatment induced relatively few (Figure S11C). Across all treatments, most DEGs were detected in gastrodermal and epidermal cell populations (Figure S11C), highlighting their sensitivity to environmental and probiotic influences. Because our analyses treated individual cells as replicates, disregarding the inherent correlation between cells from the same sample,^55^ the false positive rate may be inflated and DEGs should be interpreted cautiously.

To strengthen the validity of our interpretation, we focused our GO enrichment analysis on significant GO terms that were also detected in bulk RNA-seq data or supported by previous heat stress-related studies. In the 31°C FASW treatment, GO terms associated with heat stress, such as response to unfolded proteins, endoplasmic reticulum stress, and regulation of apoptotic signaling, were significantly upregulated and enriched in multiple gastrodermal and epidermal cell types (Figure 6). In contrast, metabolic processes, including lipid catabolism and gluconeogenesis, were specifically enriched in epidermal cells (Figure 6). These stress-induced responses were markedly reduced in the 31°C *E. acroporae* group, suggesting that this probiotic mitigates cell-wide transcriptional stress responses, consistent with the bulk transcriptomic results.

**Figure 6.**
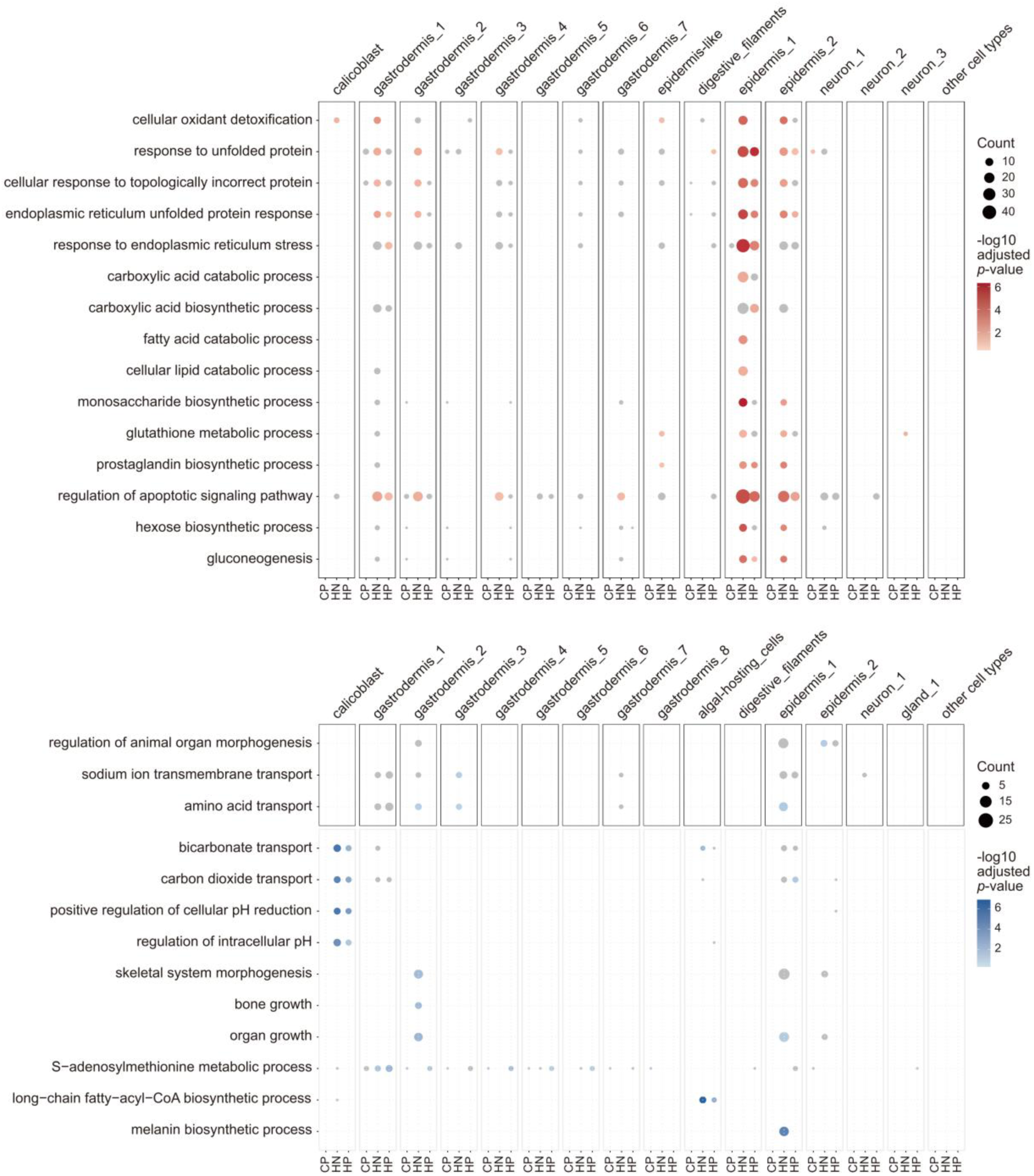
Coral host responses to heat stress and probiotic treatments across cell clusters. GO enrichment analysis of upregulated (red) and downregulated (blue) DEGs in each cell cluster across treatments. GO terms also detected in the bulk transcriptome are shown within black grids, while those identified solely from single-cell transcriptome data are shown within grey grids. Bubble size represents DEG count. Grey bubbles indicate GO terms with *p* < 0.05 but adjusted *p* > 0.05 (Benjamini-Hochberg correction). CP: 28°C *E. acroporae*; HN: 31°C FASW; HP: 31°C *E. acroporae*. See Figure S11.

GO enrichment analysis of downregulated DEGs revealed fewer overlaps with bulk RNA-seq, but highlighted additional cell type-specific effects (Figure 6). For instance, genes related to bicarbonate transport and intracellular pH regulation, processes critical for coral calcification, were downregulated in calicoblasts under heat stress (Figure 6). In algal-hosting cells, DEGs associated with long-chain fatty-acyl-CoA biosynthesis were selectively downregulated. In epidermis_1, genes involved in melanin biosynthesis were suppressed under heat stress, suggesting disruption of protective pigmentation pathways (Figure 6). Notably, the SAMe metabolic process was enriched among downregulated DEGs in gastrodermal cells under both ambient and heat-stressed conditions with *E. acroporae* Acr-14^T^, indicating potential probiotic-mediated modulation of this key methylation pathway (Figure 6).

### *E. acroporae* treatment activated *RET* and suppressed *MAT1A* in heat-stressed corals

To identify candidate genes mediating the observed probiotic effects, we specifically examined DEGs involved in processes potentially related to the probiotic effect, including response to unfolded protein, regulation of apoptotic signaling, melanin biosynthesis, and the SAMe metabolic pathway. We focused on DEGs uniquely detected in either the 31°C FASW or 31°C *E. acroporae* treatments in both bulk and single-cell transcriptomes. In the 31°C FASW treatment, genes involved in the unfolded protein response, such as *PARG*, *HSPH1*, and *ATP2A2*, were significantly upregulated (Figures S12A and S12C). In contrast, *DCTN1*, a gene involved in disassembling stress granules in heat-stressed cells,^56^ was uniquely upregulated in the 31°C *E. acroporae* treatment (Figures S12A, S12C, and S12E). Similarly, pro-apoptotic genes such as *CASP3*, *HYOU1*, and *ERP29* were upregulated only under heat stress without probiotic treatment, whereas *RET*, a gene associated with cell survival signaling, was specifically upregulated in the probiotic-treated group (Figures 7A, 7C, and 7E).

**Figure 7.**
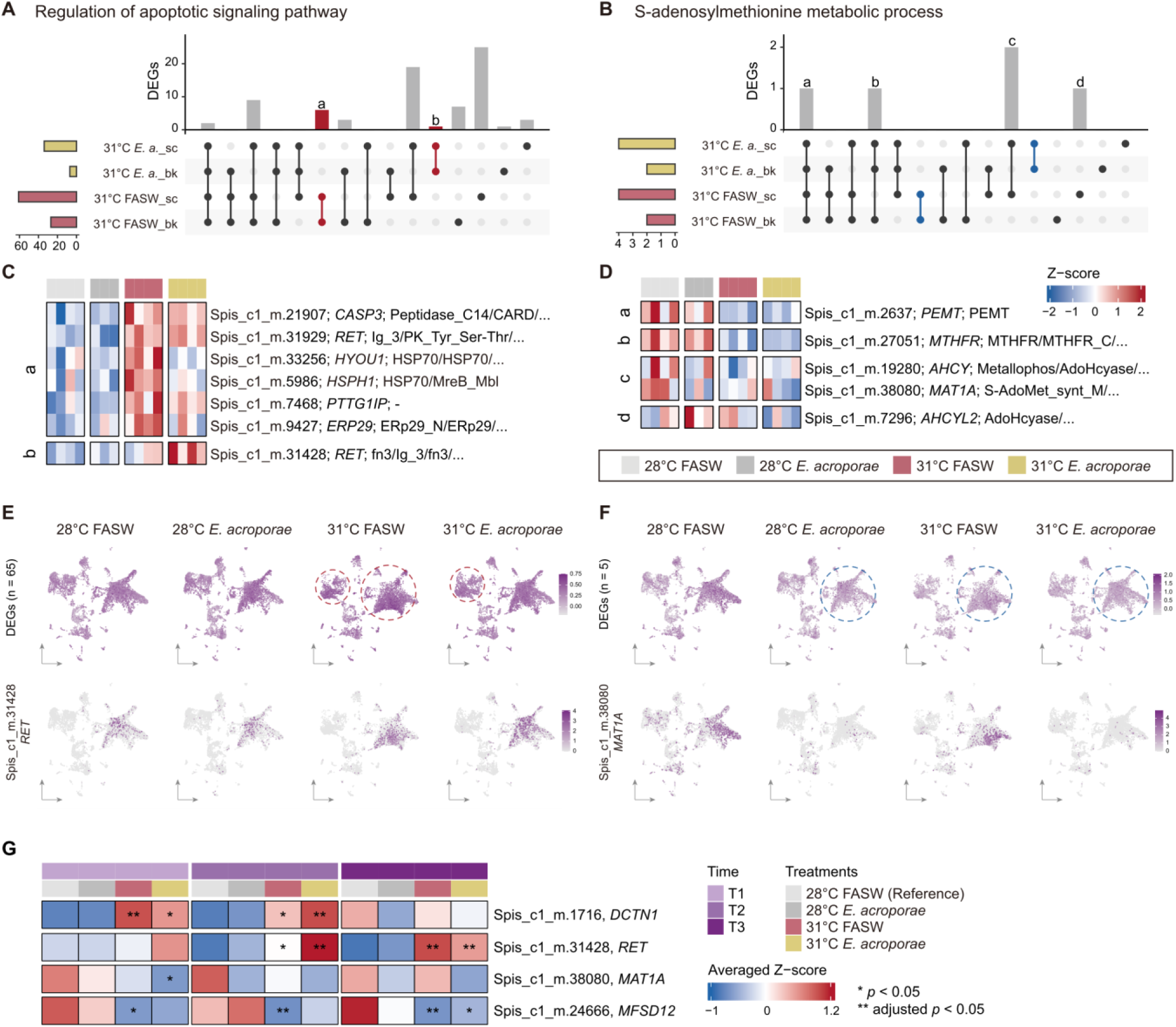
Expression profiles of candidate genes underlying the probiotic effect. (A and B) Overlaps of upregulated (red) and downregulated (blue) DEGs in the 31°C FASW and 31°C *E. acroporae* treatments (both compared to controls), associated with GO terms for regulation of apoptotic signaling pathway (A) and *S*-adenosylmethionine metabolic process (B). DEGs identified by bulk (“bk”) or single-cell (“sc”) transcriptomes are marked. Bar chart letters correspond to genes in panel (C) or (D), with colors highlighting DEGs uniquely detected in either the 31°C FASW or 31°C *E. acroporae* treatments. (C and D) Bulk RNA-seq expression of selected genes associated with the regulation of apoptotic signaling pathway (C) and *S*-adenosylmethionine metabolic process (D). Annotated gene name and Pfam domains are listed after gene ID. (E and F) Single-cell RNA-seq normalized expression of DEGs for regulation of apoptotic signaling pathway (E) or *S*-adenosylmethionine metabolic process (F). Upper panels show all corresponding DEGs, with enriched cell clusters highlighted by dashed circles. (G) Heatmap showing bulk transcriptome z-scaled expression of genes potentially contributing to the probiotic effect, with asterisks indicating statistical significance from DESeq2. (H) Schematic summarizing the proposed mechanisms by which *E. acroporae* Acr-14^T^ enhances the thermal resilience of *S. pistillata*. See Figure S12.

Genes involved in melanin biosynthesis, including *MFSD12* and *KIDINS220*, were downregulated exclusively in the 31°C FASW treatment (Figures S12B, S12D, and S12F), suggesting suppression of pigmentation-related protective mechanisms under heat stress. No SAMe pathway-associated DEGs were detected exclusively in either treatment group in both transcriptomes (Figures 7B and 7D). However, single-cell transcriptomics revealed pronounced downregulation of *MAT1A*, a key gene in SAMe synthesis, in both the 28°C *E. acroporae* and 31°C *E. acroporae* treatments (Figure 7F), indicating that this gene responds strongly to probiotic exposure.

As the single-cell transcriptome data represent only T2, we queried bulk transcriptome data from T1 to T3 to determine whether these gene expression patterns persisted throughout the experiment. *DCTN1* was significantly upregulated in the 31°C *E. acroporae* treatment at T2, but its expression level at T1 was lower than in the 31°C FASW treatment (Figure 7G). In contrast, *RET* expression during heat stress remained consistently higher in the 31°C *E. acroporae* group, whereas *MAT1A* expression was persistently lower in probiotic treatments compared with their corresponding FASW controls, indicating a sustained probiotic effect (Figure 7G). Collectively, these findings suggest that *E. acroporae* Acr-14^T^ mitigates thermal stress by suppressing pro-apoptotic and unfolded protein response pathways, promoting cell survival signaling via early *RET* activation, and modulating SAMe metabolism through *MAT1A* downregulation.

## DISCUSSION

Coral probiotics bolster thermal resilience,^17,23,57^ but their molecular underpinnings remain underexplored. While prior studies have largely focused on shifts in coral microbiota, transcriptomic responses driving probiotic-mediated thermal tolerance have rarely been investigated.^18,20^ This study establishes *E. acroporae* Acr-14^T^ as a potent probiotic that enhances *S. pistillata* thermal tolerance, validated in three independent experiments. We found that *E. acroporae* Acr-14^T^ forms stable symbiosis with recipient corals, suppresses potential pathogens, and modulates host stress signaling. Single-cell transcriptomic analysis reveals that *E. acroporae* Acr-14^T^ downregulates *MAT1A* in the coral host, potentially underpinning its thermal protective effect. The following sections evaluate evidence for *E. acroporae* Acr-14^T^ efficacy, explore its symbiosis formation strategy, review its restructuring of the coral microbiome, and explain molecular mechanisms driving thermal resilience.

### Evaluating effectiveness of *E. acroporae* Acr-14^T^

Validating probiotic effects in coral studies requires robust negative controls, typically placebo treatments like sterilized saline or seawater.^21^ By consistently comparing with the FASW control in each experiment, we repeatedly demonstrated that *E. acroporae* Acr-14^T^ mitigated *S. pistillata* heat stress responses. Moreover, detection of *E. acroporae* Acr-14^T^ in recipient corals confirmed active colonization, linking its presence to enhanced thermal tolerance. These findings set the stage for evaluating alternative microbial controls.

Selecting a non-beneficial microbe as a negative control is controversial, as some bacteria may confer unexpected benefits.^58^ Although our prior study suggested that *E. montiporae* CL-33^T^ benefits corals,^31^ the present results showed no enhancement of thermal tolerance and no evidence of stable symbiosis in *S. pistillata*. The absence of beneficial effects and poor colonization therefore supports the utility of *E. montiporae* CL-33^T^ as a negative control for *S. pistillata*, reinforcing the probiotic effectiveness of *E. acroporae* Acr-14^T^. Notably, probiotic efficacy varies with host species and experimental goals,^21^ and even the same *Endozoicomonas* lineage may elicit distinct responses across coral genera.^59^ The potential of *E. montiporae* CL-33^T^ in other contexts, such as in different coral hosts or for alternative beneficial functions, warrants further exploration.

*E. acroporae* spans a broad geographic range and diverse coral hosts,^30^ including *S. pistillata* in Taiwan.^32,33^ Our results show that *E. acroporae* Acr-14^T^ is effective in *S. pistillata* from two distinct locations, potentially bridging biogeographic boundaries between tropical coral reefs and non-reefal coral communities.^36^ Successful establishment of the same bacterial species in juvenile *Acropora kenti* from Australia through manual inoculation further highlights its host flexibility.^60^ Although the heat tolerance of treated *A. kenti* remains untested, these findings suggest a potential cross-species and cross-regional role for this coral probiotic.

Despite these promising outcomes, the *in-situ* performance of *E. acroporae* Acr-14^T^ remains unverified. Field validation is critical, yet few coral probiotic studies have reached this stage. Notably, one trail demonstrated that probiotic treatment can halt stony coral tissue loss disease (SCTLD) progression,^61^ and another reported no detectable off-target effects after three months of continuous application,^62^ supporting the feasibility of *in-situ* use. However, *Endozoicomonas* abundance often declines during bleaching,^32,63,64^ independent of Symbiodiniaceae loss,^65^ suggesting that this symbiosis may not persist under thermal stress.

Strategies such as repeated inoculation during heat events may facilitate the efficacy of *E. acroporae* Acr-14^T^ in natural environments.

### Establishment of symbiosis between *S. pistillata* and *E. acroporae* Acr-14^T^

We found consistent evidence from both 16S sequencing and FISH analyses that *E. acroporae* Acr-14^T^ establishes symbiosis with recipient corals. Notably, *E. acroporae* Acr-14^T^ formed CAMAs in *S. pistillata* tissues, similar to those observed in *A. kenti* treated with the same bacterial species.^60^ In nature, CAMAs are commonly formed by *Endozoicomonas*^66^ and are thought to contribute to host phosphate cycling.^39^ Our findings show that CAMAs can be artificially induced in *S. pistillata* using *E. acroporae* Acr-14^T^, providing a tractable model to study their functional roles and the molecular mechanisms driving *Endozoicomonas*-coral symbiosis, which remain poorly understood.

Genomic analyses indicate that *Endozoicomonas* genomes are enriched in eukaryotic-like proteins (ELPs),^67^ with ephrin ligand genes proposed to mediate host interaction.^68,69^ However, *E. acroporae* Acr-14^T^ and related strains lack these coral-like ephrin ligand genes, suggesting alternative colonization strategies.^69^ Our bulk transcriptomic data revealed significant downregulation of innate immune genes, including *CGAS*, in corals treated with *E. acroporae* Acr-14^T^, implying that immune suppression may facilitate symbiosis. Similarly, pathogens such as *Mycobacterium tuberculosis* manipulate the host cGAS-STRING pathway to establish infection in human macrophages.^70^ Given that *Endozoicomonas* species are pathogenic in other organisms,^27^ *E. acroporae* Acr-14^T^ may employ a pathogenic-like strategy to initiate mutualism in corals. Further studies are needed to validate molecular pathways involved.

### *E. acroporae* Acr-14^T^ suppresses potential coral pathogens and promotes beneficial microbes

Host microbiome restructuring is a widely reported effect in coral probiotic studies, regardless of the candidate tested.^21^ Such shifts may arise from direct probiotic effects, including antimicrobial production and competitive exclusion,^15,17,20,62^ or indirectly via ROS scavenging and host immunity modulation,^18,71^ thereby steering the microbiome toward a beneficial state. *Endozoicomonas* abundance often correlates negatively with coral stress or disease,^64,72^ a pattern mirrored in our study, where *E. acroporae* Acr-14^T^ suppressed opportunistic pathogens in *S. pistillata*. Taxa linked to SCTLD (*Alteromonas*, *Halodesulfovibrio*, *Peptostreptococcales-Tissierellales*),^42,44,45^ black band disease (*Desulfovibrio*),^41^ and thermal stress (*Legionellaceae*)^43^ were all reduced in treated corals.

Mechanistically, one possible explanation lies in the regulation of oxidative stress. *E. acroporae* Acr-14^T^ exhibits potent ROS-scavenging capacity,^29^ potentially helping to maintain redox homeostasis and limit pathogen-favored niches. Supporting this, probiotic treated corals showed elevated host catalase activity under ambient conditions and upregulation of a symbiont ROS detoxification gene during heat stress. Direct measurements of ROS levels using assays such as DCFH-DA could further clarify the extent of redox modulation mediated by *E. acroporae* Acr-14^T^ in corals.

Beyond pathogen control, *E. acroporae* Acr-14^T^ promoted health-associated taxa, including *Thalassospira*, P3OB-42, and *Halobacteriovorax*. *Thalassospira* has been associated with healthy corals^47^ and may contribute to carbon and phosphorus cycling.^73^ P3OB-42 and *Halobacteriovorax* are predatory bacteria known for their roles in coral disease resistance.^74,75^ Notably, *Halobacteriovorax* has been used successfully to control *Vibrio* infections in corals.^48^ Such restructuring of the microbial network may foster interactions that outcompete or prey upon pathogens, enhancing coral resilience. Future work isolating and co-culturing these taxa with *E. acroporae* Acr-14^T^ could clarify their functional interactions and guide the design of optimized probiotic consortia for assisting coral health.

### Cellular insights into coral thermal stress

We found that coral heat stress responses primarily occurred in gastrodermal and epidermal cells, with strong activation of genes involved in protein-folding, ER stress, and apoptotic signaling. This finding is consistent with those of prior studies that localized heat stress-associated gene expression to gastrodermal and epidermal tissues.^76^ Histological observation further supports this, revealing pronounced gastrodermal tissue damage under heat stress.^77^ Notably, we observed an increased proportion of gastrodermis_4 cells in heat-stressed corals, a shift that did not occur in the *E. acroporae* Acr-14^T^–treated group. This suggests that *E. acroporae* Acr-14^T^ may alleviate thermal stress impacts in gastrodermal cells, consistent with our functional enrichment results.

Our single-cell transcriptomic data also captured responses characteristic of aposymbiotic corals. In particular, the long-chain fatty-acyl CoA biosynthetic pathway, essential for lipid synthesis and energy metabolism, was significantly downregulated in algal-hosting cells under heat stress. As Symbiodiniaceae contribute fatty acids to the coral host,^78,79^ reduced fatty acid availability during bleaching likely drives this downregulation. We also found a heat-induced decline in melanin expression, a key innate immune molecule in cnidarians.^80^ While a prior study reported decreased melanin during bleaching,^81^ our results pinpoint this reduction to epidermal cells, suggesting impaired immune defenses and higher pathogen susceptibility. Notably, *E. acroporae* Acr-14^T^ treatment mitigated this effect, potentially enhancing coral immunity during thermal stress.

Consistent with prior studies,^82,83^ we observed suppressed calcification under heat stress, driven by disrupted bicarbonate transport in calicoblasts. This finding aligns closely with a recent single-cell study of *Orbicella faveolata*, which reported similar downregulation.^84^ In addition, heat-stressed *O. faveolata* exhibited a reduced proportion of cnidocytes.^84^ In the *S. pistillata* model, we determine that this decline likely occurred in specifically nematoblasts, which *E. acroporae* Acr-14^T^ treatment did not rescue, indicating high thermal sensitivity of this cell type.

### Mechanisms underlying coral thermal tolerance enhanced by *E. acroporae* Acr-14^T^

Heat-induced coral bleaching involves multiple factors, including elevated temperature-driven ROS accumulation, which causes protein misfolding and ER stress, further amplifies ROS production, and ultimately triggers bleaching and apoptosis.^85^ Our findings suggest that *E. acroporae* Acr-14^T^ may alleviate protein-folding stress in heat-stressed corals through both ROS scavenging and *RET* upregulation. ROS-scavenging bacteria are known to reduce ROS levels^16,71^ and suppress apoptotic signaling^20^ in heat-stressed corals. As a potent ROS scavenger, *E. acroporae* Acr-14^T^ likely mitigates ROS accumulation, thereby alleviating protein-folding stress. In addition, we observed specific *RET* upregulation in probiotic-treated corals under heat stress. Given *RET*’s role in promoting cell survival and stress tolerance,^86,87^ its early activation may help dampen coral stress responses during thermal exposure.

Beyond ROS, heat stress also disrupts nutrient cycling between coral hosts and algal symbionts. Elevated host energy demand under heat stress activates catabolic pathways, shifting nitrogen balance from net uptake to ammonium release in the coral holobiont. Excess nitrogen promotes symbiont proliferation and retention of photosynthates, reducing carbon transfer to the host and destabilizing the symbiosis.^88^ In this study, downregulation of the symbiont ammonium transport pathway was detected only in the 31°C FASW treatment. This pattern may reflect *E. acroporae* Acr-14^T^–mediated thermal protection that helps maintain host energy homeostasis, as indicated by the absence of downregulated energy-production genes and only minor upregulation of catabolic pathways in probiotic-treated corals. Such regulation may limit ammonium release, indirectly preserving coral–algal symbiosis under heat stress.

Importantly, our single-cell transcriptomic analysis revealed significant downregulation of *MAT1A* in *E. acroporae* Acr-14^T^-treated corals, independent of temperature. *MAT1A* is responsible for SAMe synthesis in mammals and is homologous to the *sams-1* gene in *Caenorhabditis elegans*.^89^ Notably, *sams-1* knockdown markedly enhances thermal tolerance in *C. elegans*,^90^ possibly by altering 1-carbon cycle function^91^ or phosphatidylcholine synthesis.^92^ *E. acroporae* Acr-14^T^ may confer similar benefits in corals by suppressing *MAT1A.* While *sams-1* knockdown also increases pathogen susceptibility in *C. elegans*,^93^ this effect was not observed in corals treated with *E. acroporae* Acr-14^T^, highlighting the dual role of this probiotic in mitigating the trade-off between stress tolerance and immune defense.

Overall, our findings demonstrate that *E. acroporae* Acr-14^T^ enhances thermal tolerance in *S. pistillata* by alleviating protein-folding stress, scavenging ROS, and modulating pro-survival and SAMe synthesis pathways. This strain also consistently suppressed potential pathogens while promoting beneficial microbial communities. We propose that these effects help corals conserve energy and stabilize coral-Symbiodiniaceae nutritional balance under heat stress, thereby mitigating bleaching (Figure 7H). Given the urgency of coral conservation, *in-situ* validation of *E. acroporae* Acr-14^T^ should be a priority for future studies to advance its application in reef restoration. Importantly, while *E. acroporae* Acr-14^T^ showed clear benefits in *S. pistillata*, these findings should not be generalized to all *Endozoicomonas*, as genomic evidence indicates that potential coral pathogens may also occur within this genus.^28^

## RESOURCE AVAILABILITY

### Lead contact

Requests for further information and resources should be directed to the lead contact, Sen-Lin Tang.

### Materials availability

This study did not generate new unique reagents.

### Data and code availability

All the raw data generated in this study have been deposited in NCBI BioProject under accession number PRJNA1248900. The Whole Genome Shotgun project for *Stylophora pistillata* has been deposited at GenBank under the accession JBNOSB000000000. The version described in this paper is JBNOSB010000000. Code used in this study and additional processed data are available on Zenodo: https://doi.org/10.5281/zenodo.15335716 and is publicly available as of the date of publication. Any additional information in this paper is available from the lead contact upon request.

## ACKNOWLEDGMENTS

We would like to thank Shinya Shikina, Chang-Feng Dai, Yu-Wei Wu, Mei-Fang Lin, Jih-Terng Wang, Lee Li Chuen, and Yung-Chi Tu for their valuable advice and suggestions regarding experimental design and data analyses. We thank Crystal J. McRae and Tung-Yung Fan for arranging coral acclimation at the National Museum of Marine Biology and Aquarium. We acknowledge the Neuroscience Core Facility at Academia Sinica (AS-CFII-110-101) for providing laser scanning confocal microscope services. We thank miss.misi.xu for illustrating the schematic probiotic mechanisms and inoculation procedures. We thank Steven D. Aird for English language editing. We thank Huei-Ying Li for conducting the PacBio full-length 16S rRNA gene amplicon sequencing. This study was supported by Academia Sinica (AS-SS-111-05 and AS-IV-114-L03) awarded to Sen-Lin Tang. Yi-Jyun Luo was supported by Academia Sinica (AS-CDA-112-L06) and National Science and Technology Council (113-2311-B-001-026).

## AUTHOR CONTRIBUTIONS

Conceptualization, C.-Y.L., Y.-J.L., and S.-L.T.; formal analysis, C.-Y.L., Y.-J.L., V.M.P.-G., T.D.L., and Y.-P.C.; funding acquisition, S.-L.T.; investigation, C.-Y.L., Y.-P.C., K.-N.S., P.-S.C., S.-P.Y., Z.-R.Y., Y.-H.C., M.C., C.-H.C., Y.-J.C., H.-F.C., and J.-H.Y.; methodology, C.-Y.L., Y.-P.C., N.W., Y.-H.C., Y.-L.C., and Y.- J.L.; project administration, C.-Y.L. and S.-L.T.; resources, Y.-L.C., I.J.-Y.L., M.-Y.J.L., and Y.-J.L.; supervision, C.-Y.L., N.W., M.-Y.J.L., Y.-J.L, and S.-L.T.; validation, C.-Y.L., Y.-P.C., P.-S.C., and S.-L.T.; visualization, C.-Y.L., Y.-P.C., T.D.L., N.W., Y.-J.L., and S.-L.T.; writing – original draft, C.-Y.L.; writing –review & editing, C.-Y.L., Y.-P.C., T.D.L., V.M.P.-G., Y.-J.L., and S.-L.T.

## SUPPLEMENTAL INFORMATION

**Table S1. *Endozoicomonas* species identification and removed ASVs, related to Figure 2 and STAR Methods**

(A) *Endozoicomonas* species-level taxonomy assignment

(B) Removed ASVs in short-length 16S sequencing data

(C) Removed ASVs in full-length 16S sequencing data

**Table S2. Differential abundance analysis of coral microbiota, related to Figure 3**

(A) Results of efficacy evaluation

(B) Results of geographic validation

(C) Results of molecular profiling at T0

(D) Results of molecular profiling at T1

(E) Results of molecular profiling at T2

(F) Results of molecular profiling at T3

**Table S3. Marker genes identified in the *S. pistillata* clade 1 single-cell atlas, related to Figure 5**

**Table S4. *S. pistillata* marker genes RBH and primer sequences used in this study, related to Figure S9 and STAR Methods**

(A) *S. pistillata* marker genes RBH

(B) primer sequences used in this study

**Table S5. *S. pistillata* clade 1 gene annotation, related to Figure S10**

**Table S6. Probiotic inoculation concentrations and water parameters, related to STAR Methods**

(A) Collected bacterial cultures and inoculation concentrations

(B) Experimental tank water parameters

## FIGURE TITLES AND LEGENDS

**Figure S1. Schematic overview of experimental tanks and replication design for each experiment, related to Figures 1 and S2**

(A) Schematic of the experimental tank used in the efficacy evaluation experiment.

(B) Schematic of the experimental tank used in the geographic validation and molecular profiling experiments.

(C) Replication design for the three coral bleaching experiments. Different coral colonies are represented by distinct shapes, with individual colony codes labeled on the left. The shape and fill color indicate which colonies were included in different assays.

**Figure S2. Study design and coral bleaching responses of efficacy evaluation and geographic validation, related to Figures 1 and S1**

(A) Treatment and replication design of efficacy evaluation. Each treatment consisted of one aquarium containing five coral nubbins, each originating from a different colony (n = 5).

(B) Treatment and replication design of geographic validation. Each treatment consisted of two aquarium systems: one for sampling and another for measuring coral photosynthetic efficiency. Each aquarium contained four coral nubbins, each from a different colony (n = 4).

(C and D) Detailed treatment regimens for efficacy evaluation (C) and geographic validation (D), including temperature settings and dates for feeding, probiotic inoculation, and sample collection.

(E and F) Photosynthetic efficiency (Fv/Fm) of nubbins in efficacy evaluation (E) and geographic validation

(F). In boxplots, dots represent outliers, and dashed lines connecting treatment means (n = 5 in efficacy evaluation; n = 4 in geographic validation; One-way ANOVA with Tukey post-hoc test).

**Figure S3. Effects of treatments on coral microbial composition and diversity, related to Figures 2 and 3 and STAR Methods**

(A-C) Relative abundance of bacterial class-level composition across treatments in efficacy evaluation (A), geographic validation (B), and molecular profiling (C), determined by short-length 16S rRNA gene sequencing. Tank water samples (ws); lost or failed to sequence samples (x).

(D) Relative abundance of *E. acroporae* in samples, determined by full-length 16S rRNA gene sequencing. All ASVs belonging to genus *Endozoicomonas* were assigned to *E. acroporae*.

(E-G) ASV richness and Shannon index across treatments in efficacy evaluation (E), geographic validation (F), and molecular profiling (G) (efficacy evaluation: n = 5, geographic validation: n = 4, molecular profiling: n = 5; Wilcoxon signed-rank tests; \**p* < 0.05; \*\**p* < 0.01).

(H) Rarefaction curves for all samples in molecular profiling, with data derived from short-length or full-length sequencing methods.

(I) ASV richness and Shannon index comparing coral (n = 78) and tank water (n = 16) samples in molecular profiling (Wilcoxon signed-rank tests).

(J) Principal coordinates analysis (PCoA) illustrating microbial community differences between coral (n = 78) and tank water (n = 16) samples in molecular profiling (PERMANOVA).

**Figure S4. FISH detection of *Endozoicomonas* in coral tissues and bacterial cultures, related to Figure 2 and STAR methods**

(A) Individual and merged channels showing a tissue section from the 31°C *E. acroporae* treatment in molecular profiling, with *Endozoicomonas* signals visible. Symbiodiniaceae exhibited strong autofluorescence under Alexa 488. ca: calicoblastic cell layer; Endo: *Endozoicomonas*; ep: epidermal layer; me: mesenterial filaments; Sym: Symbiodiniaceae. Scale bars: 50 µm.

(B) Individual and merged channels showing *Endozoicomonas* signals at higher magnification in a sample from the 28°C *E. acroporae* treatment in molecular profiling. Scale bars: 5 µm.

(C) Size of *Endozoicomonas* aggregations observed in microscope slides across the four treatments in molecular profiling (two samples per treatment, five slides per sample, 40 slides in total).

(D) Validation of FISH probe specificity using the universal prokaryotic probe EUB338mix and the Endo-Group B probe on three bacterial cultures: *E. acroporae* Acr-14^T^, *Vibrio coralliilyticus* YB strain, and *Pseudoalteromonas piratica.* Scale bars: 10 µm.

**Figure S5. Multivariate and network analyses of coral microbial community, related to Figure 3**

(A-C) Principal coordinates analysis (PCoA) illustrating the effects of experimental batches (A) or treatments in efficacy evaluation (B) and geographic validation (C) on coral microbial composition. 95% confidence ellipses were drawn around the centroid of each group (PERMANOVA; pairwise comparison *p* values adjusted using Bonferroni correction).

(D-F) SparCC correlation networks of the 60 most prevalent bacterial genera in efficacy evaluation (D), geographic validation (E), and molecular profiling (F). Small panels show networks overlaid with genera enriched specifically in the 31°C FASW or 31°C *E. acroporae* treatments. Clusters identified in these networks are labeled with “C” followed by a number.

**Figure S6. Genetic divergence between *S. pistillata* clades 1 and 4, related to STAR methods**

(A) Phylogenetic tree of *S. pistillata* ITS rDNA sequences obtained from this study, a released genome (GCA_964205215.1), and a previous study. The tree is rooted using an outgroup (*Seriatoproa* sp.).

(B) Hi-C contact map of *S. pistillata* clade 1 and menthol-induced bleaching treatment. Scale bars: 1 mm.

(C) Pairwise whole-genome alignment illustrating genetic divergence between samples. Colors in pie chart indicate genomes used for alignment.

(D) Overall alignment rate of RNA sequences mapped to clade 1 or clade 4 *S. pistillata* genomes. Each dot represents one sample, and black diamonds indicate group means (Wilcoxon signed-rank tests).

**Figure S7. Effect of heat stress and probiotic treatments on the coral host transcriptome, related to Figure 4**

(A and B) PCA plots illustrating coral host transcriptomic profiles, constructed using all pre-filtered genes

(A) or all DEGs identified at each sampling time (B). Numbers in parentheses indicate numbers of genes used for PCA construction and total gene count in the genome-guided (*S. pistillata* clade 1) transcriptome.

(C) Volcano plots showing results of differential expression analyses between the 31°C FASW and 31°C *E. acroporae* treatments. Numbers of upregulated and downregulated DEGs are indicated by n.

(D) GSEA of genes ranked by Wald statistic from DESeq2, comparing the 31°C FASW and 31°C *E. acroporae* treatments. The five most significantly enriched GO terms in each biological process category are showed (adjusted *p* < 0.05).

(E) GSEA running scores for significant GO terms identified at T1, T2, and T3.

(F and G) Catalase (F) and superoxide dismutase (G) activities in coral tissue and algal symbionts across the four treatments and sampling times. Activities were normalized to the control treatment (28°C FASW) at each sampling time (*adjusted *p* < 0.05, **adjusted *p* < 0.01; n = 5, One-way ANOVA with Tukey post-hoc test).

**Figure S8. Effect of heat stress and probiotic treatments on the algal symbiont transcriptome, related to Figure 4**

(A and B) PCA plots showing algal symbionts transcriptomic profiles. PCA was conducted using all pre-filtered genes (A) or all DEGs identified at each time point (B). Numbers in parentheses at the top indicate genes used for PCA construction and total gene count from the *C. goreaui* genome.

(C) Numbers of DEGs identified in algal symbionts from each treatment at each sampling time. Arrows indicate upregulated and downregulated DEGs (adjusted *p* < 0.05).

(D and E) GO enrichment analyses of upregulated (D) or downregulated (E) DEGs identified in algal symbionts across treatments and sampling time points. Bubble size reflects DEG count. Grey bubbles indicate GO term with *p* < 0.05 but adjusted *p* > 0.05 (Benjamini-Hochberg correction).

(F) Volcano plots showing differential expression analysis results between the 31°C FASW and 31°C *E. acroporae* treatments. Number of upregulated and downregulated DEGs are indicated by n.

(G) Gene set enrichment analysis (GSEA) of genes ranked by Wald statistic from DESeq2 results, comparing the 31°C FASW and 31°C *E. acroporae* treatments. Significantly enriched GO terms (adjusted *p* < 0.05) within the biological process category are showed.

*Note: Due to severe bleaching, DEGs from the 31°C FASW treatment at T3 were derived from 2 algal symbiont samples (colonies A and B, n = 2, STAR Methods).

**Figure S9. Cell-type classification based on orthologous marker genes identified between *S. pistillata* clades 1 and 4 cell markers, related to Figure 5**

Bump plot illustrating orthologous marker genes defined between *S. pistillata* clades 4 and 1 cell markers. The left column lists 280 markers belong to *S. pistillata* clade 4 (see Table S4). Orthologous markers are connected only to highly variable markers identified in *S. pistillata* clade 1 (average log2 fold change > 1.7; right column, 181 genes in total). Line thickness indicates average log2 fold change value. Note that marker genes identified in *S. pistillata* clade 1 may not be exclusive to a single cell cluster. Heatmap shows normalized expression of orthologous marker genes across cells.

**Figure S10. *S. pistillata* clade 1 cell atlas marker gene functions and phylostratigraphy, related to Figure 5**

(A) Expression profiles illustrating that nematoblasts share multiple marker genes with nematocytes and one marker gene with calicoblasts.

(B) Percentage of predicted secreted proteins (containing a signal peptide and no transmembrane domains) among marker genes (with average log2 fold change > 1) for each cell cluster.

(C) Total number of GPCRs genes (containing Pfam domains 7tm_1, 7tm_2, or 7tm_3) identified among marker genes (with average log2 fold change > 1) for each cell cluster.

(D) Total number of ion channels genes (based on Pfam domains, see Table S5) identified among marker genes (with average log2 fold change > 1) for each cell cluster.

(E) Mean age of marker genes (with average log2 fold change > 1.7) for each cell cluster, as defined by phylostratigraphy.

(F) Enrichment or depletion of marker genes within each cell cluster among phylostrata. Statistical significance determined by Fisher exact test (adjusted *p* < 0.1, Benjamini-Hochberg correction).

**Figure S11. Concordance between bulk and single-cell transcriptomes and treatment effects on coral cell clusters, related to Figures 5 and 6**

(A) Gene-wise comparison of pseudobulk scRNA-seq and bulk RNA-seq expression levels, with the Spearman correlation coefficient indicated.

(B) Relative differences in cell cluster proportions between control and other treatments. Red indicates significant changes (FDR < 0.01 and |Log2FC| > 1.6, permutation test).

(C) Numbers of DEGs (adjusted *p* < 0.05) identified in each cell cluster.

**Figure S12. Additional gene expression profiles potentially contributing to the probiotic effect, related to Figure 7**

(A and B) Overlaps of upregulated (red) and downregulated (blue) DEGs in the 31°C FASW and 31°C *E. acroporae* treatments (both compared to controls), associated with GO terms for response to unfolded protein (A) and melanin biosynthetic process (B). DEGs identified by bulk (“bk”) or single-cell (“sc”) transcriptomes are marked. Bar chart letters correspond to genes in panel (C) or (D), with colors highlighting DEGs uniquely detected in either the 31°C FASW or 31°C *E. acroporae* treatments.

(C and D) Bulk RNA-seq expression of selected genes associated with the response to unfolded protein

(C) and melanin biosynthetic process (D). Annotated gene name and Pfam domains are listed.

(E and F) Single-cell RNA-seq normalized expression of DEGs for response to unfolded protein (E) or melanin biosynthetic process (F). Upper panels show all corresponding DEGs, with enriched cell clusters highlighted by dashed circles.

**Figure S13. Detailed probiotic inoculation and feeding procedures, related to STAR methods**

Step 1: Water circulation between the coral tank and its sump was halted by turning off the submersible pump and disconnecting silicon tubing. Step 2: Concentrated probiotic solutions or placebo (FASW) were added to the coral tank, with water flow maintained by an air pump during inoculation. Step 3: After two hours of inoculation, coral nubbins were rinsed with FASW and transferred to a new coral tank prefilled with temperature-adjusted fresh ASW. Step 4: Once all coral nubbins were transferred, the used coral tank was replaced and cleaned. Finally, all silicon tubing was reconnected, and the submersible pump was reactivated to resume water circulation with the sump. In efficacy evaluation, as sumps were not used, procedures involving the sump were not required. For feeding, following the same procedure, replacing the probiotic solution with concentrated Artemia.

## STAR★METHODS

### KEY RESOURCES TABLE

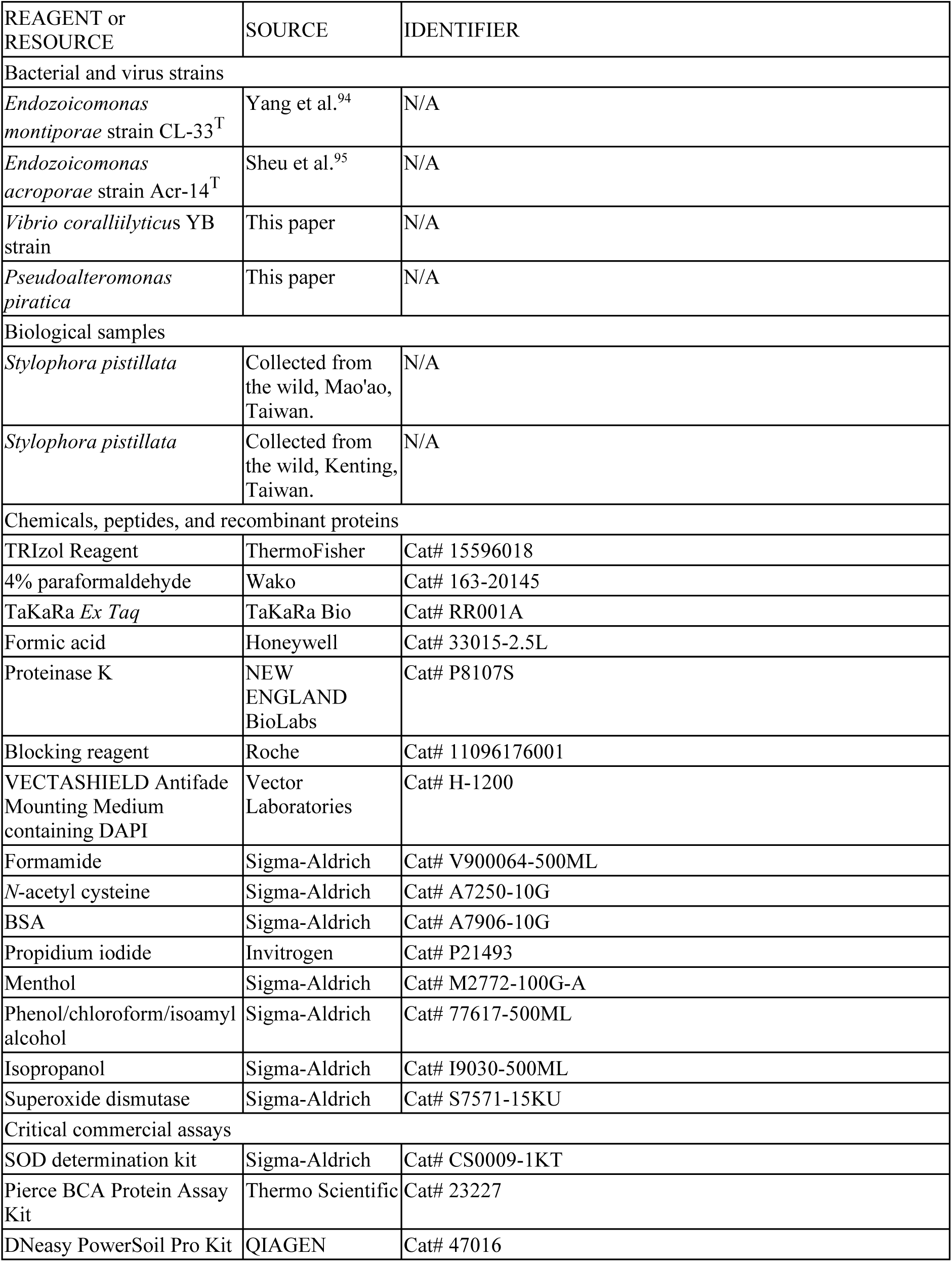
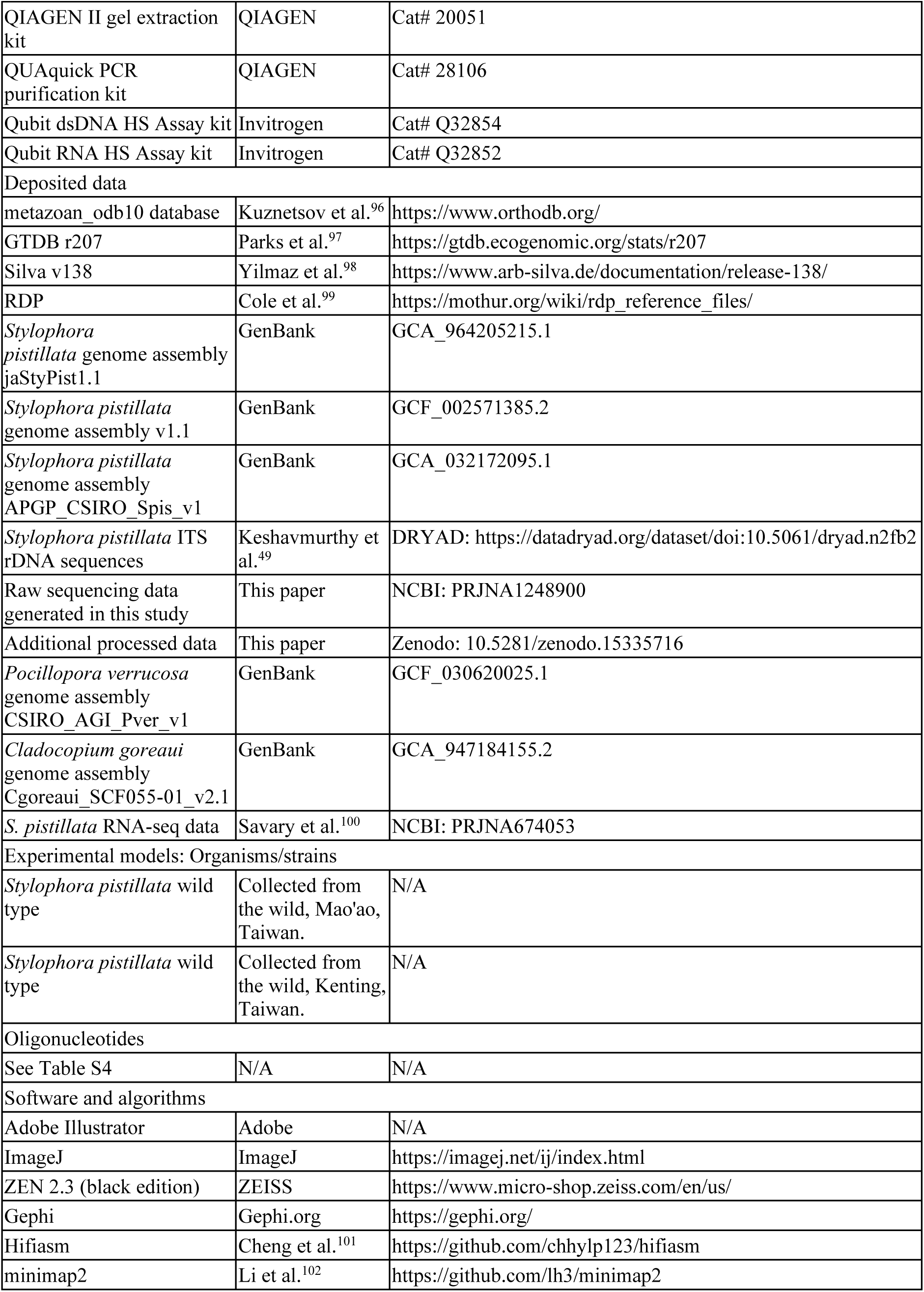
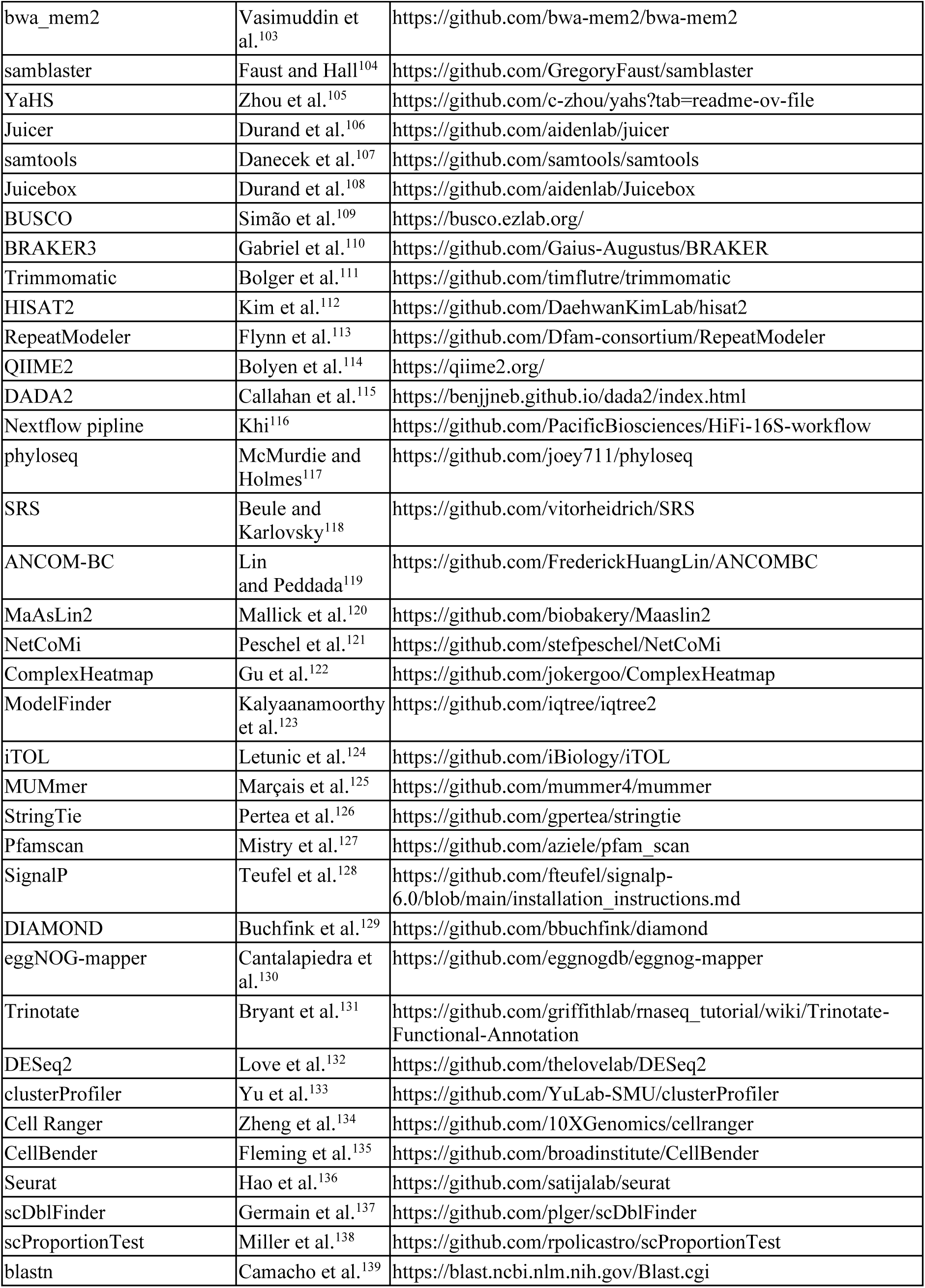
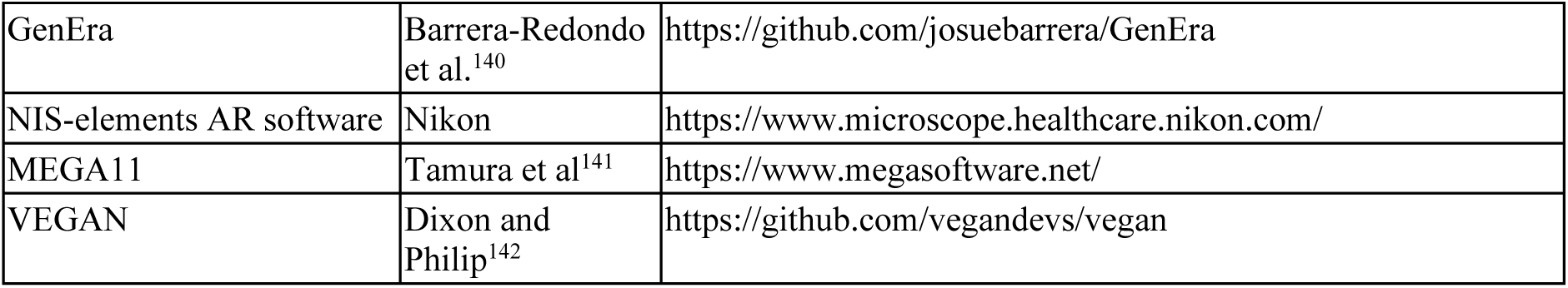

### EXPERIMENTAL MODEL AND SUBJECT DETAILS

#### Field work

All corals used in this study belong to *Stylophora pistillata* clade 1. Three coral bleaching experiments were conducted to serve different experimental purposes, including efficacy evaluation, geographic validation, and molecular profiling. For efficacy evaluation, 20 nubbins from five colonies (4 nubbins each) were collected at 10 m depth from Mao’ao, Taiwan (25.017513°N, 121.989937°E; Figure 1A) in September 2022 (seawater temperature: 26°C). For geographic validation, 40 nubbins from four colonies (10 nubbins each) were collected at 8 m depth from Kenting, Taiwan (21.930240°N, 120.745618°E; Figure 1A) in November 2022 (seawater temperature: 26°C). For molecular profiling, 80 nubbins from five colonies (16 nubbins each) were collected from the same location as geographic validation at depths of 7-10 m in June 2023 (seawater temperature: 28°C).

During sampling, only healthy colonies spaced at least 5 m apart were selected to avoid collecting clonal ramets.^143^ Nubbins approximately 3 cm in size were cut using bone cutters and placed in zip-loc bags. Nubbins from the first two experiments were transported to laboratory within one day. In molecular profiling, corals were initially transferred to the National Museum of Marine Biology and Aquarium and acclimated for two weeks in a 5,000 L in-door flow-through pond with natural sunlight and seawater maintained at 28°C. They were subsequently transported to the laboratory in iceboxes within one day.

For sequencing the *S. pistillata* clade 1 genome, a single colony was collected at 7 m depth from Kenting, Taiwan (21.930240°N, 120.745618°E; Figure 1A) in May 2023. The colony was maintained in a 150-L closed-system aquarium at laboratory prior to processing. Sampling was conducted under permits 1110006783 (Kenting) and 1110211109 (Mao’ao).

### Treatment set-up and sampling strategies

#### Efficacy evaluation experiment

After arrival at the laboratory, the 20 coral nubbins were distributed into four 2.5-L tanks (closed-system), with each tank containing five nubbins from five different colonies (n = 5; Figures S1A and S1C). Within each tank, water circulation was maintained by aeration through an air pump. Each tank represented a distinct treatment condition, including: (1) ambient temperature with 0.22 filtered artificial seawater (FASW, Red Sea Salt, Red Sea; 26°C FASW); (2) heat stress with FASW (31°C FASW); (3) heat stress with *Endozoicomonas acroporae* Acr-14^T^ (31°C *E. acroporae*);^95^ and (4) heat stress with *Endozoicomonas montiporae* CL-33^T^ (31°C *E. montiporae*).^94^ Initially, all corals were acclimated at 26°C (ambient temperature of the sampling site) for one week. Temperature in heat stress treatments was then increased by 1°C per day to 31°C, maintained at 31°C for six days, followed by a reduction of 1°C per day to 26°C (DHW = 3.4°C-weeks), and held at this temperature for a 10-day recovery period. The ambient temperature treatment remained constant at 26°C throughout the experiment (Figure S2C).

All nubbins were sampled at the end of the experiment (day 32) for 16S rRNA gene amplicon sequencing. Each nubbin was rinsed with FASW, wrapped in aluminum foil, snap-frozen in liquid nitrogen, and stored at -80°C until DNA extraction.

#### Geographic validation experiment

Ten 10-L closed-system aquaria, each consisting of a 2.5-L coral tank and a 7.5-L sump, were used in geographic validation. Water from the sump was circulated to the coral tank using a submersible pump (I-126, ISTA) and returned via a silicone tube, maintaining a flow rate of approximately 3.4 L/min (water turnover time: 44 s). Upon arrival, the 40 coral nubbins were evenly distributed among the ten aquaria, each receiving four nubbins originating from four different colonies (n = 4; Figures S1B and S1C). Five treatments were established, with each treatment consisted of two aquaria, one designated for sample collection and the other for measuring photosynthetic efficiency. Treatments included: (1) ambient temperature with FASW (26°C FASW); (2) ambient temperature with *E. acroporae* Acr-14^T^ (26°C *E. acroporae*); (3) heat stress with FASW (31°C FASW); (4) heat stress with *E. acroporae* Acr-14^T^ (31°C *E. acroporae*); and (5) heat stress with sterilized *E. acroporae* Acr-14^T^ (31°C sterilized *E. acroporae*).

All corals underwent a two-week acclimation at 26°C (ambient temperature of the sampling site). Temperature in heat stress treatments was then increased by 1°C per day to 31°C, maintained for six days, and then decreased by 1°C per day to 27°C (DHW = 2°C-weeks). Ambient temperature treatments remained constant at 26°C throughout the experiment (Figure S2D).

Coral samples for 16S rRNA gene amplicon sequencing were collected on the final day at 31°C (day 20). Each nubbin was rinsed with FASW, wrapped in aluminum foil, snap-frozen in liquid nitrogen, and stored at -80°C until DNA extraction.

#### Molecular profiling experiment

Twenty 10-L closed-system aquaria, identical to those used in geographic validation, were employed. The 80 coral nubbins were allocated evenly among the twenty aquaria, with each aquarium containing four nubbins from different colonies. Four treatments were established, each composing five independent aquaria, making individual aquaria replicates unit (n = 5, true replicate; Figures S1B and S1C). Each aquarium housed nubbins designated for collection at four specific sampling times. At each time point, five nubbins (each from different colonies) from five independent aquaria were collected per treatment.

The four treatments included: (1) ambient temperature with FASW (28°C FASW); (2) ambient temperature with *E. acroporae* Acr-14^T^ (28°C *E. acroporae*); (3) heat stress with FASW (31°C FASW); (4) heat stress with *E. acroporae* Acr-14^T^ (31°C *E. acroporae*). All corals underwent an initial two-week acclimation (in total, four weeks of acclimation) at 28°C (ambient temperature of the sampling site). Subsequently, the temperature in heat stress treatments was increased by 0.5°C per day until it reached 31°C, where it was maintained for 12 days, then decreased by 1°C per day to 28°C (DHW = 3.9°C-weeks), and maintained at this temperature for an additional 12-day recovery period. Ambient temperature treatments remained constant at 28°C throughout the experiment (Figure 1C).

The four sampling times occurred at: day 28 (post acclimation, T0); day 42 (during heat stress, T1); day 49 (end of heat stress, T2); and day 63 (post-recovery, day 63). At each sampling time, nubbins were rinsed with FASW. Each nubbin was then divided into three (for T0 and T1) or four fragments (for T2 and T3) using a bone cutter, each fragment designated for specific analyses: (i) 16S rRNA gene amplicon sequencing (all time points, n = 5); (ii) antioxidant enzyme activity measurement (all time points, n = 5); (iii) bulk RNA-seq (T1-T3 time points, n = 4); (iv) single-cell RNA-seq (T2 only, n = 1, pooled sample); and (v) fluorescence *in situ* hybridization (FISH, T3 only, n = 2) (Figure S1C).

Fragments intended for DNA extraction were wrapped in aluminum foil, snap-frozen in liquid nitrogen, and stored at -80°C. Those for antioxidant enzyme activity measurements were placed in zip-loc bags and kept on ice for further processing (see Method Details). Fragments for RNA extraction were placed in 2-mL cryotubes containing 1 mL TRIzol Reagent solution (ThermoFisher) and glass beads (0.5 mm diameter Glass beads, BioSpec), vortexed briefly, snap-frozen, and stored at -80°C. Fragments for single-cell RNA-seq were dissociated using calcium- and magnesium-free artificial seawater (CMFSW). Fragments for FISH analysis were fixed overnight in 4% paraformaldehyde (PFA, Wako) and subsequently stored in 70% ethanol at 4°C. Additionally, 500 mL of tank water from each treatment aquarium (n = 1) was collected, filtered through a 0.22 µm membrane, and stored at -20°C until DNA extraction.

### Temperature manipulation

Coral tank temperatures were maintained using water baths, with temperature monitored and controlled by Apex Neptune AquaControllers (Neptune Systems) connected to heaters (50-W heater, ISTA). A submersible pump (I-126, ISTA) was placed in individual water bath to ensure homogeneous temperature. For geographic validation and molecular profiling, an additional heater was placed in each sump. Each coral tank contained a thermometer (ISTA). All thermometers were calibrated using a calibrated thermometer (Pocket digital thermometer, DGS) prior to experiments. Chillers were not required, as room temperature was consistently maintained at 23°C.

### Culturing conditions and tank maintenance

All three coral bleaching experiments used artificial seawater (Red Sea Salt, Red Sea) for culturing. An artificial 12:12-hour light:dark cycle was established using LED aquarium lights (Illumagic). Light intensity was set to 150 μmol photons·m^−2^·s^−1^ during sunrise and sunset and gradually increased to 200 μmol photons·m^−2^·s^−1^ at midday, measured with an underwater quantum meter (MQ-510, Apogee). Freshly hatched *Artemia* (brine shrimp eggs, O.S.I.) were fed weekly to coral nubbins at a concentration of 20 individuals/mL for two hours after sunset (Figure S13), following recommended feeding frequencies to meet the nutritional needs of corals during bleaching experiments.^143^

Water parameters, including salinity (measured with a refractometer, ATAGO), ammonium, nitrate, phosphate, carbonate hardness, and calcium (measured using Salifert Test Kits) were regularly monitored (Table S6B). When required, parameters were adjusted with deionized water or sodium bicarbonate (Shimakyu) through gentle dosing. Water changes were conducted daily, replacing 100% (2.5 L) of water in efficacy evaluation and 20% (2 L) in geographic validation and molecular profiling. Aquarium air pumps (HX-700, HEXA) provided gentle aeration and water flow in each coral tank.

### Probiotic treatment preparation and inoculation

*E. acroporae* Acr-14^T^ and *E. montiporae* CL-33^T^ were originally isolated from *Acropora* sp.^95^ and *Montipora aequituberculata*,^94^ respectively, in Kenting, Taiwan. Both bacterial strains were cultured at 26°C with shaking at 200 rpm in Modified Marine Broth Version 4 (MMBV4) medium,^31^ supplemented with an additional carbon source (0.01% glucose for *E. acroporae* Acr-14^T^; 0.01% maltose for *E. montiporae* CL-33^T^). To prepare probiotic treatments, bacterial glycerol stocks were first revived for three days, followed by a one-day activation subculture and a final enrichment subculture in large volume for 20 hours. Cultures were harvested during exponential growth (optical density at 600 nm [OD_600_] = 0.73 ± 0.19, Table S6A), as determined from growth curves.

On inoculation day, bacterial cultures were centrifuged at 3,000 × *g* for 10 min at room temperature (RT) using 500-mL centrifuge bottles (Polypropylene Bottle, Beckmen). Supernatants were discarded, and cell pellets were resuspended in FASW to prepare concentrated probiotic solutions. Before probiotic addition, all submersible pumps in the sump were turned off to halt water circulation (not required in efficacy evaluation). Concentrated probiotic solution was then added to each coral tank to achieve a final OD_600_ of 0.1 (Table S6A), corresponding to 1.19 ± 0.64 × 10^7^ viable cells/mL for *E. acroporae* Acr-14^T^ and 1.16 ± 0.15 × 10^6^ viable cells/mL for *E. montiporae* CL-33^T^, as determined by colony-forming unit counts on agar plates. The same volume of FASW was added to control treatments as a placebo. After a 2-hour inoculation, coral nubbins were rinsed twice with FASW and transferred to clean coral tanks pre-filled with temperature-adjusted fresh ASW. Probiotic inoculations were conducted twice weekly, ensuring no overlap with sampling or feeding events. Detailed inoculation procedures are illustrated in Figure S13.

## METHOD DETAILS

### Monitoring coral bleaching responses

Pulse-amplitude-modulated (PAM) fluorometry (junior-PAM system, Walz GmbH) was used to measure the maximum quantum yield of photosystem II (Fv/Fm) in coral nubbins, with settings as follows: measuring light intensity = 6; gain = 1; saturation pulse intensity = 8; saturation pulse width = 0.6 s; damping = 2. After a 30-min dark acclimation period, three measurements were randomly taken per nubbin and averaged to obtain representative values. Measurements were consistently taken from the same nubbins throughout the experimental period. Differences in photosynthetic efficiency between treatments were analyzed by one-way ANOVA followed by Tukey’s post-hoc test.

For molecular profiling, at each sampling time point, collected nubbins were photographed in a 40 × 40 × 40 (cm) studio (DEEP) with two fixed light sources (5,500 K, LED), using a TG5 digital camera (OLYMPUS) at consistent camera settings. Nubbin coloration was assessed using the Coral Health Chart^144^ on a scale from D1 to D6. Differences in color scores between treatments were evaluated with paired-sample *t*-tests.

### Antioxidant enzyme activities

#### Sample processing

Catalase (CAT) and superoxide dismutase (SOD) activities were measured in the molecular profiling experiment. Coral tissue was airbrushed with ice-cold lysis buffer (50 mM phosphate, 0.1 mM EDTA, 10% [v/v] glycerol, pH 7.0). Samples were centrifuged at 2,000 × *g* for 5 min at 4°C to pellet Symbiodiniaceae cells. Supernatants used for coral host tissue were transferred to new Falcon tubes, centrifuged at 16,000 × *g* for 5 min at 4°C to remove cell debris, aliquoted, snap-frozen, and stored at -80°C until analysis. Symbiodiniaceae cell pellets were resuspended in lysis buffer and washed four times by centrifugation (2,000 × *g*, 5 min, 4°C). Cells were lysed using a bead mill (PowerLyzer 24 Homogenizer, Qiagen) in cryotubes containing 250 mg glass beads (0.5 mm, Biospec) at 3,000 rpm for 15 min, with cycles of 15 s beating followed by 30 min chilling on ice. After centrifugation at 16,000 × *g* for 5 min at 4°C, supernatants were aliquoted, snap-frozen, and stored at -80°C until analysis. We omitted one sample from the 28°C FASW treatment at T3 because the material was insufficient.

#### Antioxidant-enzyme-activity measurements

CAT activity was measured by mixing 10 µL of sample lysate with 740 µL of potassium phosphate buffer (50 mM, 0.1 mM EDTA, pH 7.0) in a quartz cuvette. The reaction was initiated by adding 50 µL of 320 mM H_2_O_2_ (final concentration of 20 mM) and monitored at 240 nm for 3 min at RT using a spectrophotometer (U-3900, Hitachi). Measurements were performed in triplicate, and enzyme activity was calculated using an extinction coefficient of 43.6 M^-1^cm^-1.145^

SOD activity was measured using an SOD determination kit (Sigma-Aldrich), following the manufacturer’s protocol. A standard curve was generated for each assay using serial dilutions of an SOD standard (Superoxide dismutase, Sigma-Aldrich). Samples were analyzed in triplicate in a 96-well plate, and absorbance was recorded at 450 nm using a microplate reader.

Total protein concentration for both coral hosts and Symbiodiniaceae samples were determined using a Pierce BCA Protein Assay Kit (Thermo Scientific) following the manufacturer’s protocol. CAT and SOD activities were normalized against the control treatment at each time point (28°C FASW). Statistical differences in enzyme activities among treatments were assessed using one-way ANOVA followed by Tukey’s post-hoc tests.

### 16S rRNA gene sequencing

#### DNA extraction

Frozen coral nubbins were thawed on ice, and tissue samples were collected using an airbrush filled with ice-cold 0.22 µm filtered TE buffer (1 mM EDTA, 10 mM Tris-HCl, pH 8.0). The resulting tissue slurry was transferred into 15-mL Falcon tubes and centrifuged at 12,000 × *g* for 10 min at 4°C to remove the supernatant. Genomic DNA from tissue pellets (or 0.22-µm membranes; tank water samples) was extracted using a DNeasy PowerSoil Pro Kit (QIAGEN) following the manufacturer’s protocol. DNA quality was verified by agarose gel electrophoresis and quantified using a NanoDrop 1000 spectrophotometer (ThermoFisher).

#### Short-length 16S rRNA gene sequencing

16S rRNA gene amplicon sequencing was conducted for samples collected during efficacy evaluation (20 coral samples), geographic validation (20 coral samples), and molecular profiling (79 coral and 16 tank water samples). Additionally, six blank samples (DNA extracted from TE buffer) were included to assess potential contamination. The bacterial V6-V8 hypervariable region of the 16S rRNA gene was amplified using the universal primers 968F (5′- AACGCGAAGAACCTTAC-3′)^146^ and 1391R (5′-ACGGGCGGTGWGTRC-3′).^147^ Each PCR reaction was performed in a total volume of 100 µL containing 1.5 U TaKaRa *Ex Taq* (TaKaRa Bio), 1 x TaKaRa *Ex Taq* buffer, 0.2 mM dNTP mix, 0.2 µM each of forward and reverse primers, template DNA, and nuclease-free water. PCR conditions were the same as a previous work,^148^ which consisted of an initial denaturation at 94°C for 5 min, followed by 30 cycles of denaturation at 94°C for 30 s, annealing at 52°C for 20 s, extension at 72°C for 45 s, and a final extension step at 72°C for 10 min. PCR products were verified by 1.5% agarose gel electrophoresis, and the target amplicon (424 bp) was excised and purified using the QIAGEN II gel extraction kit (QIAGEN). Barcode-tagging PCR (five cycles) was conducted and DNA was quantified using a NanoDrop 1000 (ThermoFisher) and Qubit HS DNA kit (Qubit Fluorometer, Invitrogen). Samples were pooled in equal quantities, further purified using the QUAquick PCR purification kit (QIAGEN), and sequenced using an Illumina MiSeq platform (300 bp paired-end chemistry) by Yourgene Biosciences (Taiwan).

#### Full-length 16S rRNA gene sequencing

Molecular profiling DNA samples were sent to Blossom Biotechnologies (Taiwan) for PCR amplification, library preparation, and sequencing. The protocol followed PacBio guidelines with minor modifications in PCR cycle number and annealing temperature. Briefly, full-length 16S rRNA genes were amplified using the universal primers 27F (5′- AGRGTTYGATYMTGGCTCAG-3′) and 1492R (5′-RGYTACCTTGTTACGACTT-3′),^149^ each tagged with unique PacBio barcodes. PCR was performed using KAPA HiFi HotStart ReadyMix (KAPA Biosystems), under the following conditions: initial denaturation at 95°C for 3 min; 30 cycles of denaturation at 95°C for 30 s, annealing at 55°C for 30 s, extension at 72°C for 60 s. Amplified DNA concentration and size were assessed using the Qubit HS DNA kit (Qubit Fluorometer, Invitrogen) and Fragment Analyzer (Agilent Technologies). Amplicons were pooled in equal concentrations, libraries were prepared using the SMRTbell prep kit 3.0, and sequencing was performed on the PacBio Sequel IIe platform.

### Fluorescence *in situ* hybridization (FISH)

#### Testing probe specificity

The Endo-Group B probe^39^ was used to detect *Endozoicomonas* signals in samples. Probe specificity to *E. acroporae* Acr-14^T^ was verified using the web-based tool TestProbe3.0 (https://www.arb-silva.de/search/testprobe/) with the Silva SSU r138.2 database. Additionally, the specificity of the Endo-Group B probe was experimentally validated with three bacterial cultures: *E. acroporae* Acr-14^T^ (positive control), *Vibrio coralliilyticus* YB strain, and *Pseudoalteromonas piratica* (both negative controls). Cultures were grown in MMBV4 medium, fixed with 4% PFA (Wako), and individually applied to microscope slides, which were then dried completely at 40°C.

QuickHCR-FISH was performed according to a previously described method.^150^ Slides were initially rinsed with 20 mM Tris-HCl (pH 8.0) for 10 min and permeabilized with 1 µg/mL proteinase K in 20 mM Tris-HCl at 37°C for 5 min. Slides were rinsed again with 20 mM Tris-HCl (pH 8.0) for 10 min and prehybridized in hybridization buffer (20 mM Tris-HCl [pH 8.0], 0.9M NaCl, 0.01% SDS, 10% dextran sulfate, 1% blocking reagent, and 25% formamide) for 5 min. For hybridization, cells were incubated first with Non338-initiatorC probe^40^ at 46°C for 1 h, followed by incubation with EUB338mix-initiatorH^38^ and Endo-Group B-initiatorR probes at 46°C for 2 h. Cells were washed using washing buffer (20 mM Tris-HCl [pH 8.0], 0.149M NaCl, 0.5mM EDTA, and 0.01% SDS).

For signal amplification, Cy3-labeled H1 and H2 amplifier probes, Cy5-labeled R1 and R2 amplifier probes, and Alexa 488-labeled C1 and C2 amplifier probes were individually prepared in amplification buffer (50 mM Na2HPO4, 0.9 M NaCl, 0.01% SDS, 10% dextran sulfate, and 1% blocking reagent), incubated at 95°C for 1.5 min, cooled at 25°C for 30 min, and mixed. Slides were pre-immersed in amplification buffer for 5 min, and then incubated with mixed amplifier probes at 35°C for 30 min. Slides were subsequently rinsed with PBS at 4°C for 10 min, washed with ddH_2_O, dehydrated with 100% ethanol, air dried, mounted in VECTASHIELD Antifade Mounting Medium containing DAPI (Vector Laboratories), covered with coverslips, and stored at 4°C until observation.

#### FISH on coral samples

Eight coral fragments (two from each treatment, n = 2) were collected at T3 during molecular profiling. Samples fixed with PFA were decalcified using Morse’s solution (10% sodium citrate and 22.5% formic acid), exchanging the solution twice over three days at 4°C. Samples were rinsed twice with 10 mM PBS and stored in 70% ethanol at 4°C. Tissue dehydration was performed using a Scientific Excelsior ES Tissue Processor (Thermo Scientific) and embedded in paraffin. Five microscope slides, each containing three 5-µm serial sections, were prepared per sample (40 slides total) using a Lecia RM2125 RTS microtome (Leica). A 50-µm thickness interval between slides was maintained to observe more tissue regions. Prior to QuickHCR-FISH, tissue sections were dewaxed in Xylene (two cycles of 15 min each), dehydrated in 100% ethanol (two cycles of 5 min each), and air-dried. QuickHCR-FISH was then performed as described above.

#### Microscope observation

Slides were observed with a 40x objective lens on a fluorescence microscope (Ni-L, Nikon) with NIS-elements AR software. The area of *E. acroporae* Acr-14 signals in tissue sections was calculated by measuring overlapping Cy3 and Cy5 signals (with no Alexa 488) using ImageJ software (version 2.14.0/1.54f). High-resolution images were obtained using a 63x objective lens on a confocal laser scanning microscope (Zeiss LSM 880) with ZEN 2.3 (black edition) software. Excitation/emission wavelengths were as follows: DAPI and autofluorescence (405 nm excitation, 417–500 nm emission), Alexa 488 (488 nm excitation, 501–543 nm emission), Cy3 (561 nm excitation, 565–641 nm emission), and Cy5 (633 nm excitation, 652–735 nm emission). Results from bacterial cultures confirmed Endo-Group B probe specificity, detecting signals exclusively in *E. acroporae* Acr-14^T^ and not in negative control bacteria (Figure S4D).

### RNA extraction and bulk RNA-seq

Coral samples were thawed on ice and homogenized for 1 min at 2,000 rpm using a homogenizer (PowerLyzer 24, Qiagen). Homogenates were centrifuged at 12,000 × *g* for 10 min at 4°C and supernatant was collected. Total RNA extraction was performed following the TRIzol reagent protocol. Extracted RNA was aliquoted and stored at -80°C. Prior to sequencing, RNA quality and quantity were assessed using a NanoDrop 1000 (ThermoFisher) and the Qubit HS RNA kit (Qubit Fluorometer, Invitrogen). RNA integrity was evaluated for all samples using a Fragment Analyzer (Agilent Technologies). In total, 51 libraries were constructed with stranded Poly-A protocol. Libraries were sequenced on the NextSeq 2000 sequencing system (Illumina) with 100 bp paired-end reads.

### Dissociation and single-cell RNA-seq

Conducting time-course single-cell experiments in corals presents significant technical challenges due to the absence of protocols for long-term preservation of dissociated single cells. Standard fixation methods are incompatible with droplet-based platforms. To overcome this, we developed a fixation-compatible workflow optimized for 10x Genomics.

#### Coral cell dissociation

At T2 of molecular profiling, three coral fragments (approximately 0.5–1 cm each) were collected from each treatment group. Each fragment was immersed in 3 mL of calcium- and magnesium-free seawater supplemented with 2% *N*-acetyl cysteine (NAC-CMFSW) in a 24-well plate for 5 min. This step, designed to reduce mucus and prevent interference with downstream cell dissociation, was repeated once. Fragments were then immersed in CMFSW alone for an additional 5 min (total immersion time: 15 min). Following this pre-treatment, fragments were transferred to a fresh well containing 2 mL of CMFSW, and cell dissociation was performed by gentle pipetting using a 1-mL pipette. All dissociation steps were completed within 30 min per sample. To assess cell viability, an aliquot of the cell suspension was stained with 2 µg/mL propidium iodide, a dye that selectively stains dead cells with compromised plasma membranes. Cell survival rate was examined using a hemocytometer under a fluorescence microscope (Ni-L, Nikon). All samples exhibited high viability, with a mean survival rate of 91 ± 1.15% (mean ± SD).

#### Acetic acid and methanol (ACME) fixation and storage

Cell suspensions from the three fragments in each treatment were pooled, resulting in one combined suspension derived from three different coral colonies. A modified ACME fixation protocol was then applied to each pooled sample.^151^ Briefly, 2 mL of the pooled cell suspension was fixed by adding 8 mL of freshly prepared ACME solution (methanol:acetic acid:glycerol:7.5% NAC:PBS at a 15:10:10:2:43 ratio) in a 15-mL Falcon tube. Samples were incubated at room temperature for 20 min on a shaker set at 45 rpm. After fixation, cells were passed through a 40-µm cell strainer (pluriSelect) to remove cell clusters and to ensure a uniform single-cell suspension, before being centrifuged at 1,200 × *g* for 5 min at 4°C. To remove the supernatant without disturbing the pellet, most of the liquid was first aspirated using a 1-mL pipette, followed by a 200-µL pipette for the final volume approaching the cell pellet. The cell pellet was then washed with 10 mL of PBS containing 3% BSA (Sigma-Aldrich). This step is critical for trapping dead cells and debris, thereby improving the overall quality of the suspension. After a second centrifugation under the same conditions, the supernatant was discarded using the same pipetting method, and cells were resuspended in 1 mL of PBS containing 3% BSA and 10% dimethyl sulfoxide (DMSO). DMSO served as a cryoprotectant to prevent cell damage during freezing. Before storage, each fixed suspension was aliquoted into two Lobind tubes (Eppendorf): one for cell counting and viability assessment, and the other for downstream 10x Genomics single-cell sequencing. All samples were stored at -80°C. All reagents were filtered through 0.22 µm membranes prior to use.

#### Single-cell library preparation and RNA sequencing

Prior to library preparation, fixed cell suspensions were thawed on ice, and the storage solution was replaced with PBS by centrifugation. An aliquot from each cell suspension was stained with Trypan Blue, and cell counts were preformed using an EVOS microscope (Invitrogen) with a 10x objective lens and a C-Chip hemocytometer (NanoEntek). Approximately 10,000 cells from each sample were used for single-cell library preparation with the Chromium Single Cell 3’ Reagent Kit v3, following the manufacturer’s protocol. Four libraries, corresponding to the four experimental treatments, were generated and sequenced on an Illumina NextSeq 2000 platform using a P3 flow cell (100-cycle kit, 1.0 Cells configuration).

### *S. pistillata* chromosome-level genome sequencing

#### Algal symbiont removal

Obtaining uncontaminated, high-molecular-weight DNA from coral tissue is challenging due to the presence of intracellular symbiotic algae. To overcome this, we applied menthol treatment to induce bleaching^152^ before DNA extraction, thereby minimizing algal contamination.^153^ Briefly, coral nubbins were maintained in a 150-L aquarium system. Menthol treatment was conducted in a separate 2-L tank equipped with an air pump to provide aeration and water circulation. Coral nubbins were exposed to 0.2 mM menthol in ASW for no more than 8 h per day, depending on coral condition, and then returned to the main aquarium for recovery. After one week, these nubbins appeared completely bleached with no detectable Symbiodiniaceae under a stereomicroscope (Leica; Figure S6B).

#### DNA extraction and sequencing

To extract total genomic DNA from *S. pistillata*, bleached coral nubbins were rinsed with FASW, cut into 0.5 cm fragments, and incubated in 1-mL chaotropic lysis buffer (4.3 M guanidine thiocyanate, 17 mM *N*-Lauroylsarcosine sodium salt, and 25 mM Tris-HCl) in a 2-mL centrifuge tube. Samples were then shaken at 100 rpm and incubated at 55°C for 2 days. Subsequently, 250 µL of the solution was transferred to a new centrifuge tube, mixed with 250 µL PEB buffer (100 mM Tris-HCl [pH 8.0], 10 mM EDTA, and 0.1% SDS), and then mixed thoroughly with 250 µL Phenol/chloroform/isoamyl alcohol (PCI) solution. After centrifuging at 15,000 × *g* for 10 min, the supernatant was transferred into a new tube, mixed with 500 µL chloroform/isoamyl alcohol, and centrifuged again at 15,000 × *g* for 10 min. This chloroform wash step was repeated once. Approximately 400 µL of supernatant was then collected, mixed with 40 µL 3 M sodium acetate and 280 µL isopropanol, and centrifuged at 15,000 × *g* for 20 min at 4°C. The DNA pellet was washed twice with 1 mL ice-cold 70% ethanol, centrifuged at 15,000 × *g* for 10 min at 4°C, air-dried, and finally dissolved in TE buffer. Extracted DNA was aliquoted and stored at -20°C. DNA quality and concentration were assessed using a NanoDrop 1000 (ThermoFisher) and a Qubit HS DNA kit (Qubit Fluorometer, Invitrogen). A whole-genome library was prepared using the SMRTbell Prep Kit 3.0 and sequenced on a PacBio Sequel IIe platform.

#### Hi-C sequencing

For Hi-C library preparation, we applied a low-speed centrifuge approach to reduce Symbiodiniaceae contamination in the sample. Briefly, coral cells were dissociated from healthy coral fragments using CMFSW as described above. This suspension was centrifuged at 100 × *g* for 5 min at 4°C, and the supernatant was collected, filtered through a 40-µm cell strainer, and centrifuged again under the same conditions. The resulting supernatant was centrifuged at 1,300 × *g* for 5 min at 4°C to pellet cells, which were then snap-frozen in liquid nitrogen and stored at -80°C. This method reduced the proportion of Symbiodiniaceae from 29.1% to 3.3%, as verified using a hemocytometer. The procedure was repeated until approximately 6 × 10^7^ coral cells were obtained. The Hi-C proximity ligation library was prepared with all collected cells using the Proximo Hi-C (Animal) Kit (Phase Genomics) following the manufacturer’s instructions, and sequencing was performed using a NextSeq 2000 sequencing platform with 150-bp paired-end reads.

### *S. pistillata* chromosome-level genome assembly

#### Primary assembly

The PacBio Sequel IIe platform yielded 2.1 million HiFi reads, totaling 31.7 Gb of sequencing data, with a mean read length of 18 kb, corresponding to approximately 69-fold genome coverage. Reads were assembled de novo using Hifiasm (v0.19.9-r616)^101,154,155^ without incorporating Hi-C data, which generated 194 scaffolds with a total size of approximately 492.9 Mb and a N50 scaffold length of 26.9 Mb. Primary contigs were extracted by converting the GFA output to FASTA format. To remove residual haplotigs and erroneous duplications, we used the Purge_Dups (v1.2.5) pipeline. Briefly, PacBio HiFi reads were aligned to the Hifiasm primary contigs using minimap2 (v2.28-r1209).^102^ Sequencing coverage was estimated by pbcstat and calcuts, generating coverage cutoffs for downstream analysis. Self-alignment of the assembly was performed, followed by the purge of haplotigs based on depth of coverage and overlap. Purged primary contigs were extracted, resulting in a haplotype-purged assembly comprising 59 scaffolds with a total size of 471.2 Mb.

#### Hi-C based scaffolding

Hi-C Illumina sequencing yielded 166.7 million read pairs, with 50.3 Gb read bases. Hi-C reads were aligned to the scaffold purged contigs using bwa_mem2^103^ and duplicates were marked with samblaster (v0.1.26).^104^ The resulting BAM file was used as input for YaHS (v1.2a.1),^105^ which utilizes Hi-C linkage information to scaffold contigs into chromosome-scale scaffolds. This step produced a draft chromosome-level assembly that was then used to generate Hi-C contact maps.

#### Assembly curation and final polishing

A Hi-C contact matrix was generated using the Juicer^106^ pipeline based on read alignments. Chromosome size information was obtained from the final scaffolds using samtools (v1.21).^107^ The draft assembly and corresponding Hi-C contact map were loaded into Juicebox (v1.11.08)^108^ for manual curation. Scaffolding corrections were recorded, and an updated AGP file was generated using Juicer post-processing tools. Following manual curation, the final chromosome-level assembly totaled 471.2 Mb, with 97.15% of the sequence anchored into 14 chromosome-scale pseudomolecules. The scaffold N50 was 31 Mb. Assembly completeness was evaluated using BUSCO (v5.7.1)^109^ with the metazoan_odb10 database^96^, which identified 96.0% of complete BUSCO genes.

#### Gene model construction

Gene prediction was performed using BRAKER3 (v3.0.8),^110,156,157,158,159,160,161,162,163,164,165,166,167^ which integrates GeneMark-ES/SP and AUGUSTUS for ab initio gene prediction. Bulk RNA-seq datasets were trimmed with Trimmomatic (v0.39),^111^ and post-trimming read quality was assessed using FastQC (v0.12.1). Cleaned short reads were then mapped to the genome assembly using HISAT2 (v2.2.1).^112^ To account for repetitive elements, RepeatModeler (v2.0.6)^113^ was used to construct a de novo repeat library, and the genome was subsequently soft-masked using RepeatMasker (v4.1.5). Additionally, the metazoan_odb10 database^96^ was downloaded. The resulting BAM file, soft-masked genome, and metazoan protein database were used as inputs for BRAKER3. The final annotation contained 25,918 genes encoding 28,789 transcripts. The completeness of predicted gene models was evaluated using BUSCO (v5.7.1)^109^ with the metazoan_odb10 dataset, yielding a 97.0% BUSCO completeness (single: 84.0%, duplicated: 13.0%, fragmentated: 0.9%, missing: 2.1%).

## QUANTIFICATION AND STATISTICAL ANALYSIS

### Microbiome analysis

#### Short-length 16S data processing

Amplicon sequencing data generated by the Illumina MiSeq platform were processed using QIIME2 (v2021.8.0).^114^ Raw reads were reoriented, primers trimmed, and sequences demultiplexed using the cutadapt plugin (v3.4).^168^ Forward and reverse reads were truncated at quality score threshold of 25 and denoised using the DADA2 algorithm.^115^ Taxonomy of ASVs was assigned using the classifier-consensus-vsearch plugin^169^ against the SILVA v138 SSU NR99 reference database.^98,170^

#### Full-length 16S data processing

PacBio full-length 16S sequencing data were analyzed using the Nextflow pipeline (v0.7).^116^ Briefly, HiFi reads with a Q20 quality threshold were trimmed, and primers were oriented using cutadapt.^168^ ASVs were obtained by denoising using DADA2^115^ in QIIME2.^114^ Taxonomic classification was performed using VSEARCH against GTDB r207,^97^ Silva v138,^98,170^ and RefSeq + RDP^99^ databases. No minimum frequency or sample filter was applied, and the minimum sequence length retained was set to 900 bp.

#### Data filtering and decontamination

ASVs assigned to chloroplast, mitochondria, eukaryota, archaea, or unclassified kingdoms and phyla were removed, as were singleton ASVs. Decontamination was performed by removing ASVs whose averaged reads counts were higher in blanks than in formal samples. In total, 34 ASVs and 37 ASVs were identified as background microbial signals in the short-length and full-length 16S datasets, respectively (Table S1B and S1C). After filtering and decontamination, 668, 1,371, 2,861, and 3,027 ASVs remained, with average read counts of 46,526 (efficacy evaluation), 36,115 (geographic validation), 22,154 (short-length, molecular profiling), and 6,266 (full-length, molecular profiling).

#### Community profiling

Microbial relative abundance profiles at the class level were visualized using phyloseq^117^ and ggplot2.^171^ To examine probiotic strains in corals, sequences assigned to *Endozoicomonas* were searched against a custom database build from the genomes of *E. acroporae* Acr-14^T^ (JBQGQA000000000) and *E. montiporae* CL-33^T^ (GCF_001583435.1) using BLASTn with an e-value of 0 and an identity threshold of ≥ 99.5%. This step was omitted for short-length molecular profiling data, as full-length sequencing provided species-level resolution and confirmed all *Endozoicomonas* ASVs as *E. acroporae* Acr-14^T^.

#### Diversity analysis

Rarefaction curves were generated using the “ggrare” function in phyloseq-extended package. For short-length data, a sample from the 31°C FASW treatment at T1 was excluded from downstream analysis due to low read count (< 1,000 reads). Further analyses were not performed on full-length data due to low sequencing depth (< 1,000 reads for approximately one-third of samples) (Figure S3H), primarily caused by abundant chloroplast sequences in coral samples.

Alpha diversity metrics, including AVS richness and Shannon index, were calculated using the microbiome package after rarifying samples to an even depth (efficacy evaluation: 18,749 reads; geographic validation: 16,905 reads; molecular profiling: 1,694 reads) using the SRS package.^118^ Differences between treatments were assessed using Wilcoxon signed-rank tests. Multivariate analyses were performed at the ASV level using relative-abundance-transformed data. Bray-Curtis distances between treatments were calculated using vegan^142^ and visualized with principal coordinates analysis (PCoA). Microbial community differences between treatments were assessed by permutational multivariate analysis of variance (PERMANOVA) with 999 permutations using the “adonis” function. Multivariate dispersion of treatments was evaluated with “betadisper” and tested using ANOVA.

#### Differential abundance analysis

Differential abundance analyses at the genus level employed a consensus approach using ANCOM-BC^119^ and MaAsLin2^120^ for robust interpretation.^172^ The control treatment (ambient temperature with FASW) was used as the reference. ANCOM-BC was executed with parameters: “tax_level” set to genus, “conserve” set to TRUE, and “prv_cut” at 0.05. For MaAsLin2, data were agglomerated at the genus level, transformed to relative abundances, and analyzed using default parameters. Only genera significantly different by both methods (*p* < 0.05) were accepted. Genera with low relative abundance (< 0.1% across all treatments within each experiment) were excluded. Results highlighting genera potentially contributing to probiotic effects were visualized with Venn diagrams and ComplexHeatmap (v2.18.0).^122^

#### Association network analysis

Microbial association networks at the genus level were constructed using NetCoMi (v1.1.0).^121^ To mitigate data sparsity, the top 60 genera with the highest prevalence across samples were retained using the “filtTax” and “filtTaxPar” functions. Data were normalized via centered log-ratio (clr) transformation. Genera associations were measured using SparCC correlations, and filtered with a threshold of 0.3. Clusters were identified through greedy optimization of modularity, and the resulting network was visualized using Gephi (v0.10.1). All analyses for microbiome data were conducted in R (v4.3.2) and colorblind safe color schemes were utilized for figures.

### *S. pistillata* clade 1 and clade 4 genome comparison

#### Genetic identity based on ITS rDNA

To confirm the genetic identity of these samples, we amplified the ITS1-5.8S-ITS2 region using the anthozoan-universal primer pairs 1S (5′-GGTACCCTTTGTACACACCGCCCGTCGCT-3′) and 2SS (5′-GCTTTGGGCGGCAGTCCCAAGCAACCCGACTC-3′).^173^ PCR conditions were performed as follows: initial denaturation at 95°C for 4 min; 30 cycles of denaturation at 94°C for 30 s, annealing at 55°C for 60 s, extension at 72°C for 2 min; and a final extension at 72°C for 10 min. PCR products were summited to Genomics Biotechnologies (Taiwan) for Sanger sequencing. Additionally, an *S. pistillata* genome recently released in NCBI without geographic origin information (GCA_964205215.1) was included by extracting its ITS sequence. Obtained sequences were aligned with 172 publicly available *S. pistillata* ITS rDNA sequences^49^ using MEGA11.^141^ Phylogenetic analysis of the 174 ITS sequences was performed using IQ-TREE (v2.0.6) with ModelFinder enabled^123^ and 1,000 bootstrap replicates. The resulting treefile was imported into iTOL (v7.0)^124^ to construct a phylogenic tree rooted with an outgroup. As expected, *S. pistillata* samples used in this study clustered with clade 1, whereas the GCA_964205215.1 sequence clustered with clade 4, suggesting its origin from the Red Sea-Persian/Arabian Gulf-Kenya region (Figure S6A).

#### Pairwise whole-genome alignments

To investigate genomic conservation and divergence between *S. pistillata* clades 1 and 4, pairwise whole-genome alignments were conducted using genomes from: GCA_964205215.1 (*S. pistillata*, location unknown); GCF_002571385.2 (*S. pistillata*, Red Sea); GCA_032172095.1 (*S. pistillata*, Australia); GCF_036669915.1 (*Pocillopora verrucosa*, Spratly Islands); GCA_014529365.1 (*P. verrucosa*, Red Sea); GCF_030620025.1 (*P. verrucosa*, Australia); and the genome assembled in this study (*S. pistillata*, Taiwan). Genome scaffold lengths were determined using samtools (v1.21).^107^ Scaffolds shorter than 1,000 bp were excluded from the analysis. Pairwise alignments were performed using MUMmer (v4.0.0).^125^ One-to-one alignments were filtered using the delta-filter function, and alignment coordinates were extracted with show-coords. Based on these coordinates, average nucleotide identity (ANI) was calculated as followed: ANI = ∑ (ID%*Length of Alignment)/∑ (Length of Alignment). The alignment fraction (AF) was calculated as: AF = ∑ (Length of Alignment)/ (Length of the Query Genome). These metrics were adapted from previously described methods.^174^ Results were visualized in R using the scatterpie package.

#### Comparative alignment rates

RNA-seq data from *S. pistillata* samples from Taiwan (this study) and the Red Sea^100^ were used to compare alignment rate against the clade 1 (this study) and clade 4 (GCA_964205215.1) genomes as references. Reference genome indices were built using HISAT2 (v2.2.1),^112^ incorporating exon and splice-site annotations from their respective gene models. Trimmed bulk RNA-seq reads, with or without removal of Symbiodiniaceae sequences (*Cladocopium goreaui* for clade 1, GCA_947184155.2; *Symbiodinium microadriaticum* for clade 4, GCA_001939145.1), were then aligned to each reference genome index.

### Bulk RNA-seq data analysis

#### Mapping reads to host genome

Demultiplexing was performed by the sequencing facility. Adapter sequences and low-quality bases were trimmed using Trimmomatic (v0.39)^111^ and checked with FastQC (v0.12.1). We followed recommended best practices for transcriptome analysis, employing highly accurate tools.^175^ A genome index for *S. pistillata* clade 1 was built as described above, and trimmed reads were aligned using HISAT2 (v2.2.1).^112^ Unaligned reads were saved for downstream analysis of the Symbiodiniaceae transcriptome. Across 51 samples, alignment rates to the host genome ranged from 58.42% to 92.52%, with bleached samples showing notably higher alignment rates. SAM files were converted to compressed BAM format and sorted using samtools (v1.21).^107^

#### Mapping reads to the Symbiodiniaceae genome

To identify Symbiodiniaceae lineages present in corals from molecular profiling, the *LSU* region was amplified from sample DNA using primers 28S-forward (5′-CCCGCTGAATTTAAGCATATAAGTAAGCGG-3′) and 28S-reverse (5′-GTTAGACTCCTTGGTCCGTGTTTCAAGA-3′)^176^ with the following PCR conditions: initial denaturation at 90°C for 2 min, followed by 35 cycles of 94°C for 1 min, 60°C for 1 min, 72°C for 1 min, and a final extension at 72°C for 5 min. PCR products were submitted to Genomics Biotechnologies (Taiwan) for Sanger sequencing. BLAST searches against the NCBI core nucleotide database indicated that all five coral colonies were predominantly colonized by *Cladocopium sp.* (100% query coverage, 99.23% identity). The *Cladocopium goreaui* genome (GCA_947184155.2) was downloaded from NCBI. Read alignment was performed as for the host while using the *C. goreaui* genome and gene models to build the reference index. Alignment rates using host-unaligned reads ranged from 42.4% to 75.7%, except for two severely bleached samples (9.7% and 1.3% alignment), which were excluded from further analyses.

#### Estimating transcript abundance

Sorted BAM files were assembled using StringTie (v2.2.3),^126^ guided by BRAKER-derived gene models. Assemblies were merged into a unified transcriptome annotation using the --merge function, resulting in 41,541 genes identified in the *S. pistillata* transcriptomes. Transcript abundance per sample was estimated using StringTie (v2.2.3),^126^ with the *S. pistillata* merged annotation as the reference for host transcripts and the *C. goreaui* gene models as the reference for symbiont transcripts.

#### Transcriptome functional annotation

Transcript sequences were extracted from the genome using gffread (v0.12.7) based on the merged annotation. Open reading frames (ORFs) were predicted using TransDecoder (v5.7.0) with default parameters. Predicted protein functions were annotated using the following approaches: (i) Pfam domain identification using Pfamscan and the Pfam database (v37.0);^127^ (ii) signal peptide prediction using SignalP (v6.0)_;_^128^ (iii) gene names assignment via DIAMOND aligner (v2.1.10)^129^ against the human genome (vGRCh38, Ensembl release 113) with an e-value threshold of 1e-5; (iv) gene ontologies annotation using eggNOG-mapper (v2.1.12)^130^ with the eggNOG database (v5.0.2);^177^ and (v) comprehensive transcriptome annotation using Trinotate (v4.0.2).^131^ For the Symbiodiniaceae transcriptome, only methods (iv) and (v) were performed.

#### Differential expression analysis

Transcript abundance estimates were imported into R with tximport,^178^ utilizing a tx2gene table linking transcripts to genes. Differential gene expression analysis was performed at the gene level using DESeq2.^132^ Pre-filtering was performed to remove genes with fewer than 10 counts in the smallest group size (n = 3 in the 28°C *E. acroporae* treatment at T2), leaving 30,708 of 41,541 genes for the coral host and 25,130 of 40,390 genes for Symbiodiniaceae. Comparisons were made at each time point against the control (28°C FASW) or directly between the 31°C FASW and 31°C *E. acroporae* treatments. Genes with adjusted *p* < 0.05 were considered differentially expressed (DEGs). Results were visualized with the EnhancedVolcano package.

Expression profiles were explored using principal component analysis (PCA) plots generated from variance-stabilized (*vst*) normalized counts in DESeq2 and visualized with ggplot2.^171^ Upregulated and downregulated DEGs from each comparison were illustrated using Venn diagrams at each sampling time point, created with the VennDiagram package. Expression levels of selected genes were visualized using ComplexHeatmap (v2.18.0)^122^ with data normalized by z-scores.

#### Functional analysis

Gene Ontology (GO) enrichment analysis was performed using clusterProfiler (enricher function).^133^ GO terms and gene information was prepared from the annotated transcriptome, with all annotated, pre-filtered genes serving as the background. Terms significantly enriched (adjusted *p* value < 0.05 and *p* value < 0.0005) were visualized with ggplot2,^171^ presenting the top 10 (sort by adjusted *p* value) biological processes. Terms redundant with highly similarity, e.g., response to topologically incorrect protein vs cellular response to topologically incorrect protein, or irrelevant to coral biology, e.g., wing disc morphogenesis, were excluded.

Gene set enrichment analysis (GSEA) was conducted using clusterProfiler (GSEA function with fgsea algorithm).^133^ Genes were pre-ranked using the Wald statistic from DESeq2 results, and enrichments were considered significant at adjusted *p* < 0.05. GSEA results were visualized using ggplot2 and enrichplot packages.

### scRNA-seq data analysis

#### Generating count matrix and data filtering

Demultiplexing was performed by the sequencing facility. Cell Ranger (v8.0.1)^134^ was used for sequence alignment using the cellranger count function with –expect-cells set to 10,000 for all four libraries. The reference was generated from *S. pistillata* clade 1 genome and the StringTie-derived transcriptome annotation with genes lacking strand information removed. For the four libraries, sequences aligned to the host transcriptome captured 8,855, 9,219, 10,015, and 8,626 cells, respectively, corresponding to four treatments: 28°C FASW, 28°C *E. acroporae*, 31°C FASW, and 31°C *E. acroporae*. Median genes per cell were 732, 881, 748, and 715, respectively, while median unique molecular identifiers (UMIs) per cell were 1,738, 2,104, 1,699, and 1,839. Sequencing saturation ranged from 72% to 78.5%.

CellBender (v0.3.2)^135^ was used to mitigate background noise in the count matrices using parameters: - expected-cells 10,000, -total-droplets-included 30,000, -fpr 0.01, and -epochs 150. Additional parameters adjusted according to the ELBO learning curve included -learning-rate (2e-5 to 1e-5) and -low-count-threshold (set at 100 or disabled). Noise-removed matrices were imported into Seurat (v5.1.0)^136^ using scCustomize (v3.0.1). Cells with low (< 100 UMIs) or very high (> 8,000 UMIs) molecule counts were removed. Potential doublets were recognized and removed using scDblFinder (v1.16.0).^137^ After filtering, an average of 7,409 cells remained per sample, with a median of 684 genes and 1,488 UMIs per cell, comparable to previous stony coral single-cell studies.^52^

#### Cell clustering and cell type annotation

After filtering, the four Seurat objects were merged, normalized with the NormalizeData function, and subset to the top 2,000 most variable features using the FindVariableFeatures function. Data were scaled, and PCA dimensional reduction was performed. Dimensionality for clustering analysis was determined with an elbow plot, selecting the first 30 principal components. Clusters were identified using FindNeighbors and

FindClusters with a resolution of 0.6. Clusters were visualized via UMAP (DimPlot function). Marker genes defining clusters were identified with FindAllMarkers (parameters: min.diff.pct = 0.1, only.pos = TRUE). Violin plots (VlnPlot) and heatmaps (ComplexHeatmap) were generated to visualize marker gene expression. Differences in cluster proportions between treatments were tested and visualized using scProportionTest.^138^

To annotate cluster identity, 491 highly variable genes from a published single-cell data set of *S. pistillata* (clade 4)^52^ were aligned against single-cell markers of *S. pistillata* (clade 1) using blastn (v2.16.0).^139^ Reciprocal best hits (RBH) identified orthologous genes with thresholds: percent identity ≥ 85%, e-value ≤ 1e-5 (bidirectional), and query coverage ≥ 50% (at least one direction; Table S4). A total of 280 orthologous marker genes were identified, and 181 highly variable markers (average log2 fold-change > 1.7) were used to classify cell clusters. Visualization of results was performed using ggbump package.

#### Gene age estimation

For each gene in the *S. pistillata* clade 1 genome in BRAKER-derived gene models, a phylostratigraphic age^54,179^ was estimated using GenEra (v1.2.0).^140^ GenEra uses DIAMOND^129^ searches of the NR database to assign genes to one of 12 categorical age classes (or phylostrata), from youngest (rank 12, *S. pistillata*-specific) to oldest (rank 1, found in all cellular organisms). These numerical ranks were used to calculate the average age of marker genes for each single cell cluster. The log2 fold change in the proportion of marker genes of each age versus the proportion of all genes of each age was calculated as described previously.^52^ Data were presented as heatmaps plotted in R with ComplexHeatmap (v2.18.0).^122^ Benjamini-Hochberg-corrected Fisher’s exact tests were applied to test whether the set of marker genes for each cluster is enriched or depleted for genes from each phylostratum.

#### Differential expression and functional analysis

Differential expression analysis was performed at the single-cell level using Seurat (v5.1.0)^136^ FindMarkers function with the MAST algorithm. Differential expression was tested between the control (28°C FASW) and each of the other three treatments in the same cell type. Genes with an adjusted *p* < 0.05 were considered significantly differentially expressed. GO enrichment analysis was performed as described above. Significantly enriched GO terms related to coral heat stress responses were visualized using ggplot2. Intersection analysis of significantly expressed genes identified in bulk and single-cell RNA-seq data was visualized using UpSetR^180^ package. Expression levels of key genes were visualized on UMAP plots using the Seurat (v5.1.0)^136^ FeaturePlot function.

### Statistical analysis

All statistical analyses were performed in R (v4.3.2) using RStudio (v2023.12.1+402). Specific statistical tests used are indicated in each method section or figure legend. Data normality and homogeneity of variance were assessed using Shapiro-Wilk and Levene’s tests, respectively. Sample sizes or gene numbers for each analysis are indicated by “n” in corresponding figure panels or legends.

