## Supplemental Figures S1-S13 for "Microbial profiling and single-cell transcriptomics reveal probiotic mechanisms of coral thermal resilience"

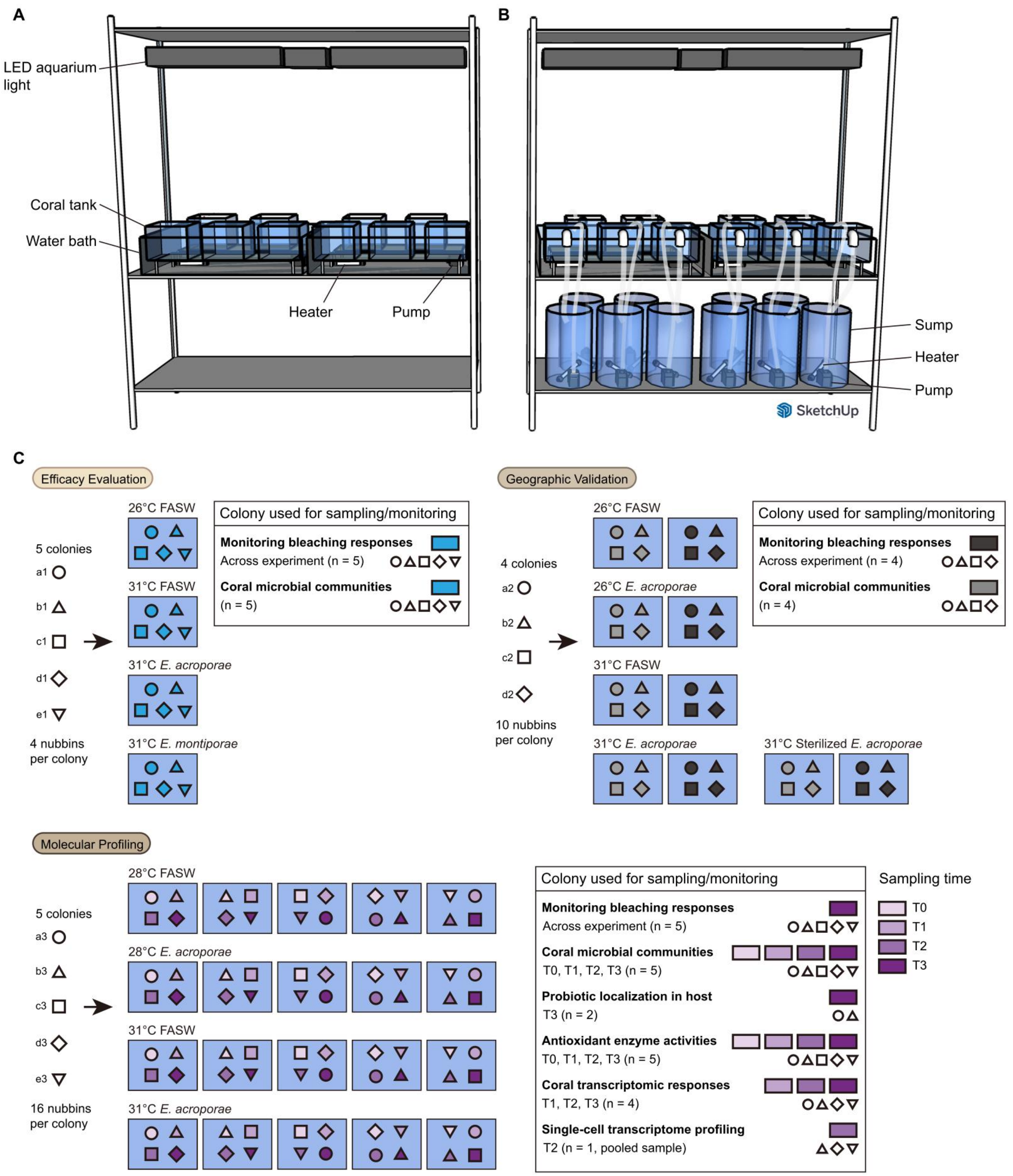


**Figure S1. Schematic overview of experimental tanks and replication design for each experiment, related to Figures 1 and S2**

(A) Schematic of the experimental tank used in the efficacy evaluation experiment.

(B) Schematic of the experimental tank used in the geographic validation and molecular profiling experiments.

(C) Replication design for the three coral bleaching experiments. Different coral colonies are represented by distinct shapes, with individual colony codes labeled on the left. The shape and fill color indicate which colonies were included in different assays.

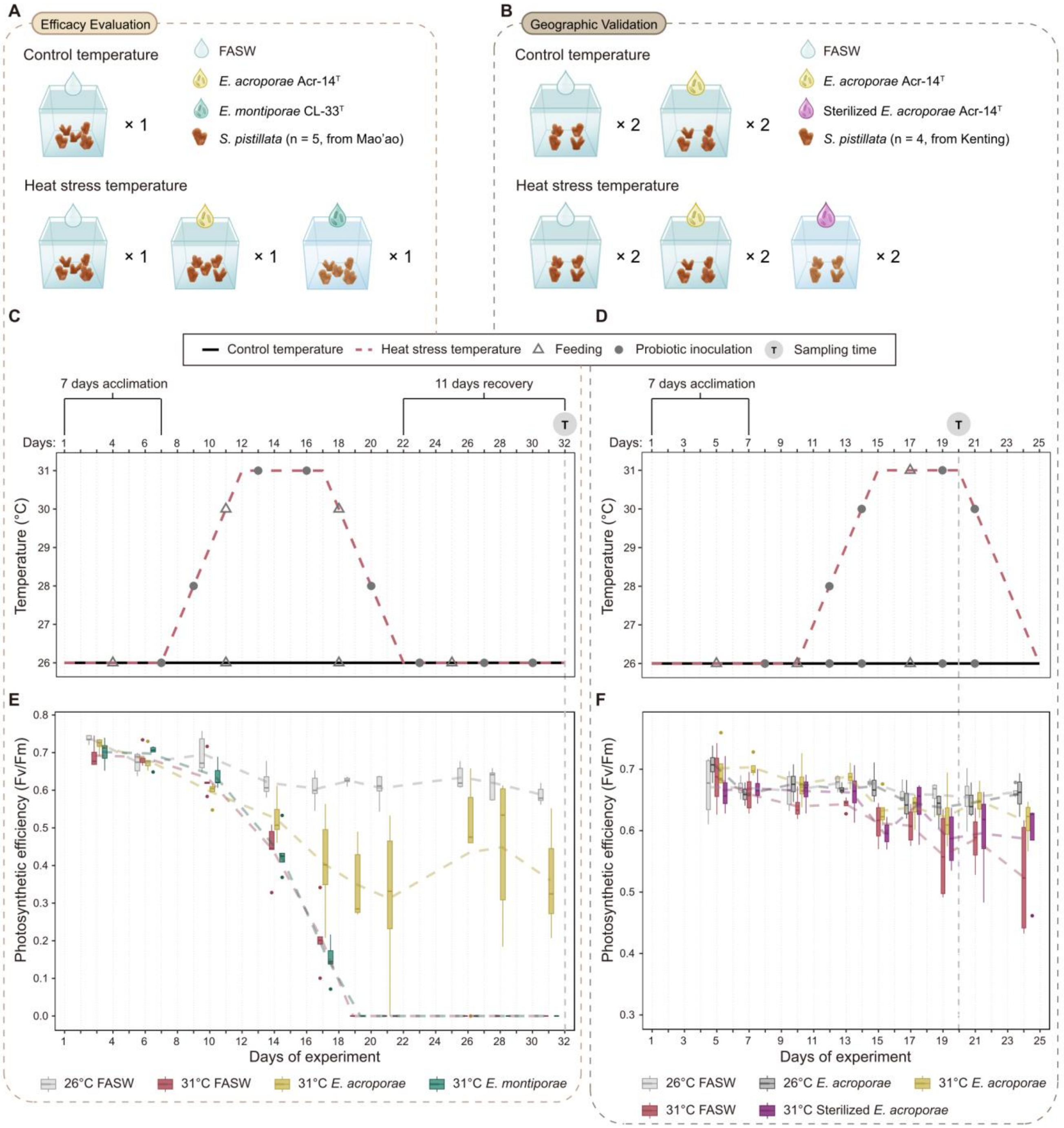


**Figure S2. Study design and coral bleaching responses of efficacy evaluation and geographic validation, related to Figures 1 and S1**

(A) Treatment and replication design of efficacy evaluation. Each treatment consisted of one aquarium containing five coral nubbins, each originating from a different colony (n = 5).

(B) Treatment and replication design of geographic validation. Each treatment consisted of two aquarium systems: one for sampling and another for measuring coral photosynthetic efficiency. Each aquarium contained four coral nubbins, each from a different colony (n = 4).

(C and D) Detailed treatment regimens for efficacy evaluation (C) and geographic validation (D), including temperature settings and dates for feeding, probiotic inoculation, and sample collection.

(E and F) Photosynthetic efficiency (Fv/Fm) of nubbins in efficacy evaluation (E) and geographic validation (F). In boxplots, dots represent outliers, and dashed lines connecting treatment means ($n = 5$ in efficacy evaluation; $n = 4$ in geographic validation; One-way ANOVA with Tukey post-hoc test).

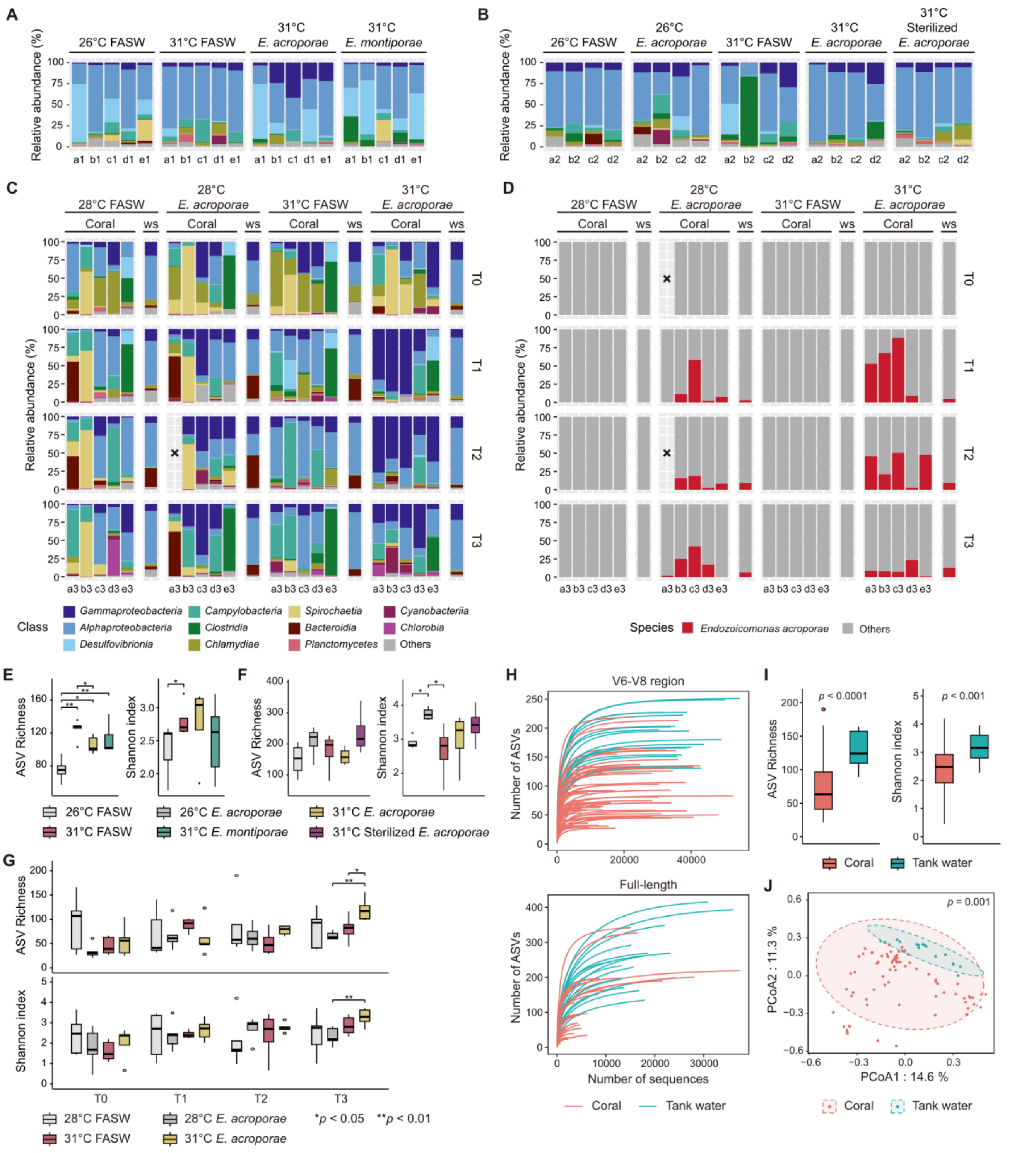


**Figure S3. Effects of treatments on coral microbial composition and diversity, related to Figures 2 and 3 and STAR Methods**

(A-C) Relative abundance of bacterial class-level composition across treatments in efficacy evaluation (A), geographic validation (B), and molecular profiling (C), determined by short-length 16S rRNA gene sequencing. Tank water samples (ws); lost or failed to sequence samples (x).

(D) Relative abundance of *E. acroporae* in samples, determined by full-length 16S rRNA gene sequencing. All ASVs belonging to genus *Endozoicomonas* were assigned to *E. acroporae*.

(E-G) ASV richness and Shannon index across treatments in efficacy evaluation (E), geographic validation (F), and molecular profiling (G) (efficacy evaluation: n = 5, geographic validation: n = 4, molecular profiling: n = 5; Wilcoxon signed-rank tests; *$p < 0.05$; **$p < 0.01$).

(H) Rarefaction curves for all samples in molecular profiling, with data derived from short-length or full-length sequencing methods.

(I) ASV richness and Shannon index comparing coral (n = 78) and tank water (n = 16) samples in molecular profiling (Wilcoxon signed-rank tests).

(J) Principal coordinates analysis (PCoA) illustrating microbial community differences between coral (n = 78) and tank water (n = 16) samples in molecular profiling (PERMANOVA).

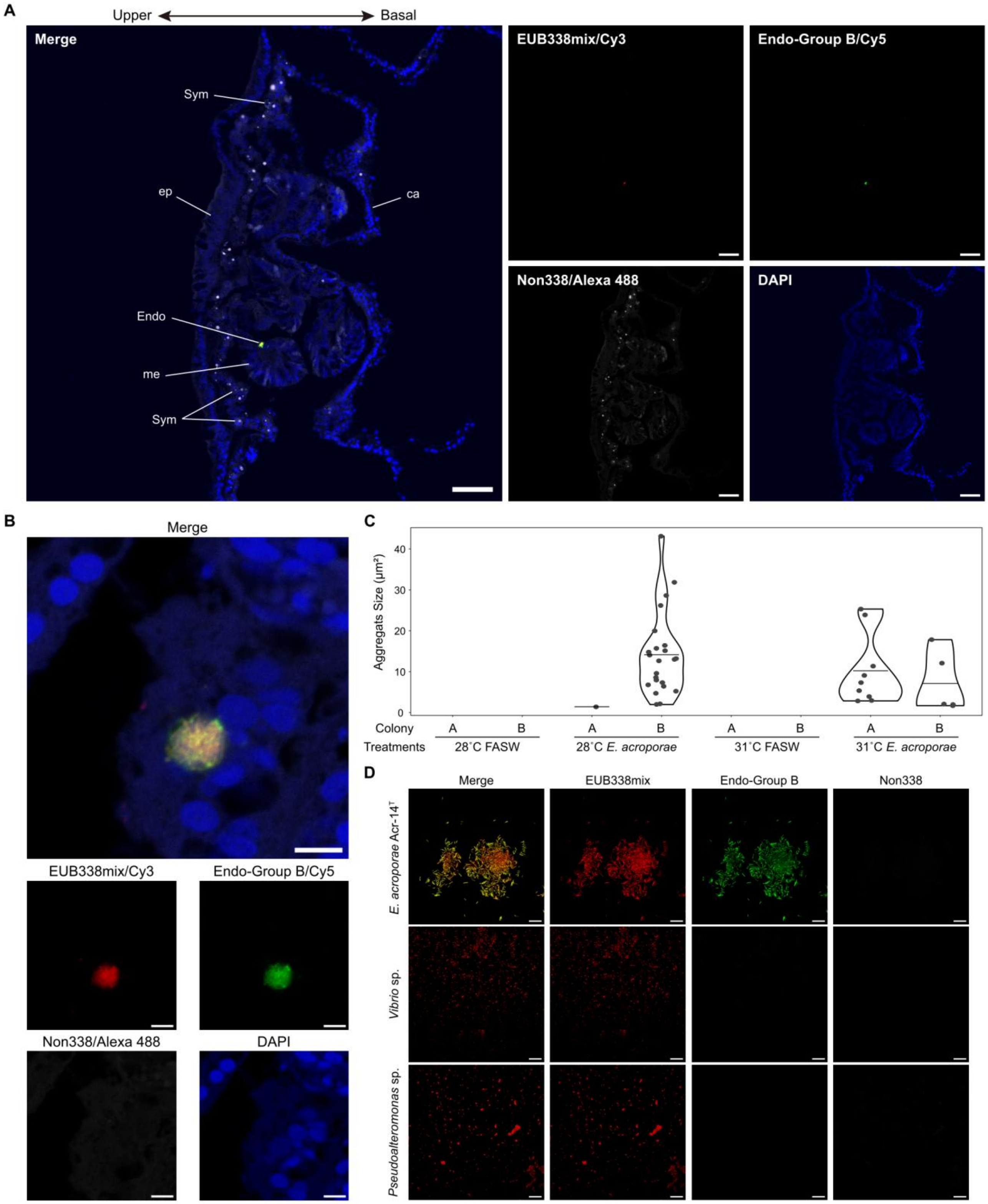


**Figure S4. FISH detection of *Endozoicomonas* in coral tissues and bacterial cultures, related to Figure 2 and STAR methods**

(A) Individual and merged channels showing a tissue section from the 31°C *E. acroporae* treatment in molecular profiling, with *Endozoicomonas* signals visible. Symbiodiniaceae exhibited strong autofluorescence under Alexa 488. ca: calicoblastic cell layer; Endo: *Endozoicomonas*; ep: epidermal layer; me: mesenterial filaments; Sym: Symbiodiniaceae. Scale bars: 50 µm.

(B) Individual and merged channels showing *Endozoicomonas* signals at higher magnification in a sample from the 28°C *E. acroporae* treatment in molecular profiling. Scale bars: 5 µm.

(C) Size of *Endozoicomonas* aggregations observed in microscope slides across the four treatments in molecular profiling (two samples per treatment, five slides per sample, 40 slides in total).

(D) Validation of FISH probe specificity using the universal prokaryotic probe EUB338mix and the Endo-Group B probe on three bacterial cultures: *E. acroporae* Acr-14$^T$, *Vibrio coralliilyticus* YB strain, and *Pseudoalteromonas piratica.* Scale bars: 10 µm.

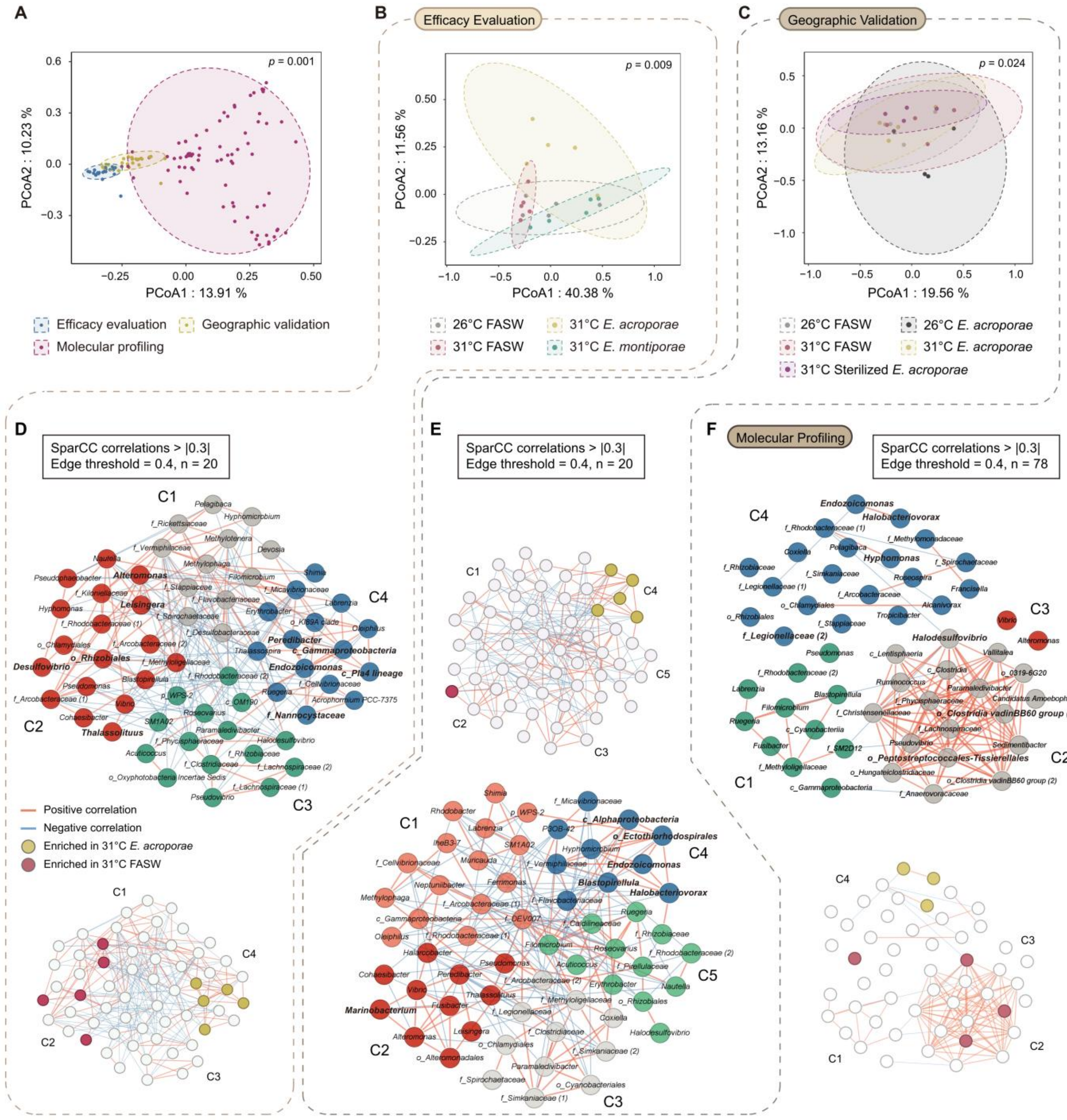


**Figure S5. Multivariate and network analyses of coral microbial community, related to Figure 3**

(A-C) Principal coordinates analysis (PCoA) illustrating the effects of experimental batches (A) or treatments in efficacy evaluation (B) and geographic validation (C) on coral microbial composition. 95% confidence ellipses were drawn around the centroid of each group (PERMANOVA; pairwise comparison *p* values adjusted using Bonferroni correction).

(D-F) SparCC correlation networks of the 60 most prevalent bacterial genera in efficacy evaluation (D), geographic validation (E), and molecular profiling (F). Small panels show networks overlaid with genera enriched specifically in the 31°C FASW or 31°C *E. acroporae* treatments. Clusters identified in these networks are labeled with “C” followed by a number.

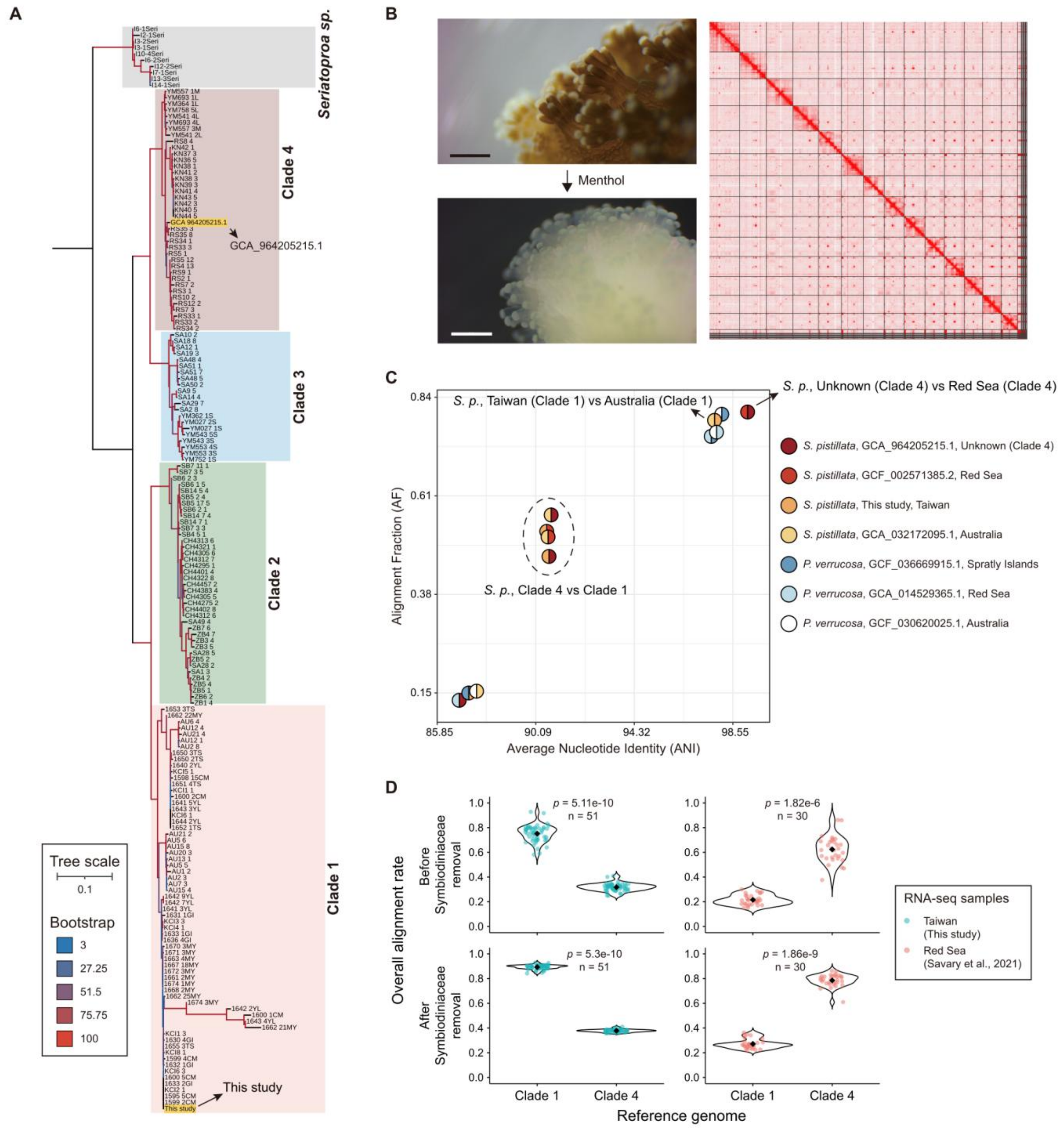


**Figure S6. Genetic divergence between *S. pistillata* clades 1 and 4, related to STAR methods**

(A) Phylogenetic tree of *S. pistillata* ITS rDNA sequences obtained from this study, a released genome (GCA_964205215.1), and a previous study. The tree is rooted using an outgroup (*Seriatopora* sp.).

(B) Hi-C contact map of *S. pistillata* clade 1 and menthol-induced bleaching treatment. Scale bars: 1 mm.

(C) Pairwise whole-genome alignment illustrating genetic divergence between samples. Colors in pie chart indicate genomes used for alignment.

(D) Overall alignment rate of RNA sequences mapped to clade 1 or clade 4 *S. pistillata* genomes. Each dot represents one sample, and black diamonds indicate group means (Wilcoxon signed-rank tests).

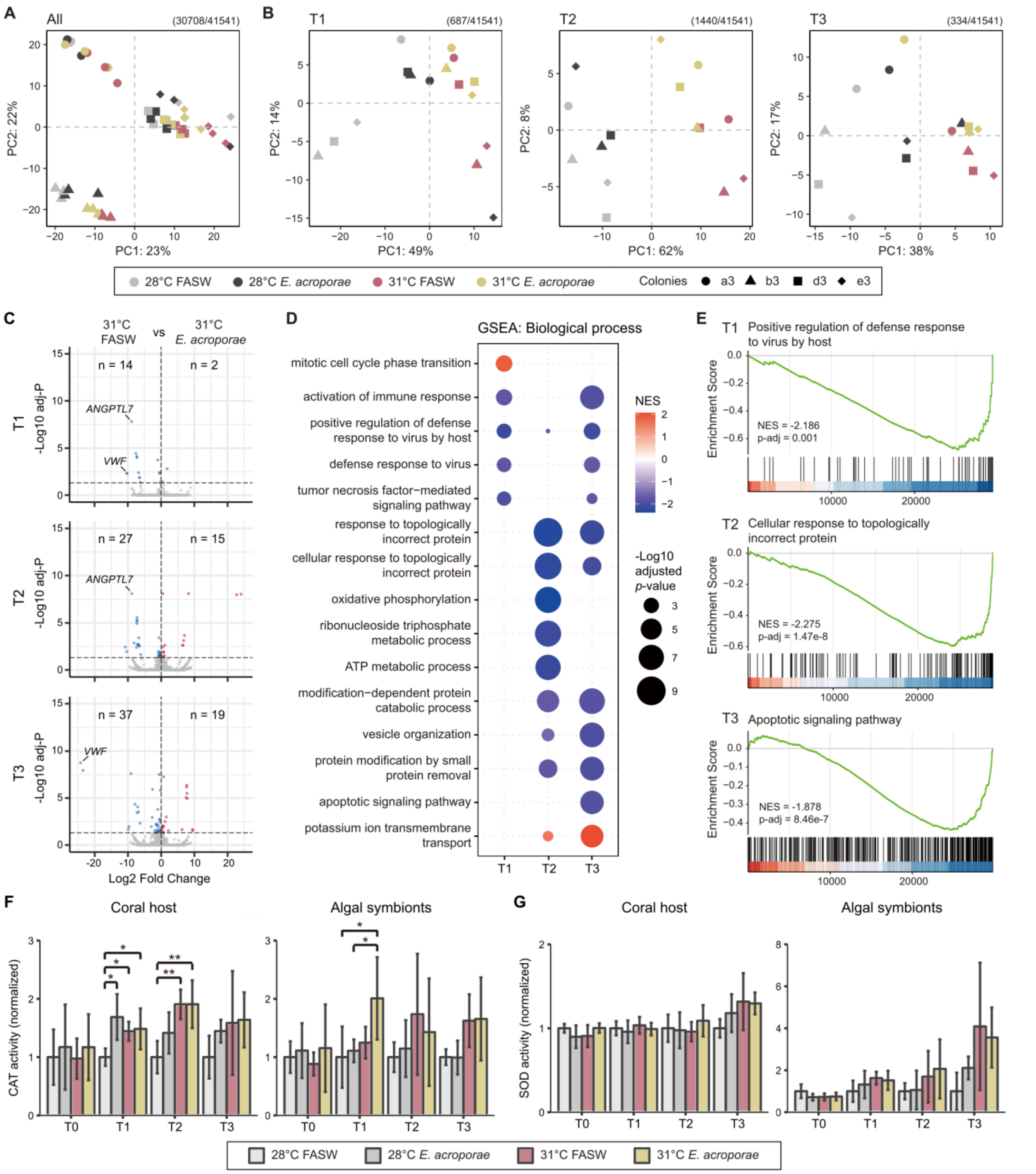


**Figure S7. Effect of heat stress and probiotic treatments on the coral host transcriptome, related to Figure 4**

(A and B) PCA plots illustrating coral host transcriptomic profiles, constructed using all pre-filtered genes (A) or all DEGs identified at each sampling time (B). Numbers in parentheses indicate numbers of genes used for PCA construction and total gene count in the genome-guided (*S. pistillata* clade 1) transcriptome.

(C) Volcano plots showing results of differential expression analyses between the 31°C FASW and 31°C *E. acroporae* treatments. Numbers of upregulated and downregulated DEGs are indicated by n.

(D) GSEA of genes ranked by Wald statistic from DESeq2, comparing the 31°C FASW and 31°C *E. acroporae* treatments. The five most significantly enriched GO terms in each biological process category are showed (adjusted $p < 0.05$).

(E) GSEA running scores for significant GO terms identified at T1, T2, and T3.

(F and G) Catalase (F) and superoxide dismutase (G) activities in coral tissue and algal symbionts across the four treatments and sampling times. Activities were normalized to the control treatment (28°C FASW) at each sampling time (*adjusted $p < 0.05$, **adjusted $p < 0.01$; n = 5, One-way ANOVA with Tukey post-hoc test).

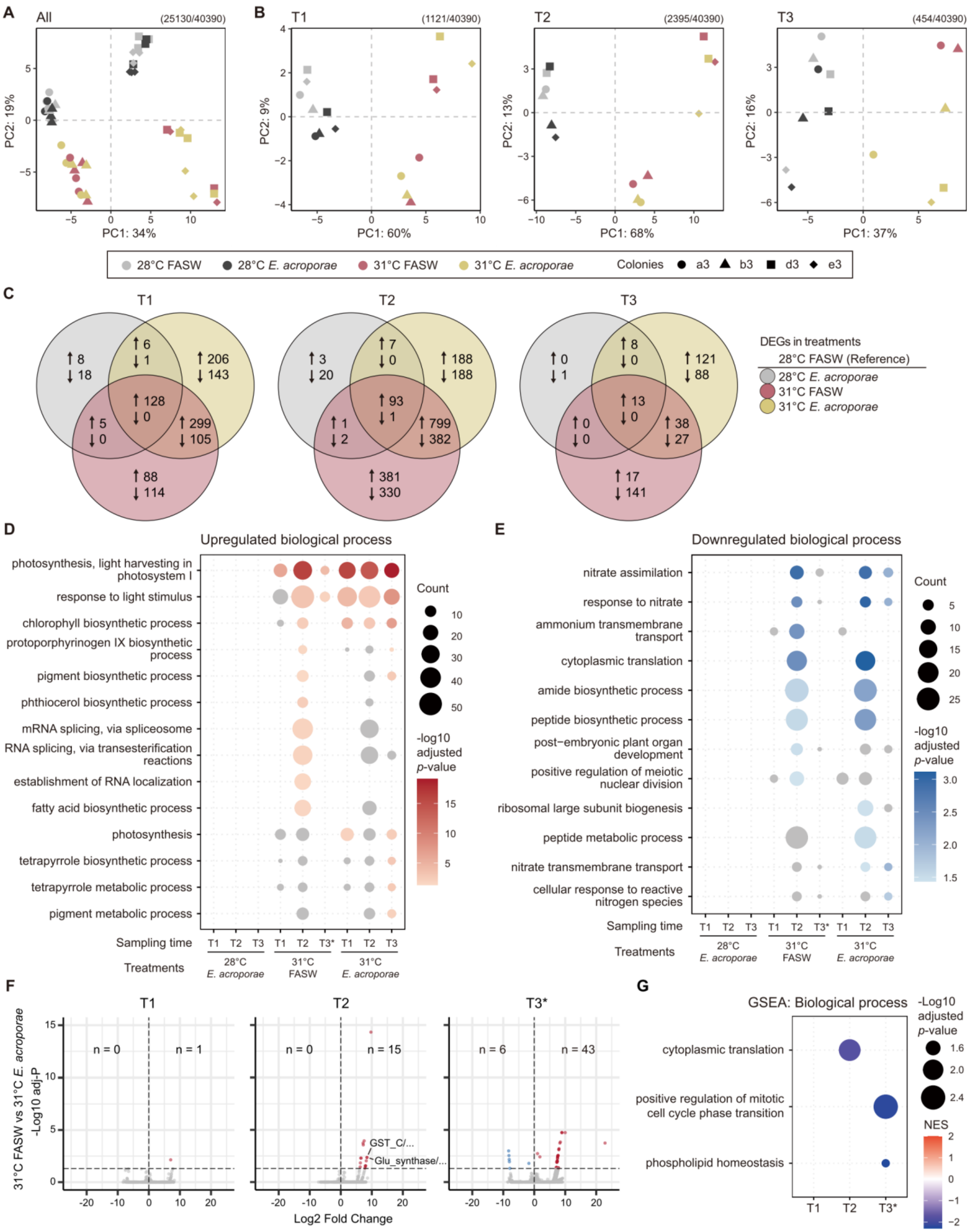
A
All
(25130/40390)
PC1: 34%
PC2: 19%
B
T1
(1121/40390)
PC1: 60%
PC2: 9%
T2
(2395/40390)
PC1: 68%
PC2: 13%
T3
(454/40390)
PC1: 37%
PC2: 16%
28°C FASW
28°C E. acroporae
31°C FASW
31°C E. acroporae
Colonies
a3
b3
d3
e3
C
T1
↑8 ↓18
↑6 ↓1
↑206 ↓143
↑128 ↓0
↑5 ↓0
↑299 ↓105
↑88 ↓114
T2
↑3 ↓20
↑7 ↓0
↑188 ↓188
↑93 ↓1
↑1 ↓2
↑799 ↓382
↑381 ↓330
T3
↑0 ↓1
↑8 ↓0
↑121 ↓88
↑13 ↓0
↑0 ↓0
↑38 ↓27
↑17 ↓141
DEGs in treatments
28°C FASW (Reference)
28°C E. acroporae
31°C FASW
31°C E. acroporae
D
Upregulated biological process
photosynthesis, light harvesting in photosystem I
response to light stimulus
chlorophyll biosynthetic process
protoporphyrinogen IX biosynthetic process
pigment biosynthetic process
phthiocerol biosynthetic process
mRNA splicing, via spliceosome
RNA splicing, via transesterification reactions
establishment of RNA localization
fatty acid biosynthetic process
photosynthesis
tetrapyrrole biosynthetic process
tetrapyrrole metabolic process
pigment metabolic process
Count
-log10 adjusted p-value
Sampling time
T1 T2 T3 T1 T2 T3* T1 T2 T3
Treatments
28°C E. acroporae
31°C FASW
31°C E. acroporae
E
Downregulated biological process
nitrate assimilation
response to nitrate
ammonium transmembrane transport
cytoplasmic translation
amide biosynthetic process
peptide biosynthetic process
post−embryonic plant organ development
positive regulation of meiotic nuclear division
ribosomal large subunit biogenesis
peptide metabolic process
nitrate transmembrane transport
cellular response to reactive nitrogen species
Count
-log10 adjusted p-value
Sampling time
T1 T2 T3 T1 T2 T3* T1 T2 T3
Treatments
28°C E. acroporae
31°C FASW
31°C E. acroporae
F
T1
n = 0
n = 1
T2
n = 0
n = 15
GST_C/...
Glu_synthase/...
T3*
n = 6
n = 43
31°C FASW vs 31°C E. acroporae
-Log10 adj-P
Log2 Fold Change
G
GSEA: Biological process
cytoplasmic translation
positive regulation of mitotic cell cycle phase transition
phospholipid homeostasis
-Log10 adjusted p-value
NES
T1 T2 T3*

**Figure S8. Effect of heat stress and probiotic treatments on the algal symbiont transcriptome, related to Figure 4**

(A and B) PCA plots showing algal symbionts transcriptomic profiles. PCA was conducted using all pre-filtered genes (A) or all DEGs identified at each time point (B). Numbers in parentheses at the top indicate genes used for PCA construction and total gene count from the *C. goreaui* genome.

(C) Numbers of DEGs identified in algal symbionts from each treatment at each sampling time. Arrows indicate upregulated and downregulated DEGs (adjusted $p < 0.05$).

(D and E) GO enrichment analyses of upregulated (D) or downregulated (E) DEGs identified in algal symbionts across treatments and sampling time points. Bubble size reflects DEG count. Grey bubbles indicate GO term with $p < 0.05$ but adjusted $p > 0.05$ (Benjamini-Hochberg correction).

(F) Volcano plots showing differential expression analysis results between the 31°C FASW and 31°C *E. acroporae* treatments. Number of upregulated and downregulated DEGs are indicated by n.

(G) Gene set enrichment analysis (GSEA) of genes ranked by Wald statistic from DESeq2 results, comparing the 31°C FASW and 31°C *E. acroporae* treatments. Significantly enriched GO terms (adjusted $p < 0.05$) within the biological process category are showed.

*Note: Due to severe bleaching, DEGs from the 31°C FASW treatment at T3 were derived from 2 algal symbiont samples (colonies a3 and b3, n = 2, STAR Methods).

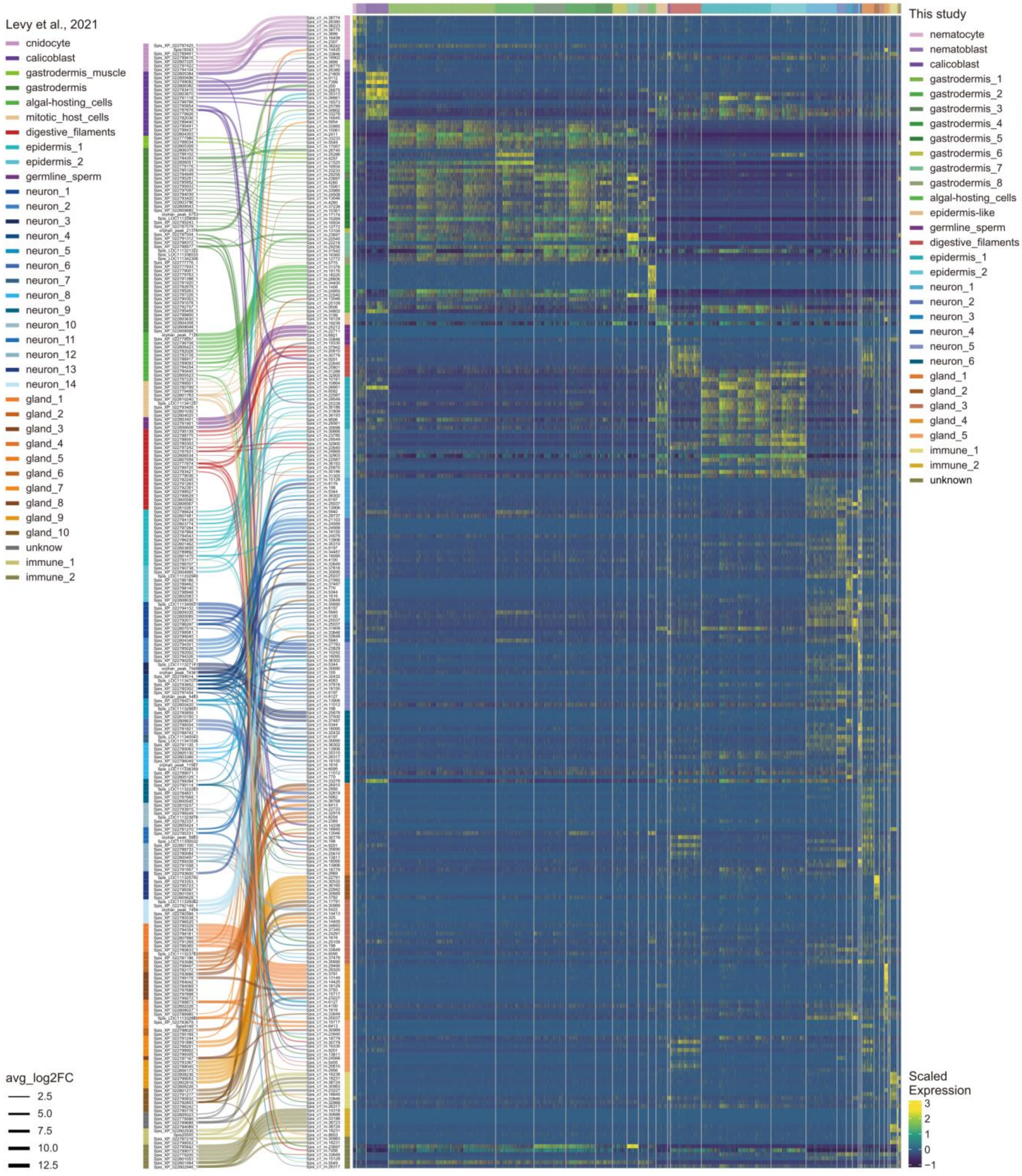


**Figure S9. Cell-type classification based on orthologous marker genes identified between *S. pistillata* clades 1 and 4 cell markers, related to Figure 5**

Bump plot illustrating orthologous marker genes defined between *S. pistillata* clades 4 and 1 cell markers. The left column lists 280 markers belong to *S. pistillata* clade 4 (see Table S4). Orthologous markers are connected only to highly variable markers identified in *S. pistillata* clade 1 (average log2 fold change > 1.7; right column, 181 genes in total). Line thickness indicates average log2 fold change value. Note that marker genes identified in *S. pistillata* clade 1 may not be exclusive to a single cell cluster. Heatmap shows normalized expression of orthologous marker genes across cells.

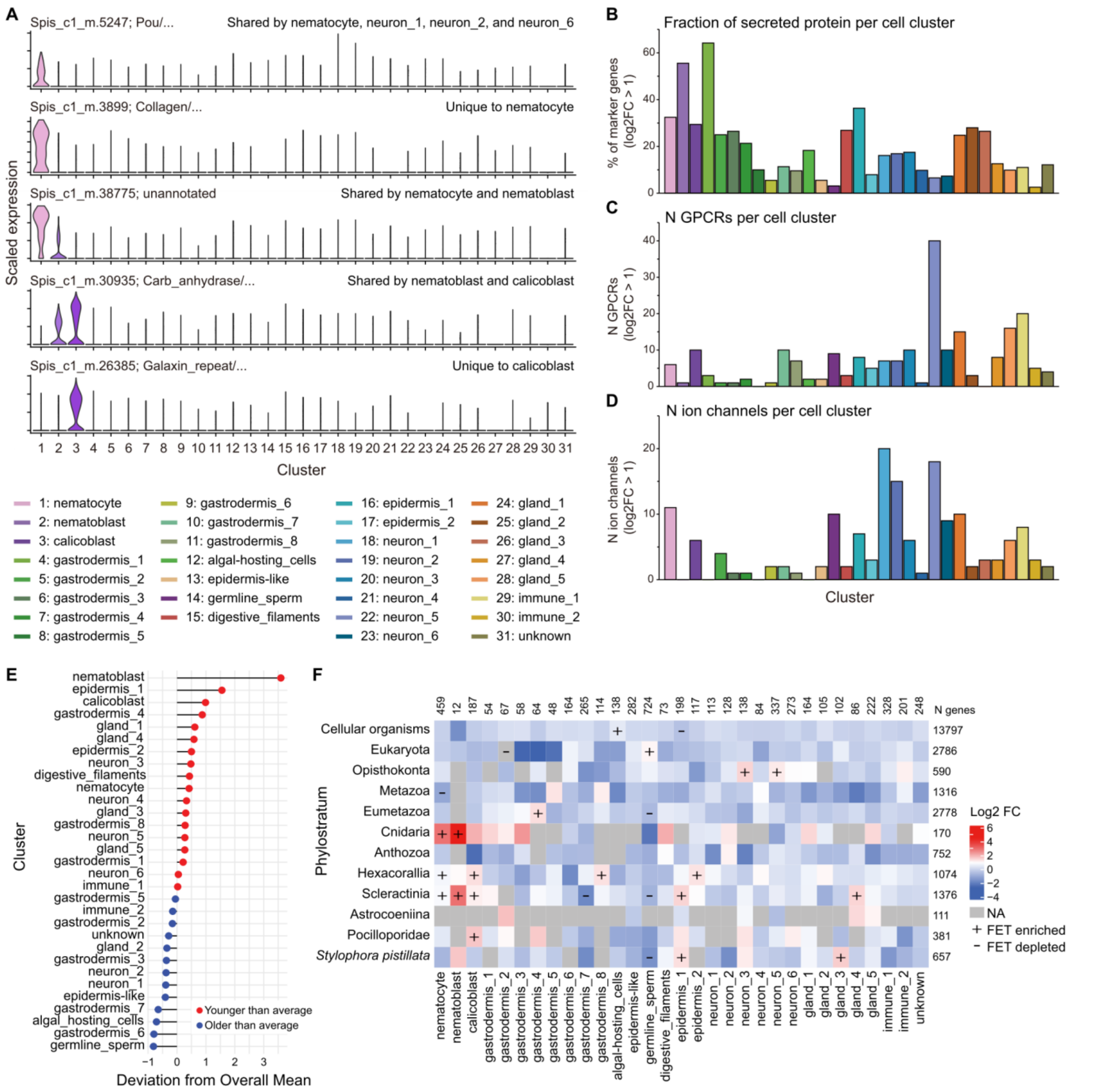

**Figure S10. *S. pistillata* clade 1 cell atlas marker gene functions and phylostratigraphy, related to Figure 5**

(A) Expression profiles illustrating that nematoblasts share multiple marker genes with nematocytes and one marker gene with calicoblasts.

(B) Percentage of predicted secreted proteins (containing a signal peptide and no transmembrane domains) among marker genes (with average log2 fold change > 1) for each cell cluster.

(C) Total number of GPCRs genes (containing Pfam domains 7tm_1, 7tm_2, or 7tm_3) identified among marker genes (with average log2 fold change > 1) for each cell cluster.

(D) Total number of ion channels genes (based on Pfam domains, see Table S5) identified among marker genes (with average log2 fold change > 1) for each cell cluster.

(E) Mean age of marker genes (with average log2 fold change > 1.7) for each cell cluster, as defined by phylostratigraphy.

(F) Enrichment or depletion of marker genes within each cell cluster among phylostrata. Statistical significance determined by Fisher exact test (adjusted $p < 0.1$, Benjamini-Hochberg correction).

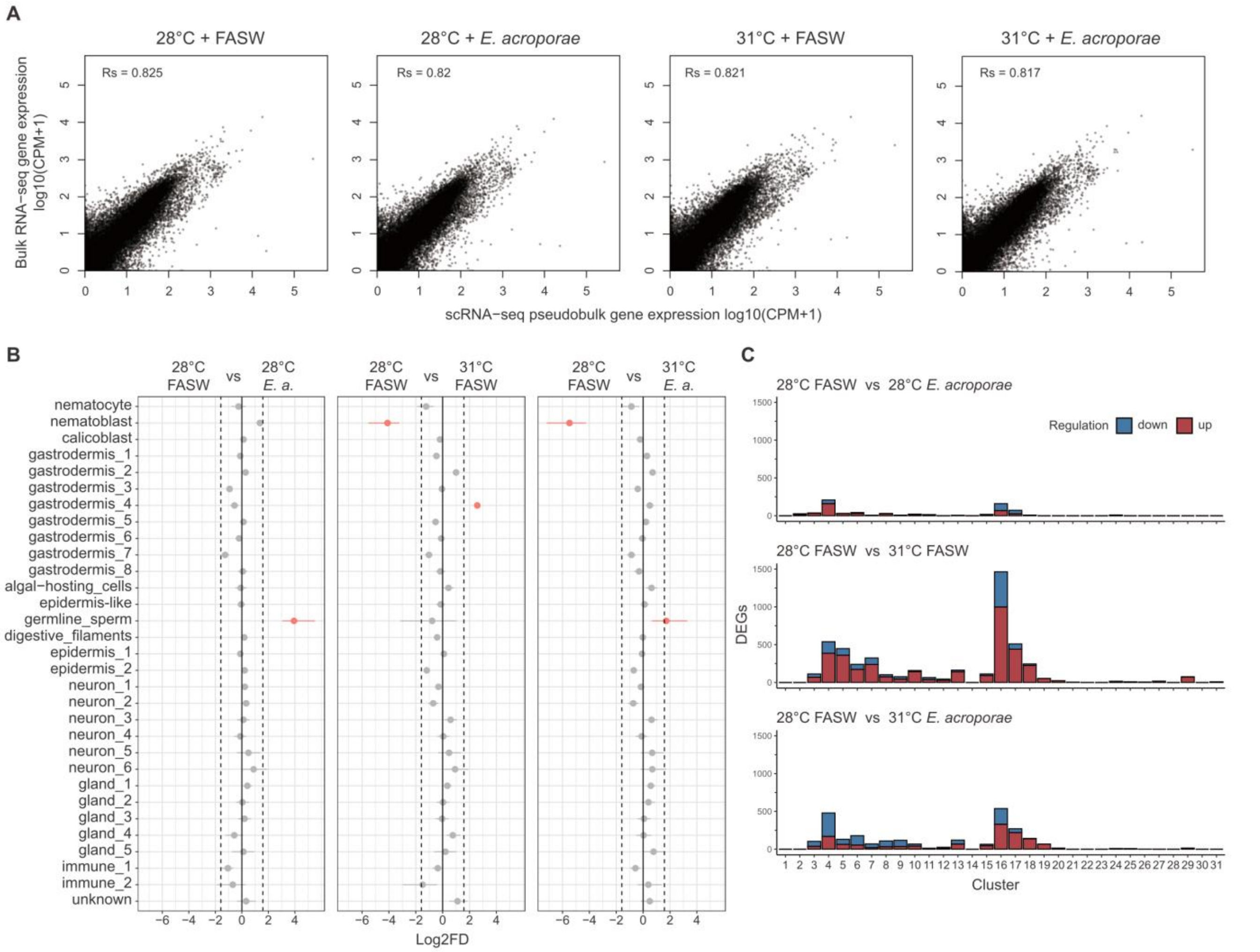


**Figure S11. Concordance between bulk and single-cell transcriptomes and treatment effects on coral cell clusters, related to Figures 5 and 6**

(A) Gene-wise comparison of pseudobulk scRNA-seq and bulk RNA-seq expression levels, with the Spearman correlation coefficient indicated.

(B) Relative differences in cell cluster proportions between control and other treatments. Red indicates significant changes (FDR < 0.01 and |Log2FC| > 1.6, permutation test).

(C) Numbers of DEGs (adjusted $p < 0.05$) identified in each cell cluster.

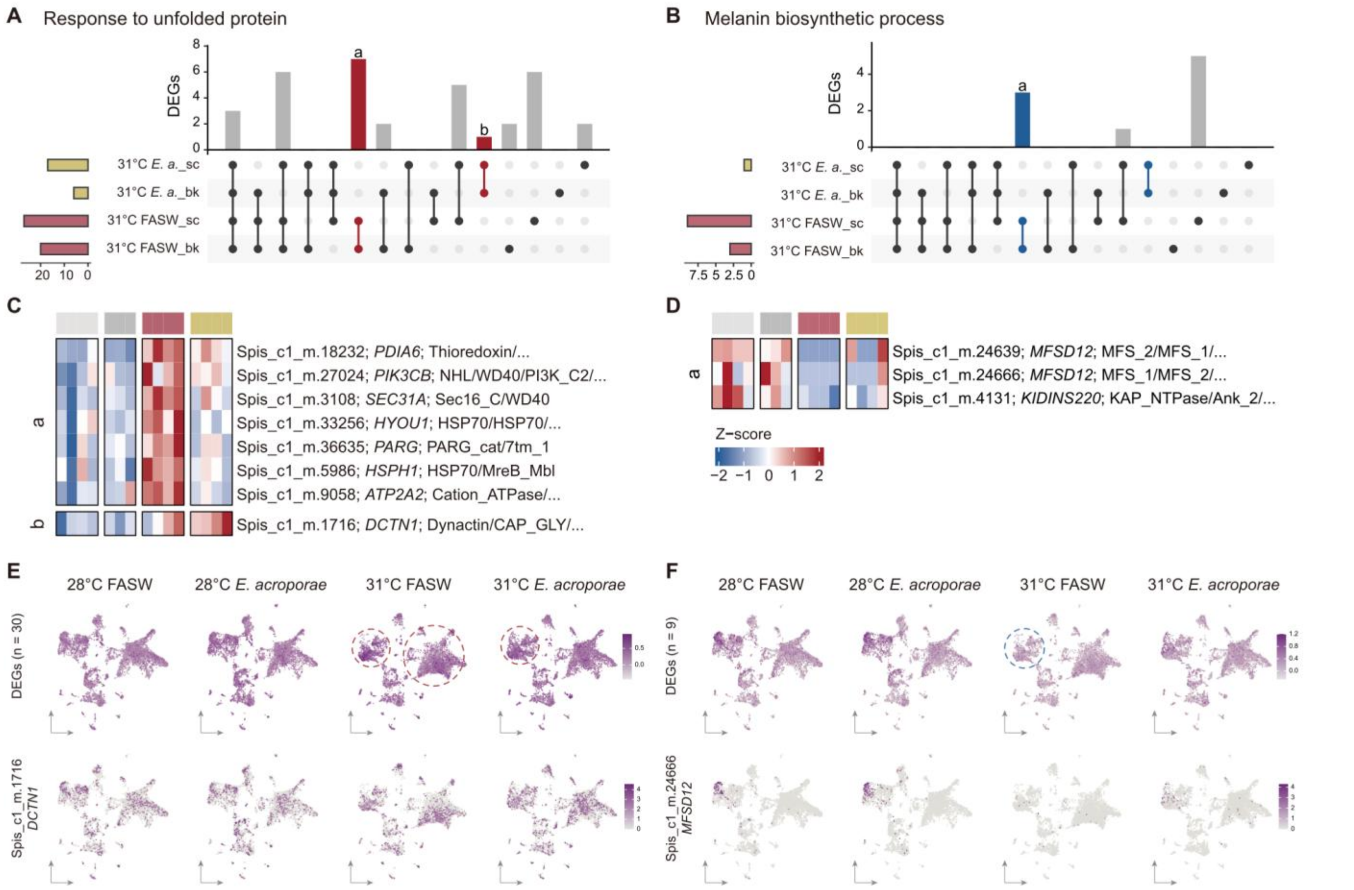


**Figure S12. Additional gene expression profiles potentially contributing to the probiotic effect, related to Figure 7**

(A and B) Overlaps of upregulated (red) and downregulated (blue) DEGs in the 31°C FASW and 31°C *E. acroporae* treatments (both compared to controls), associated with GO terms for response to unfolded protein (A) and melanin biosynthetic process (B). DEGs identified by bulk (“bk”) or single-cell (“sc”) transcriptomes are marked. *Bar chart letters correspond to genes in panel (C) or (D), with colors highlighting DEGs* uniquely detected in either the 31°C FASW or 31°C *E. acroporae* treatments.

(C and D) Bulk RNA-seq expression of selected genes associated with the response to unfolded protein (C) and melanin biosynthetic process (D). Annotated gene name and Pfam domains are listed.

(E and F) Single-cell RNA-seq normalized expression of DEGs for response to unfolded protein (E) or melanin biosynthetic process (F). Upper panels show all corresponding DEGs, with enriched cell clusters highlighted by dashed circles.

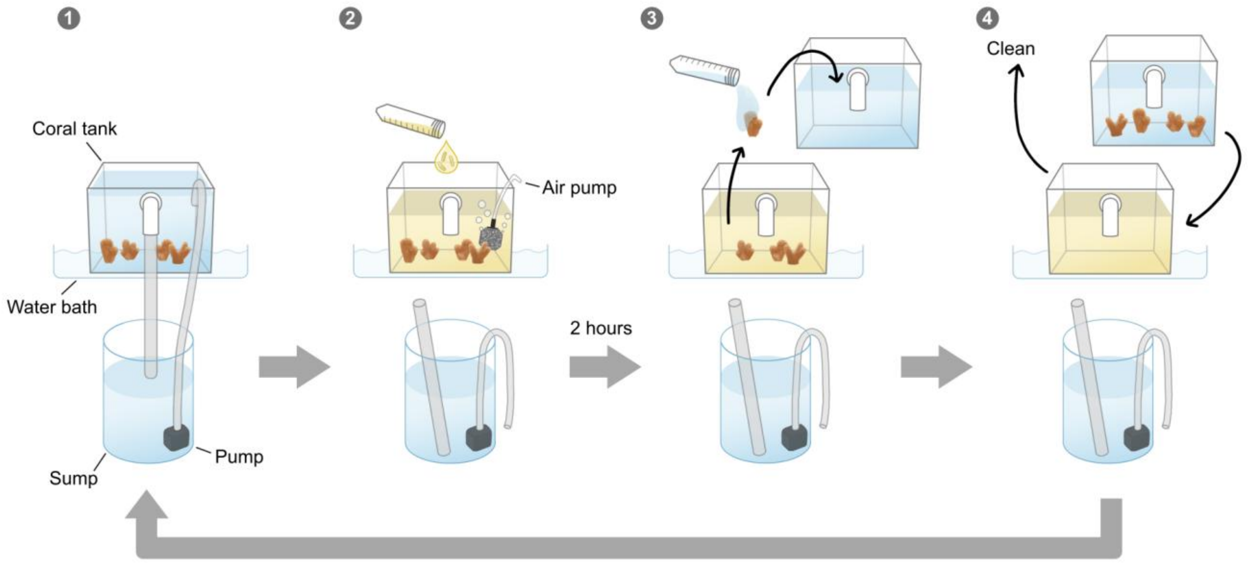


**Figure S13. Detailed probiotic inoculation and feeding procedures, related to STAR methods**

Step 1: Water circulation between the coral tank and its sump was halted by turning off the submersible pump and disconnecting silicon tubing. Step 2: Concentrated probiotic solutions or placebo (FASW) were added to the coral tank, with water flow maintained by an air pump during inoculation. Step 3: After two hours of inoculation, coral nubbins were rinsed with FASW and transferred to a new coral tank prefilled with temperature-adjusted fresh ASW. Step 4: Once all coral nubbins were transferred, the used coral tank was replaced and cleaned. Finally, all silicon tubing was reconnected, and the submersible pump was reactivated to resume water circulation with the sump. In efficacy evaluation, as sumps were not used, procedures involving the sump were not required. For feeding, following the same procedure, replacing the probiotic solution with concentrated Artemia.
